# Warfarin analogs target disulfide bond-forming enzymes and suggest a residue important for quinone and coumarin binding

**DOI:** 10.1101/2024.02.18.580799

**Authors:** Dariana Chavez, Gwendolyn Nita Amarquaye, Adrian Mejia-Santana, Dyotima, Kayley Ryan, Lifan Zeng, Cristina Landeta

## Abstract

Disulfide bond formation has a central role in protein folding of both eukaryotes and prokaryotes. DsbB and VKOR enzymes catalyze the oxidation of the oxidoreductase partner and the formation of *de novo* disulfide bonds using quinone as cofactor. We have used *E. coli* and a family of warfarin analogs to study *de novo* disulfide bond formation. We found that human VKORc1 can function in *E. coli* by removing two positive residues, allowing the search for novel anticoagulants. One analog was capable of inhibiting both bacterial DsbB and VKOR, and a second one antagonized only the mammalian enzymes. We identified the two amino acid residues responsible for binding. One of these is also essential for quinone binding in both DsbB and VKOR. Our studies highlight a conserved role of this residue in *de novo* disulfide-generating enzymes and enable the design of novel anticoagulants or antibacterials using coumarin as a scaffold.

## Introduction

Disulfide bonds (DSBs) are covalent linkages between the sulfur atoms of two cysteine residues. DSB formation is a fundamental aspect of protein folding in both eukaryotes and prokaryotes. In bacteria, DSBs play critical roles in the folding and stability of proteins involved in important cellular processes, including cell division, outer membrane biogenesis, virulence, and antibiotic resistance^1–4^. Hence, DSB-forming enzymes represent a new drug target simultaneously affecting several proteins localized in the cell envelope.

In *Escherichia coli*, DSB formation occurs in the periplasm and is catalyzed by DsbA and DsbB enzymes, which work together to introduce DSBs into many proteins through their catalytic cysteines. DsbA is a periplasmic protein, a member of the thioredoxin family, that catalyzes the formation of DSBs into substrate proteins through its Cys-X-X-Cys (CXXC) active site^5^. DsbB is a membrane protein that regenerates DsbA’s activity by transferring the electrons to quinones. First, DsbB interacts with DsbA through its CX_n_C motif forming a covalent complex, then it uses a separate CXXC motif to form *de novo* a DSB by reducing a quinone molecule^6–9^. Aerobically, DsbB transfers electrons to ubiquinone, while anaerobically to menaquinone (aka Vitamin K2)^8,10,11^.

Most gram-negative bacteria use DsbA-DsbB to introduce DSBs however, actinobacteria, cyanobacteria, and δ-proteobacteria, use an alternative enzyme named VKOR (for <u>v</u>itamin <u>K</u> ep<u>o</u>xide <u>r</u>eductase) that performs the analogous function of DsbB^12^. DsbB and VKOR share no protein sequence identity, but they exhibit similar structural features and contain a quinone cofactor to generate a DSB *de novo*^13,14^. In fact, the interactions between the redox-active cysteines of DsbA and VKOR proceed in the same steps seen between DsbA and DsbB^14,15^. While DsbB has no homolog in eukaryotes, bacterial VKOR has eukaryotic homologs in plants, arthropods, vertebrates, and humans^16^. In humans, VKORc1 is an integral membrane protein of the endoplasmic reticulum and catalyzes the reduction of Vitamin K^17^. The reduced Vitamin K is then used by gamma-glutamyl carboxylase to catalyze the conversion of glutamic acid to γ-carboxyglutamic acid in several blood clotting factors^17,18^. This post-translational modification activates the blood coagulation cascade^19^. VKORc1 is indeed the target of oral anticoagulants such as Warfarin (aka Coumadin®) and Bromindione^17,18,20,21^. These anticoagulants, also known as coumarin-based vitamin K antagonists, remain the main therapy to prevent and treat thromboembolic diseases^22^. The mechanism of action of oral anticoagulants has recently been solved by crystallography and involves the formation of two hydrogen bonds with two amino acid residues in human VKOR^23^.

DsbB and VKOR share a common architecture of four transmembrane (TM) helices that form the active site with a quinone binding pocket containing the CXXC motif ^24,25^. However, one difference is the inversion of the locations for CXXC and CX_n_C motifs. In DsbB proteins the CXXC motif is at the beginning of TM2 while in VKOR at the beginning of TM4. The CX_n_C motif is on the opposite side of the enzyme, in DsbB it is located between TM3 and TM4 while in VKOR is between TM1 and TM2. Despite DsbB and VKOR sharing the presence of CX_n_C and CXXC motifs that perform the same covalent reactions with the partner protein and quinone respectively, structural evidence suggests different steps in *de novo* DSB catalysis^14,23,25,26^.

We have previously developed a cell- and target-based assay to find molecules that inhibit DsbB and VKOR proteins of pathogenic bacteria^27,28^. This assay uses *E. coli* cells expressing a β-galactosidase sensor (β-Gal^dbs^) that is a secreted fusion of β-galactosidase to the membrane protein MalF^5,29^. β-Gal^dbs^ is inhibited by DSB formation in the periplasm and is only active in cells with a disabled DSB formation pathway^5,6^. Using an *E. coli* Δ*dsbB* mutant that expresses the β-Gal^dbs^ sensor and is complemented with a plasmid carrying either *Pseudomonas aeruginosa dsbB1* or *Mycobacterium tuberculosis vkor* genes, we perform parallel screens of compounds to identify inhibitors of each of these enzymes^28^. The screens are performed in parallel to provide reciprocal controls that eliminate inhibitors that influence β-Gal activity by acting directly on *E. coli* DsbA or affecting membrane protein assembly because those molecules would likely appear as inhibitors of both strains. Given the differences in the mechanism of *de novo* DSB formation between DsbB and VKOR enzymes, specific inhibitors usually register as hits against one strain or the other^27,28^. However, in rare cases, it is possible that single compounds could target both enzymes by blocking common intermediates presumably shared between the two enzymes.

Using this methodology in previous screens of ∼280,000 synthetic molecules, we found a family of compounds structurally similar to warfarin that inhibited both DsbB and VKOR^28^. We reasoned that this family of drugs could be blocking quinone binding in both enzymes and the structural differences compared to warfarin could reveal common steps in the *de novo* DSB formation by DsbB and VKOR.

Here, we report the functional expression of human VKORc1 mutant derivative in our *E. coli*-based assay system to search for molecules that target the mammalian enzyme. We also performed a structure-activity relationship study of warfarin-like molecules that we identified in a previous *E. coli* screen to understand the differences in selectivity. In an attempt to narrow down the residues binding to these analogs, we discovered that the fourth amino acid residue downstream of the CXXC motif is essential for both DsbB and VKOR enzymes in ubiquinone and menaquinone binding. Our studies highlight a conserved role of this residue in *de novo* disulfide-generating enzymes.

## Results

### Functional expression of human VKORc1 in *E. coli* requires the substitution of an additional positive residue

Bacterial VKOR proteins can complement an *E. coli* strain lacking the *dsbB* gene^12^. However, mammalian VKORc1 homologs could not complement a *dsbB* mutant due to problems with protein insertion into the *E. coli* membrane^30^. Vertebrate VKORc1 proteins have an excess of positive charges in the extracytoplasmic loop between TM segments 1 and 2. This violation of the positive inside rule in which cytosolic loops contain more positively charged amino acids causes protein degradation and impedes proper insertion into the *E. coli* membrane^30,31^. A spontaneous deletion of three amino acids (ΔA31A32R33 aka ΔA31AR), including the deletion of a positive charge from the second loop of the protein, allowed the functional expression of rat VKORc1 (*Rn*VKOR) in *E. coli*^30^. Additional chromosomal mutations that modified the insertase YidC (T362I) and knocked out the protease HslV (C160Y) further stabilized the *Rn*VKOR protein and enhanced expression, which is consistent with the topological insertion problem^30^.

While these modifications worked for *Rn*VKOR, the removal of that same positive charge expressed in the *E. coli* YidC_T362I_ HslV_C160Y_ mutant did not allow human VKORc1 (*Hs*VKOR) to complement DsbB’s function^30^. Since these two proteins share 83% identity (Figure 1A), we converted three residues in the amino terminus of *Hs*VKOR to the ones present in *Rn*VKOR to determine whether these additional mutations together with the previous ΔA31AR deletion would allow *Hs*VKOR to complement an *E. coli* Δ*dsbB* mutant. These mutations introduce a positive charge in loop 1 (W10R) and both, remove a positive and add a negative charge in loop 2 (D36N, R37E) (Figure 1A). We tested the combined mutant (ΔA31AR W10R D36N R37E) by its ability to complement DSB formation in *E. coli* lacking *dsbB* in two backgrounds, one with YidC_T362I_ HslV_C160Y_ previously found to stabilize *Rn*VKOR and the other with wildtype YidC HslV. Disruption of DSB formation leads to a decrease in motility due to FlgI misfolding^32^ and an increase in β-Gal activity when the β-Gal^dbs^ sensor is expressed^5^. We observed that expression from an IPTG-inducible promoter^33^ of *Hs*VKOR with the three additional changes complemented Δ*dsbB* motility in both backgrounds (Figure 1B) while decreased β-Gal^dbs^ activity was only observed in the YidC_T362I_ HslV_C160Y_ background (Figure 1C). This is expected since we have previously observed β-Gal^dbs^ requires more extensive DSB formation activity to inactivate and misfold β-Gal while less activity is required for FlgI folding.

**Figure 1.**
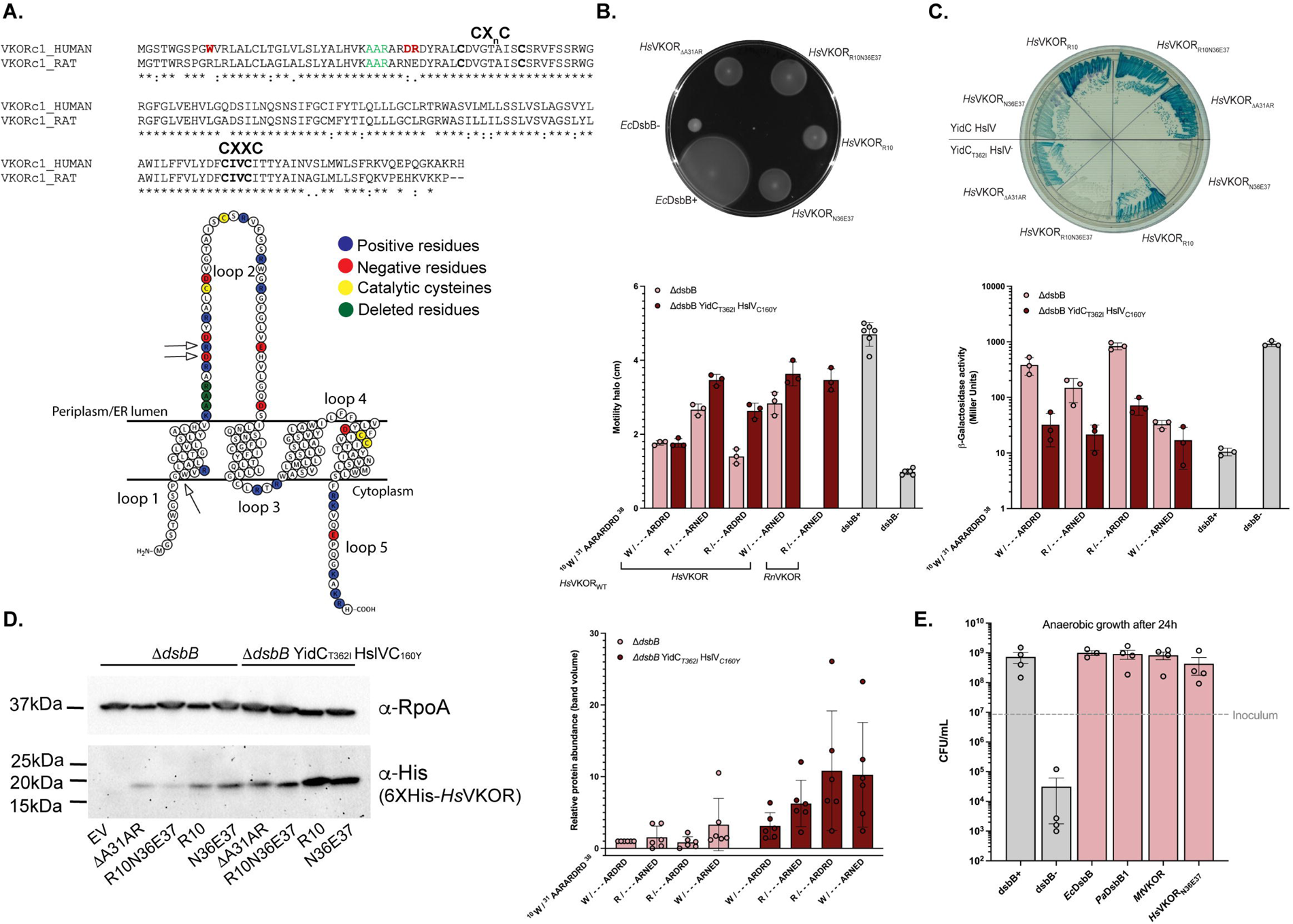
*Hs*VKOR is functional in *E. coli* by mutating two positive residues in the extracytoplasmic loop. A) Alignment of Human (Q9BQB6) and rat (Q6TEK4) VKOR using clustal omega (https://www.ebi.ac.uk/jdispatcher/msa/clustalo). Residues highlighted in red were analyzed in this study. Residues highlighted in green were deleted based on a previous spontaneous mutation. B) *Hs*VKOR variants complement motility of *E. coli* Δ*dsbB* mutant. Swarming halos of *Hs*VKOR mutants expressed in *E. coli* Δ*dsbB* or Δ*dsbB* YidC_T362I_ HslV_C160Y_. Top, a representative image. Bottom, halos measured after 48 h incubation at 30°C. Data represents average±SD of at least three independent experiments. C) *Hs*VKOR variants complement the β-Gal^dbs^ phenotype of *E. coli* Δ*dsbB* YidC_T362I_ HslV_C160Y_ strain. Top, a representative image. Bottom, β-Gal activity was quantified after 18 h growth at 30°C. Data represents average±SD of three independent experiments. D) Mutations in residues 36-37 of *Hs*VKOR help stabilize the protein. Western blotting using anti-His antibody was used to detect *Hs*VKOR variants and anti-RpoA as a loading control. A representative image is shown. The adjusted total band volume of His and RpoA bands was done using ChemiDoc™ MP. Data represents average±SD of six independent experiments. E) *Hs*VKOR_N36E37_ complements anaerobic growth of *E. coli* Δ*dsbB*. Aerobically grown cells were diluted to ∼9x10^6^ CFU/mL (gray dotted line) into anaerobic minimal medium containing 100 mM potassium nitrate. Bacteria were enumerated after 24 h of anaerobic incubation at 37°C. Data represents the average±SEM of at least three independent experiments.

We then independently reverted the mutations to wildtype *Hs*VKOR (R10W and N36D/E37R) to determine the residue(s) responsible for the observed activity. Reverting N36D and E37R while keeping only R10 decreased motility and increased the β-Gal activity compared to the mutant with the three changes, thus indicating that R10 is not the change that allows complementation (Figure 1BC). On the other hand, reverting only R10W while retaining the N36E37 mutations allowed complementation of motility and β-Gal activity in both genetic backgrounds (Figure 1BC). Consequently, the removal of a positive charge and the addition of a negative charge in loop 2 (N36E37) allows *Hs*VKOR to complement *dsbB* in *E. coli*.

We determined the expression of *Hs*VKOR by western blot analysis to see whether the phenotypes observed are due to protein stabilization. Indeed, the amount of protein observed for *Hs*VKOR with N36E37 mutations is 3-fold higher in the wildtype background and 10-fold higher in the YidC_T362I_ HslV_C160Y_ strain compared to only the A31AR deletion (Figure 1D). Overall, this strain has higher *Hs*VKOR protein levels than the wildtype given that the protease HslV is catalytically dead and allows YidC_T362I_ to insert *Hs*VKOR into the *E. coli* membrane. However, the addition of a positive residue (R10) in loop 1 has a relatively equivalent amount of protein to N36E37 mutant in YidC_T362I_ HslV_C160Y_ strain (Figure 1D) but gives no β-Gal^dbs^ complementation (Figure 1C), perhaps due to adopting the incorrect topology upon insertion in the *E. coli* membrane and thus preventing interaction with *E. coli* DsbA. This may suggest that it is the overall number of positive charges in both cytoplasmic and periplasmic loops that prevent the insertion of *Hs*VKOR rather than only the positive charges in the periplasmic loop as previously suggested^30^. Substitution of the two arginine residues (R33 and R37) with glycine residues did not complement the motility phenotype while deleting only the two arginine residues with the neighboring negative charge (D36) produced small halos (Supplementary Figure 1). This suggests that in addition to removing the positive charges, the removal of two additional residues is required to shorten the distance of loop 2, which is responsible for the interaction with DsbA.

*E. coli* uses predominantly ubiquinone-8 (UQ-8) under aerobic growth whereas menaquinone-8 (MK-8) is used under anaerobiosis (Figure 2A)^8,34^. Our above experiments under aerobic growth indicate that *Hs*VKOR when expressed in *E. coli* can use ubiquinone as an electron acceptor. To test whether the *Hs*VKOR variant can also use menaquinone as an electron acceptor we measured growth anaerobically. We proceeded with the best complementing strain henceforward, the *Hs*VKOR_ΔA31AR-D36N-R37E_ expressed in the YidC_T362I_ HslV_C160Y_ strain. While the DsbA and DsbB are dispensable aerobically, under anaerobic conditions the mutants fail to grow due to lack of disulfides in two essential proteins^35^. That is, when we inoculate ∼9x10^6^ CFU/mL from an overnight aerobic culture, the *E. coli dsbB* mutant is not viable after 24 h of anaerobic incubation yielding ∼10^4^ CFU/mL (Figure 1E). However, when we express *Hs*VKOR variant in the *dsbB* mutant, the growth is restored to ∼4x10^8^ CFU/mL (Figure 1E). Therefore, the modified *Hs*VKOR can use both UQ-8 and MK-8 when expressed in *E. coli*. Similarly, the expression of *M. tuberculosis* VKOR (*Mt*VKOR) and *P. aeruginosa* DsbB1 (*Pa*DsbB1) proteins allows anaerobic growth of *E. coli* (Figure 1E).

**Figure 2.**
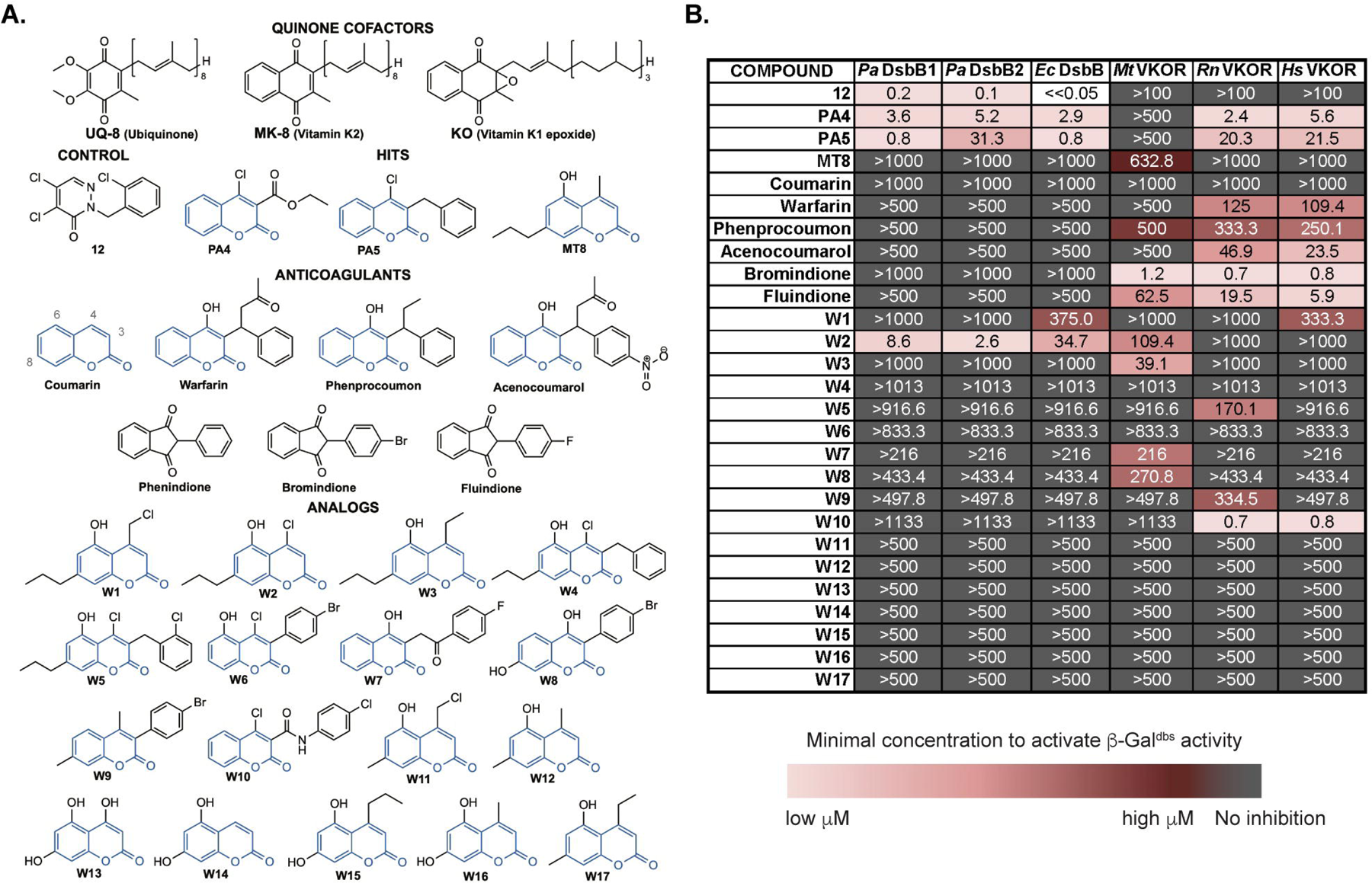
*In vivo* inhibition of DsbB and VKOR enzymes with warfarin analogs using a heterologous *E. coli* platform expressing β-Gal^dbs^. A) Chemical structures of cofactors, anticoagulants, and warfarin analogs tested against DsbB and VKOR. B) Structure-activity relationship analysis of warfarin analogs. *E. coli* Δ*dsbB* strain expressing the β-Gal^dbs^ carrying *P. aeruginosa* (*Pa*) DsbB1, *E. coli* (*Ec*) DsbB, *M. tuberculosis* (*Mt*), rat (*Rn*), or human (*Hs*) VKOR proteins. Strains were grown for 24 h at 30°C in 384-well minimal media plates containing X-Gal and a serial dilution of drugs (often 0 to 500 µM). The minimal concentration of drug to produce a pale blue color was reported as the minimal concentration to activate β-Gal^dbs^. The results represent the average of at least three independent experiments using color coding.

### Anticoagulant screening using *E. coli* heterologous platform

We tested whether the *E. coli* expressing *Hs*VKOR mutant would respond to anticoagulant treatments using the β-Gal^dbs^ sensor. We determined β-Gal activity on 384-well plates using agar minimal media with X-Gal and serial dilutions of several oral anticoagulants including warfarin, phenprocoumon, acenocoumarol, bromindione, and fluindione (Figure 2A). While *E. coli* strains expressing native DsbB or *P. aeruginosa* DsbB1 or DsbB2 proteins show no signs of β-Gal activity at high concentrations of anticoagulants (>500 µM), the *E. coli* strain expressing *Hs*VKOR is inhibited by all anticoagulants in the low to medium micromolar range (Figure 2B). On the other hand, our control compound 12 only targeted strains expressing DsbB but not any of the VKOR-expressing strains as we have shown before^27^ (Figure 2B). Thus, we can detect inhibition of *Hs*VKOR in a high throughput format enabling the identification of new anticoagulants using a bacterial assay.

### Structure activity-relationship analysis of warfarin-like molecules

We have previously identified a family of drugs that target DsbB or VKOR enzymes of pathogenic bacteria^28^. This family of molecules, PA4, PA5, and MT8, have a coumarin ring as a common structure (Figure 2A). Compounds PA4 and PA5 were found as inhibitors of *Pa*DsbB1 but did not target *Mt*VKOR, while compound MT8 inhibits *Mt*VKOR but not *Pa*DsbB1^28^. There are two major differences in the structure, the absence/presence of a hydroxyl group at position 6 or the presence of a chloride/methyl at position 4 in the coumarin ring (Figure 2A). We reasoned these groups could give selectivity towards DsbB or VKOR and thus potentially allow us to also study the mechanism of *de novo* DSB formation of these enzymes using *E. coli* as a test platform. In addition, the structure of these molecules is also shared with warfarin, a 4-hydroxy-coumarin (Figure 2A). Hence, we analyzed the structure-activity relationship (SAR) using commercially available (W11-W17) and rationally designed analogs of coumarin (W1-W10).

We measured the inhibition of DsbB or VKOR expressed in *E. coli* by their ability to activate β-Gal^dbs^ in our high-throughput assay. We found that the combination of chloride at position 4 with a hydroxyl at position 6 (compound W2) gives selectivity towards the *P. aeruginosa* DsbB1 and DsbB2, *E. coli* DsbB and *M. tuberculosis* VKOR but no activity against mammalian VKOR proteins (>130-fold less active, Figure 2B) in the β-Gal^dbs^ assay. The addition of an ethyl group at position 4 combined with a hydroxyl at position 6 (compound W3) gave selectivity against mycobacterial VKOR only. Moreover, maintaining a chloride at position 4 with a hydroxyl at position 6 but adding a phenyl moiety (compounds W4-W6) at position 3 completely abolished the activity against *Pa*DsbB1 and *Mt*VKOR (Figure 2B).

To determine the redox state of the four catalytic cysteines of *Pa*DsbB1 when treated with compound W2, we used *in vivo* alkylation with MalPEG2k which derivatizes the free thiols adding 2 kDa of molecular weight to the protein per cysteine. *Pa*DsbB1 is found in an oxidized state with four catalytic cysteines in a disulfide-bonded state (Figure 3B, lane 3). Upon treatment with compound W2, the migration of the protein indicates that ∼50% of *Pa*DsbB1 is found with one of the four catalytic cysteines reduced (Figure 3, lanes 4-5), meaning that two cysteines are in a disulfide bond and the third one is unavailable potentially forming a covalent adduct with compound W2 (Figure 6C).

**Figure 3.**
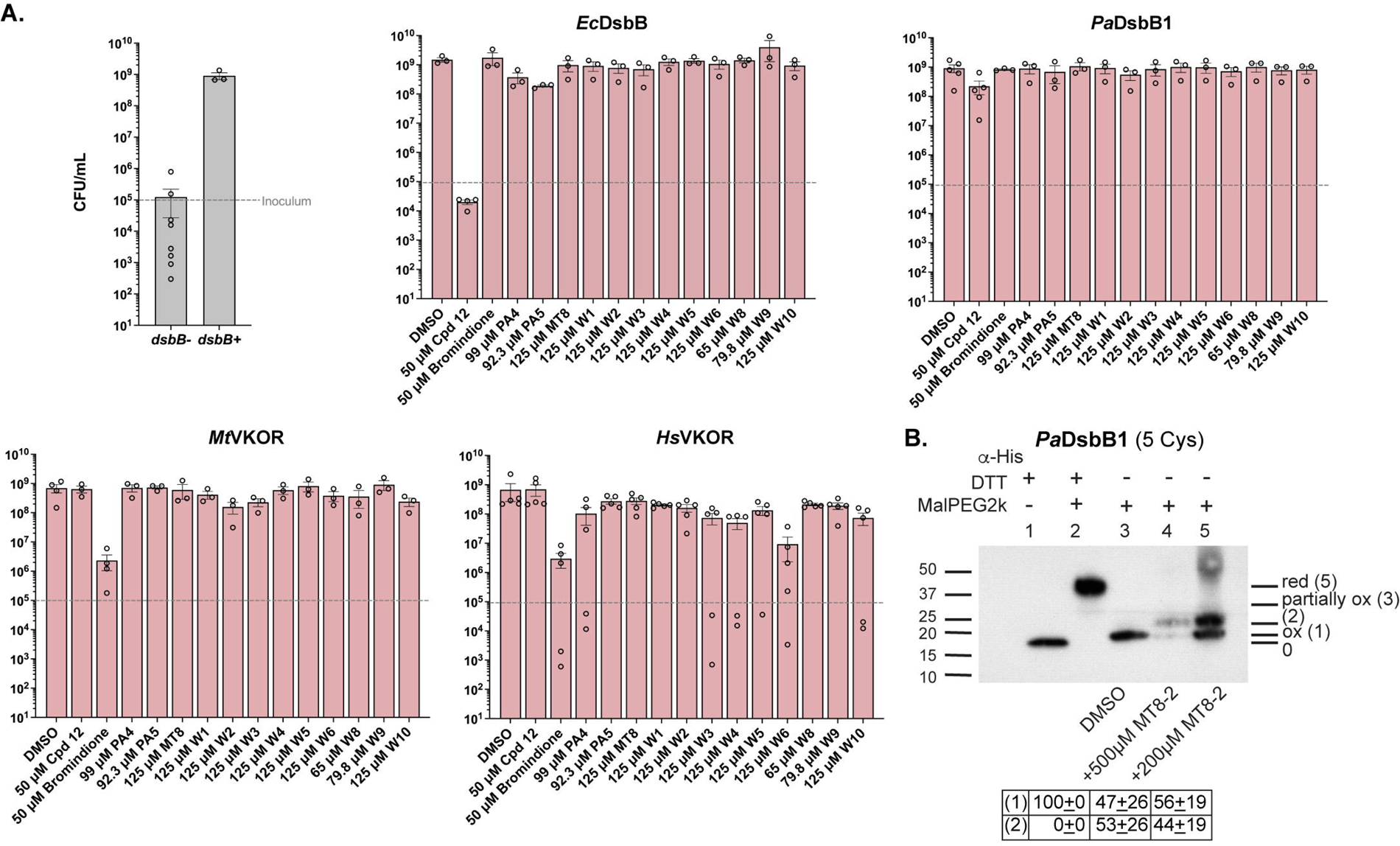
*In vivo* inhibition of DsbB and VKOR enzymes with warfarin analogs using a heterologous *E. coli* platform under anaerobiosis. A) Strains were grown aerobically to mid-log phase and diluted to ∼1x10^5^ CFU/mL (gray dotted line) into minimal medium containing 100 mM potassium nitrate. DMSO-diluted drugs were added to one final concentration with a 0.25% concentration of DMSO. Bacteria were enumerated after 24 h of anaerobic incubation at 37°C. Data represents the average±SEM of at least three independent experiments. B) *In vivo* alkylation was performed in *E. coli* cells expressing *Pa*DsbB1 grown aerobically in minimal medium with DMSO or compound W2. TCA-precipitated proteins were alkylated with 12.5 mM of MalPEG-2kDa. Samples were run by SDS-PAGE and immunoblotted with anti-His antibody. 100 mM dithiothreitol (DTT) was used for reducing DSBs. There are four catalytic cysteines in *Pa*DsbB1 plus one non-catalytic. Ox, oxidized the four catalytic cysteines are forming two DSB; red, reduced all cysteines are in the thiol form; partially ox, two catalytic cysteines reduced and two oxidized. The number in parenthesis indicates the cysteines that are in the free thiol form and were derivatized with MalPEG2k (2kDa addition per cysteine).

We then sought to determine whether these molecules can antagonize MK-8 by testing anaerobic growth inhibition. A single concentration of the analogs dissolved in minimal medium was tested using ∼10^5^ log-phase *E. coli* cells as inoculum. At these concentrations (∼70-100 µM) the analogs failed to inhibit the anaerobic growth of the *E. coli* expressing either bacterial or mammalian enzymes (Figure 3A). However, 50 µM of our controls, compound 12 inhibited *E. coli* DsbB-expressing strain and bromindione inhibited both mycobacterial and human VKOR-expressing strains, therefore our assay can detect inhibition of these enzymes anaerobically (Figure 3A). In addition, we observed that our control compound 12 did not inhibit the anaerobic growth of the *Pa*DsbB1-expressing strain, suggesting it is unable to block MK-8 (see Discussion).

Collectively, compounds W2 and W3 can prevent UQ-8 binding but are less effective or unable to prevent MK-8 binding to *Pa*DsbB1 and *Mt*VKOR enzymes. Similarly, W10 can prevent UQ-8 but less so MK-8 binding to *Hs*VKOR, and vice versa W6 is unable to prevent UQ-8 but can mildly prevent MK-8 binding in *Hs*VKOR. Compound W2 binds covalently to one of the cysteines of *Pa*DsbB1, most likely the quinone-binding cysteine as we have seen previously with compound 12^27^.

### Residues binding to warfarin-like molecules

To find the residues responsible for binding to the warfarin analogs we sought to select DsbB and VKOR variants resistant to them. For this, we generated plasmid libraries of randomly mutagenized *Mtvkor* and *PadsbB1* genes under *trc* and *trc*206 promoters, respectively, and transformed them into the strain deleted of *dsbB* and harboring the β-Gal^dbs^ sensor. Colonies harboring a sensitive enzyme would display a blue color on X-Gal plates with inhibitor, while resistant mutants would be white. Thus, we plated the mutant libraries on X-Gal minimal media with two compounds that we have in sufficient quantities for a screen, 10 μM bromindione for *Mt*VKOR or 8 μM PA5 for *Pa*DsbB1. We then extracted plasmids from the white colonies, transformed them into a fresh strain to ensure functional β-Gal^dbs^, confirmed resistance, and sequenced the gene and the promoter. Our screen with *Mt*VKOR yielded 33 mutants and 16 out of them had changes in residue N81 mutated to either I (5 mutants), Y (5 mutants), D (4 mutants), or S (2 mutants). On the other hand, we were unable to find intragenic *PadsbB1* mutations (see below).

To complement our approach, we used the recent crystal structure of *Hs*VKOR with warfarin to model the binding. Warfarin forms two hydrogen bonds with *Hs*VKOR, one between the 4-hydroxyl group of warfarin with Y139 and the other one between the 2-ketone group of warfarin with N80 of *Hs*VKOR^23^. Similarly, phenindione, a close analog of bromindione (Figure 2A) interacts with *Hs*VKOR by forming hydrogen bonds between the 1,3-diketones of the indandione ring with N80 and Y139^23^. Thus, the reason why we found mutations in N81 residue of *Mt*VKOR, which is the analogous residue of N80 in *Hs*VKOR. Structural alignment between *Hs*VKOR and the AlphaFold^36^ predicted structure of *M. tuberculosis* VKOR highlights conservation between N80 residue while W146 is present instead of Y139 (Figure 4A, left). When we aligned the TM segments that contain the quinone binding pocket of *Hs*VKOR (TM1-2) with the AlphaFold^36^ predicted structure of *P. aeruginosa* DsbB1 (TM3-4), R47 is found in place of Y139 and S147 in place of N81 (Figure 4A, right). Note that Y139, W146, and R47 are located three residues apart from the CXXC motif in both DsbB and VKOR enzymes.

**Figure 4.**
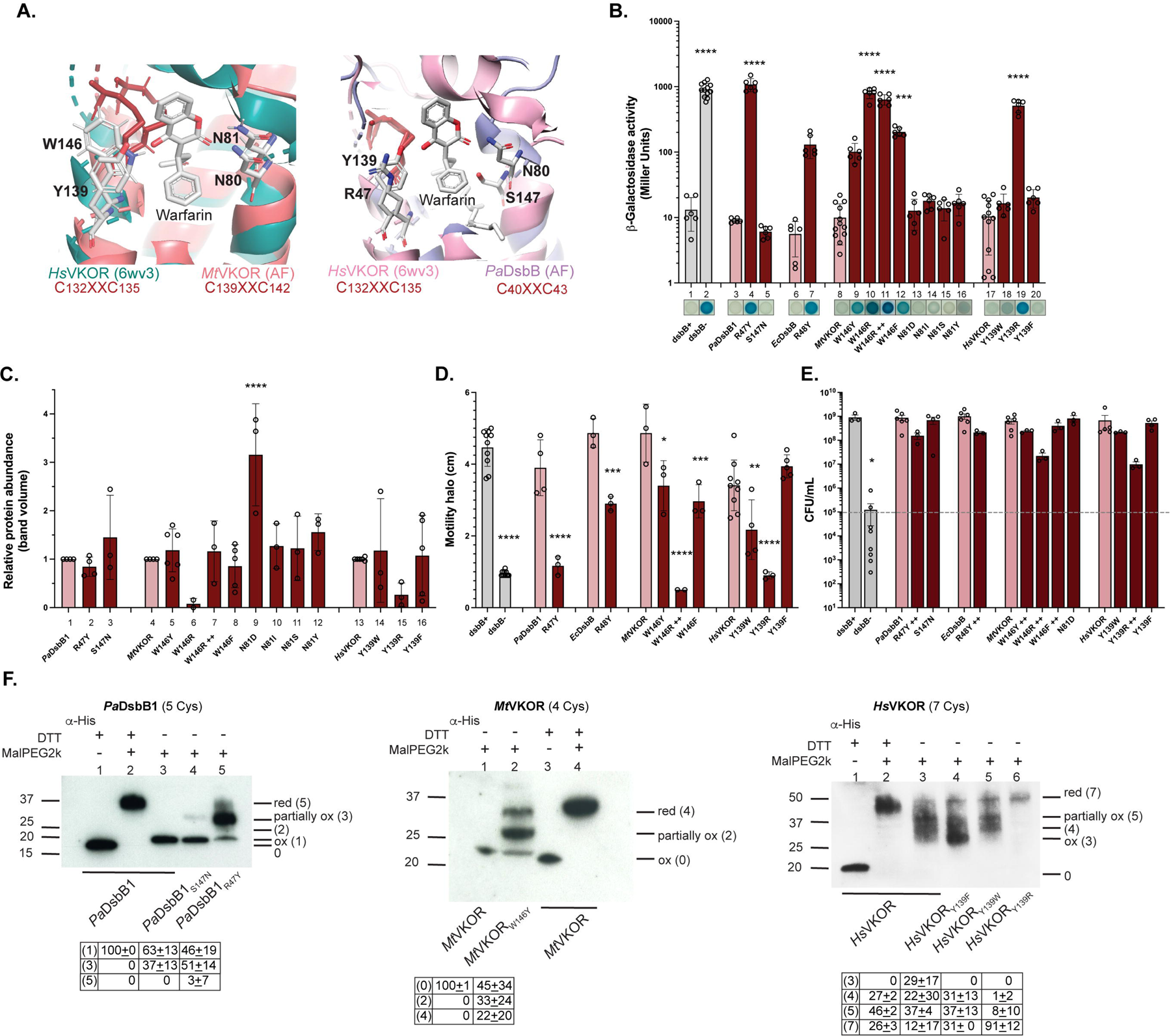
The fourth residue downstream the catalytic CXXC motif is required for *de novo* oxidation of these cysteines in DsbB and VKOR. A) Structural alignment of *Hs*VKOR-warfarin (6wv3) with the AlphaFold^36^ predicted structure of *Mt*VKOR (left, I6X5W1) using PyMOL (The PyMOL Molecular Graphics System, Version 2.0 Schrödinger, LLC). Structural alignment of *Hs*VKOR-warfarin (6wv3, residues 97-155) with the AlphaFold predicted structure of *Pa*DsbB1 (right, Q02TM7, residues 1-60). Catalytic cysteines are highlighted in red, and residues that align with N80 and Y139 of *Hs*VKOR are indicated in both alignments. B) Mutants in the CXXC+4 in both DsbB and VKOR display high β-Gal activity. β-Gal activity in these strains was quantified after 18 h growth at 30°C. A representative image is shown on the X-axis. ++ indicates more IPTG. Data represents average±SD of at least three independent experiments. C) Protein abundance across CXXC+4 mutants was determined by western blot using anti-His antibody and anti-RpoA as loading control compared to their wildtype. Data represents average±SD of at least three independent experiments. D) CXXC+4 mutants are less motile than the wildtype enzymes. Data represents average±SD of at least three independent experiments. E) CXXC+4 mutants grow anaerobically. Log-phase aerobically grown cells were diluted to ∼1x10^5^ CFU/mL (gray dotted line) into anaerobic minimal medium containing 100 mM potassium nitrate. Bacteria were enumerated after 24 h of anaerobic incubation at 37°C. Data represents the average±SEM of at least three independent experiments. Statistical tests were done using Ordinary one-way ANOVA multiple comparisons. p-value ≤0.0001 (****), 0.0002 (***), 0.021 (**), 0.0332 (*), and ns ( ). F) CXXC+4 mutants are found more predominantly in partially reduced states. *In vivo* alkylation was performed in cells grown aerobically in minimal medium. TCA-precipitated proteins were alkylated with 12.5 mM of MalPEG2k. Samples were run by SDS-PAGE and immunoblotted with anti-His antibody. 100 mM dithiothreitol (DTT) was used for reducing DSBs. The total amount of cysteines present in each protein is indicated in parenthesis. Images are representative of at least two independent experiments. There are four catalytic cysteines in each protein plus one non-catalytic in *Pa*DsbB1 or three non-catalytic cysteines in *Hs*VKOR. Ox, oxidized the four catalytic cysteines are forming two DSB; red, reduced all cysteines (catalytic and non-catalytic) are in the thiol form; partially ox, two catalytic cysteines reduced and two oxidized. The number in parenthesis indicates the cysteines that are in the free thiol form and were derivatized with MalPEG (2kDa addition per cysteine).

### CXXC+4 residue is important for quinone binding in both DsbB and VKOR proteins

Kobayashi *et al*., have previously altered the fourth residue downstream of the quinone-binding motif (hereafter CXXC+4) of *Ec*DsbB by changing the R48A and observed no effect on the DsbB redox state but a slight defect on the oxidation of DsbA^37^. Separately Kadokura *et al*., also isolated weak Lac^+^ mutants of β-Gal^dbs^ in this residue including R48H and R48C^38^. The mutants functioned reasonably well aerobically where DsbB is found mostly in the oxidized form, but poorly semi-aerobically where DsbB mutants were found in the reduced state. Indeed, the R48H mutant displayed a reduced ability to bind ubiquinone and negligible binding to menaquinone *in vitro*^38^. Additionally, patients who are resistant to the anticoagulant effects of warfarin map to changes of Y139F in *Hs*VKOR, which indicates the enzyme is still functional^17,39^. Thus, CXXC+4 may have been studied separately in DsbB and VKOR enzymes but the role of this residue in binding to quinones could be critical in the differences we observed with warfarin-like inhibitors. We reasoned that if we swapped the CXXC+4 and N80 analogous residues between DsbB and VKOR proteins, we would be able to see differences in the inhibition/resistance to the warfarin analogs. We thus mutated the two residues found in the *P. aeruginosa* DsbB1 to those found in *Hs*VKOR (R47Y and S147N), and the residue found in *M. tuberculosis* VKOR protein to that of *Hs*VKOR (W146Y). To our surprise, the mutants in the CXXC+4 were Lac^+^ for both DsbB and VKOR (Figure 4B, 4 and 9), perhaps the reason why our selections were unable to identify mutants in this residue. In comparison, the N81D (or I, S, Y) and S147N mutants were Lac^-^, similar to their wildtype counterparts (Figure 4B, 5 and 13-16). The substitutions R48Y in *Ec*DsbB (Figure 4B, 7) and Y139R in *Hs*VKOR (Figure 4B, 19) also displayed higher β-Gal activity. Changes to hydrophobic residues in *Hs*VKOR had fewer defects (Figure 4B, 18 and 20). This is consistent with *Hs*VKOR_Y139F_ variant being also functional in humans^17^. However, *Mt*VKOR with mutations in W146 to Y, R, or F were unable to complement the lac^+^ phenotype of the *dsbB* mutant (Figure 4B, 9, 10, and 12).

We evaluated the protein expression levels produced by these strains using anti-His antibody, and we found that *Mt*VKOR_W146R_ and *Hs*VKOR_Y139R_ were not sufficiently produced compared to wildtype (see Discussion). Thus, we increased to 10 times the concentration of IPTG (25 µM, indicated as ++) to increase the expression of *Mt*VKOR_W146R_ and achieved similar wildtype protein levels to test for activity (Figure 4C, 6 vs 7). Even under these conditions, the *Mt*VKOR_W146R_ mutant was unable to revert the Lac^+^ phenotype of the β-Gal^dbs^ (Figure 4B, 11). As for *Hs*VKOR, we were unable to induce with more IPTG because it failed to grow perhaps due to toxicity. When inducing on X-Gal minimal media plates with higher IPTG concentrations the phenotypes of all except *Pa*DsbB_R48Y_, *Mt*VKOR_W146R,_ and *Hs*VKOR_Y139R_ were restored (Supplementary Figure 2A). We also tested motility and all mutants except *Hs*VKOR_Y139F_ displayed less motility compared to their WT counterparts (Figure 4D) and similarly more induction increased motility except for *Pa*DsbB_R48Y_, *Mt*VKOR_W146R_ and *Hs*VKOR_Y139R_ mutants (Supplementary Figure 2B). Overall, the changes made in CXXC+4 of DsbB and VKOR, except for the Y139F change in *Hs*VKOR, made the enzymes less active, and changing W/Y residues to R destabilized the proteins.

We then investigated the *in vivo* redox state of the catalytic cysteines (CXXC and CX_n_C motifs) which are oxidized in wildtype DsbB^9,40^. *In vivo* alkylation revealed that only 46% of the total *Pa*DsbB1_R47Y_ is found oxidized, while 51% is partially oxidized (two catalytic cysteines reduced and two oxidized) and 3% is fully reduced (Figure 4F, lanes 5 vs 3). Also, *Pa*DsbB1_S147N_ shows a slight accumulation (37%) of the partially oxidized state (Figure 4F, lanes 4 vs 3) even though it fully complements the phenotype (Figure 4, column 5) thus suggesting that this residue also participates in quinone binding. Similarly, *Mt*VKOR_W146Y_ shows 33% partially oxidized and 22% reduced states compared to the wildtype found in the oxidized form (Figure 4F, lane 2 vs 1). As for *Hs*VKOR, it is found in the three redox states previously identified in human cells^41^; 45% is in partially oxidized state together with 24% in reduced state (Figure 4F, lane 3). *Hs*VKOR_Y139W_ displays similar states to the modified *Hs*VKOR (Figure 4F, lanes 5 vs 3), while 91% of *Hs*VKOR_Y139R_ is found in the reduced state (Figure 4F, lane 6). Surprisingly, 29% of *Hs*VKOR_Y139F_ is found in the oxidized state with 37% in partially oxidized, and 10% in reduced form (Figure 4F, lane 4). Therefore, the redox states observed suggest that the mutants are less active because they are unable to transfer electrons to UQ-8 and form *de novo* a disulfide.

Finally, to test whether the mutants were also unable to bind menaquinone, we used anaerobic growth as a measure of MK-8 binding. All mutants grew less compared to their wildtype counterparts (Supplementary Figure 2C). Once again, we supplemented more IPTG to rescue the growth by expressing more protein, and all except *Mt*VKOR_W146R_ and *Hs*VKOR_Y139R_ were able to reach close to wildtype CFU yields (Figure 4E). Although the *Mt*VKOR_W146R_ and *Hs*VKOR_Y139R_ mutants gave 10-fold less bacterial yield, they were able to grow despite the drastic phenotypes observed aerobically. Therefore, mutations in the CXXC+4 also affected menaquinone binding and overexpression can compensate for the decrease in function.

### Mutations in the CXXC+4 residue not only confer resistance but sensitivity to warfarin-analogs

Our β-Gal^dbs^ assay using the CXXC+4 mutants would fail to detect drug sensitivity because the mutants display blue color and our assay uses this readout for drug inhibition. We instead used the anaerobic growth assay to determine the susceptibility of the warfarin-like analogs. We found that the inhibition pattern of all CXXC+4 mutants changed drastically, the mutants were more prone to be antagonized by 6-hydroxy-4-chloro-coumarins with phenyl substitutions at position 3 (W4, W5, W6, and W10), but not when they lacked it as in compound W2 (Figure 5). On the other hand, *Hs*VKOR_Y139F_ together with *Mt*VKOR_W146F_ were more resistant to Bromindione (Figure 5) as expected based on previous findings^17,39^. In contrast, mutants S147N of *Pa*DsbB1 and N81D of *Mt*VKOR were not affected by these molecules as the CXXC+4 mutants were. Consequently, changes in the CXXC+4 residue can not only confer resistance but also sensitivity to 4-chloro-coumarins with substitutions at position 3.

**Figure 5.**
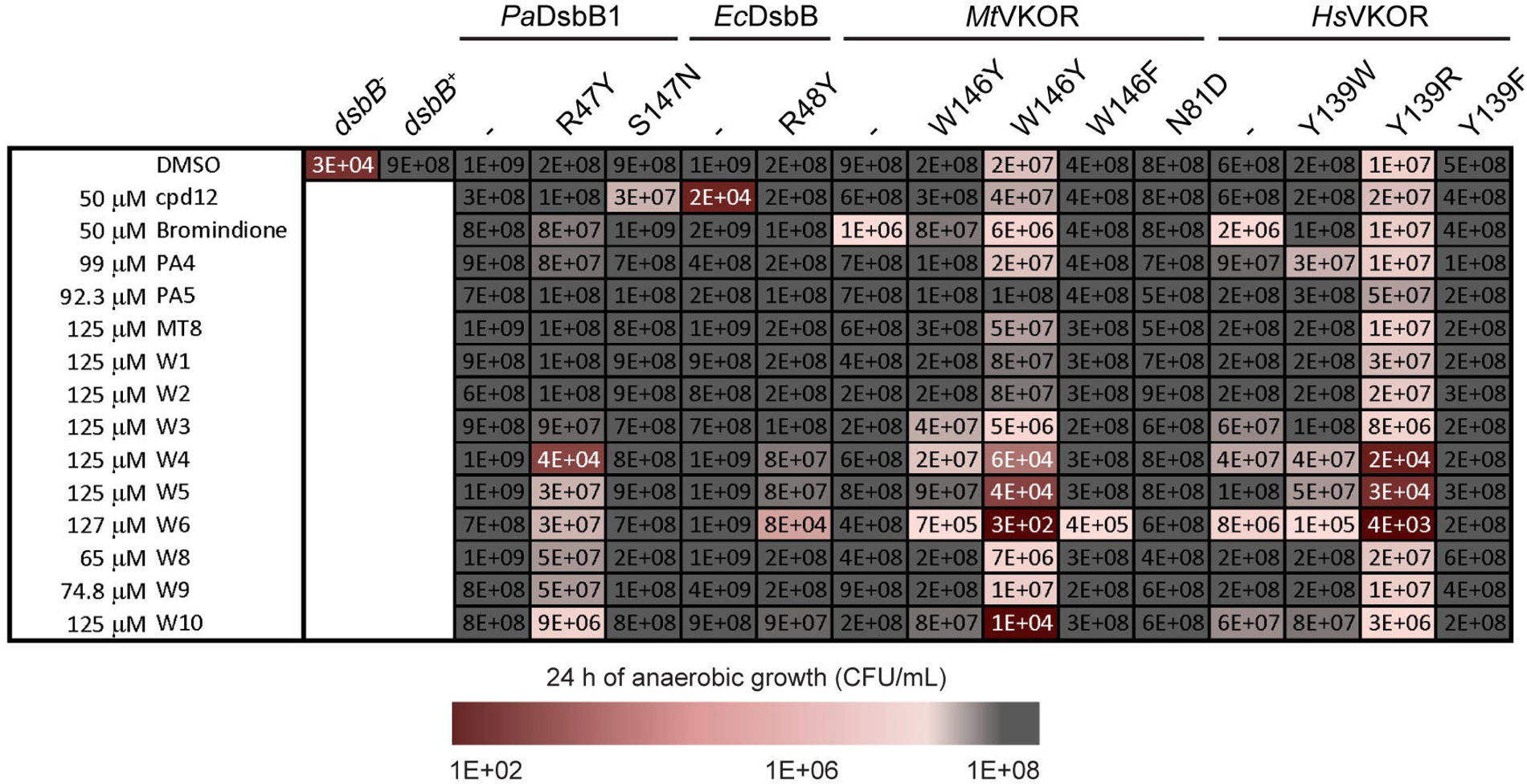
CXXC+4 mutants are sensitized to 4-chloro-coumarin analogs with substitutions at position 3. Log-phase aerobically grown cells were diluted to ∼1x10^5^ CFU/mL into anaerobic minimal medium containing 100 mM potassium nitrate. DMSO-diluted drugs were added to one final concentration indicated (final 0.25% DMSO). Bacteria were enumerated after 24 h of anaerobic incubation at 37°C. The results represent the average of at least three independent experiments using color coding. See Supplementary Figure 3 for complete data.

## Discussion

We have functionally expressed human VKORc1 in *E. coli* by removing a positive residue in the first extracytoplasmic loop in addition to a spontaneous deletion of three amino acids that also removes a second positive residue in this loop and has previously helped to stabilize rat VKORc1^30^. These changes decrease the excess of positive residues outside of the cytoplasmic membrane that violate the positive-inside rule, hence stabilizing human VKORc1 and allowing its proper insertion into the *E. coli* membrane. In addition, the removal of three residues was also necessary for complementation, possibly shortening the first extracytoplasmic loop is required to allow productive interaction with *E. coli* DsbA. In humans, VKORc1 participates in two reduction steps involving four electrons, two electrons from vitamin K epoxide to vitamin K and two more from vitamin K to the hydroquinone form^25^. However, the expression of the modified human VKORc1 in *E. coli* participates only in the last reduction step involving two electrons from ubiquinone (aerobically) or menaquinone (anaerobically) to the hydroquinone form allowing the oxidation of DsbA. The epoxide form is produced by gamma-glutamyl carboxylase in vertebrates and is not known to be produced in bacteria.

We have also shown that, unlike *E. coli* DsbB-based strains, the *E. coli* expressing the modified human VKORc1 is sensitive to coumarin-derived oral anticoagulants and this strain could enable the search for novel anticoagulants using our previously developed high throughput β-Gal assay^27^. Additionally, the high throughput β-Gal assay may allow us to find molecules that target the *Hs*VKOR_Y139F_ variant found in patients who do not respond to warfarin treatment, and to determine whether other Y139 variants are more resistant or sensitive to current oral anticoagulants.

We performed a SAR analysis of warfarin-like molecules that we found in a previous *E. coli* screen targeting independently the bacterial VKOR or DsbB enzymes^28^. It seemed unlikely to identify a single drug that inhibits both DsbB and VKOR enzymes due to their differences in the mechanism of *de novo* DSB formation. However, we have found a class of molecules represented by compound W2, with a coumarin ring as the common structure that can target both bacterial DsbB and VKOR and prevent electron transfer to ubiquinone. The antagonism of this class of drugs seems to be only against ubiquinone rather than menaquinone, in principle, this could be an advantage since it could guide bacterial inhibition towards one lifestyle rather than the other (for instance anaerobic vs aerobic growth). However, further analysis is needed to understand the reason for this difference. Perhaps such studies could reveal a difference in an intermediate state between ubiquinone vs menaquinone that these inhibitors block. Further substitutions at positions 3 and 4 of the coumarin ring could lead to increased efficacy and selectivity against bacterial or mammalian enzymes.

In an attempt to elucidate which residues associate with these coumarin analogs, we discovered that the fourth residue downstream the CXXC motif is essential for both DsbB and VKOR enzymes in ubiquinone and menaquinone binding. Our screens to isolate resistant mutants did not allow us to determine the CXXC+4 binding residue given that these mutants, which were probably the ones allowing stronger resistance, were most likely not functional in the β-Gal^dbs^ assay. However, we were able to identify bromindione-resistant mutants in N81 residue of *Mt*VKOR because modifications in this residue are still functional.

The CXXC+4 mutants with the swapped-residues between DsbB and VKOR were sensitive to the same class of drugs (compounds W4 and W6) during electron transfer to menaquinone. It is tempting to speculate that a similar intermediate in the DsbB and VKOR mutants is formed and targeted by these molecules. Interestingly, we also found that *Ec*DsbB_R48Y_ is more resistant to compound 12 anaerobically. Compound 12, a drug that we previously discovered, inhibits only DsbB enzymes by reacting with C44 of DsbB. The covalent interaction occurs only when the charge transfer complex of *E. coli* DsbA-DsbB with ubiquinone is formed but not when DsbB is fully oxidized^27^. In this state, C44 would be more exposed allowing compound 12 to bind covalently to it. The fact that *Ec*DsbB_R48Y_ mutant is more resistant to compound 12 indicates that it has a decreased ability to form the charge transfer complex with MK-8, hence the inhibitor is not as effective. It is possible that the other CXXC+4 mutants may be behaving this way too, decreasing their ability to stabilize the charge-transfer complex with MK-8. Unexpectedly, 50 µM of compound 12 did not inhibit the anaerobic growth of the *Pa*DsbB1-expressing strain, while it inhibits aerobically at low concentrations (0.2 µM). Compound 12 blocks both DsbB homologs in *P. aeruginosa* grown aerobically^27,28^, indeed the major quinone is UQ-9 in both aerobic and anaerobic growth^42^. The observation that compound 12 inhibits the electron transfer of *Pa*DsbB1 to ubiquinone but less so to menaquinone could be the consequence of differences in the formation of the charge-transfer complex between *E. coli* and *P. aeruginosa* DsbB proteins with menaquinone. In agreement with this, the *Pa*DsbB1_R47Y_ mutant was more severely affected in anaerobic growth, β-Gal^dbs^ activity, and motility than *Ec*DsbB_R48Y_ mutant. Additionally, *Pa*DsbB1 has hydrophobic as opposed to charged residues in the catalytic triad, that is, I46 instead of E47 next to the conserved R47 residue, while L91 instead of H91 near the catalytic C43 of the CXXC motif. Thus, *Pa*DsbB1 may potentially work similar to the hydrophobic residue mechanism of VKOR enzymes.

The CXXC+4 residue is a highly conserved residue, an arginine, among DsbB proteins (Figure 6A); while not as conserved in VKOR enzymes and often A, W, or Y (Figure 6B), probably due to VKOR being more widely distributed in bacteria, eukaryotes, and archaea. A recent crystal structure of *Ec*DbsB revealed a spatial arrangement of a catalytic triad like the one for cysteine or serine proteases^43^. First, E47 residue of *Ec*DsbB (I46 in *Pa*DsbB1) assists deprotonation of H91 (L91 in *Pa*DsbB1), then deprotonated H91 abstracts the proton from the sulfhydryl group of C44, generating a nucleophile to attack the quinone. This facilitates the formation of the cysteine-quinone charge-transfer complex (Figure 6A). The existence of a transient charge-transfer complex between the C44 thiolate and ubiquinone/menaquinone was identified by the appearance of a characteristic pink/purple color when *Ec*DsbB interacts with DsbA_C33S_ mutant, which stabilizes the complex^10,11^. The negative charge of this complex is consequently stabilized by R48 (Figure 6A), which interacts with the opposite side of the quinone ring as opposed to the C44 thiolate and helps polarize the ring to form the charge-transfer complex^25,43^. Hence, R48 mutants of *Ec*DsbB have previously been shown to result in accumulation of the DsbA-DsbB complex, reduced quinone binding, and low quinone reduction activity^37,38^. Correspondingly, the *Pa*DsbB1_R47Y_ mutant was found mostly in a partially oxidized state, with two cysteines reduced and two in oxidized state, thus unable to form a *de novo* DSB.

**Figure 6.**
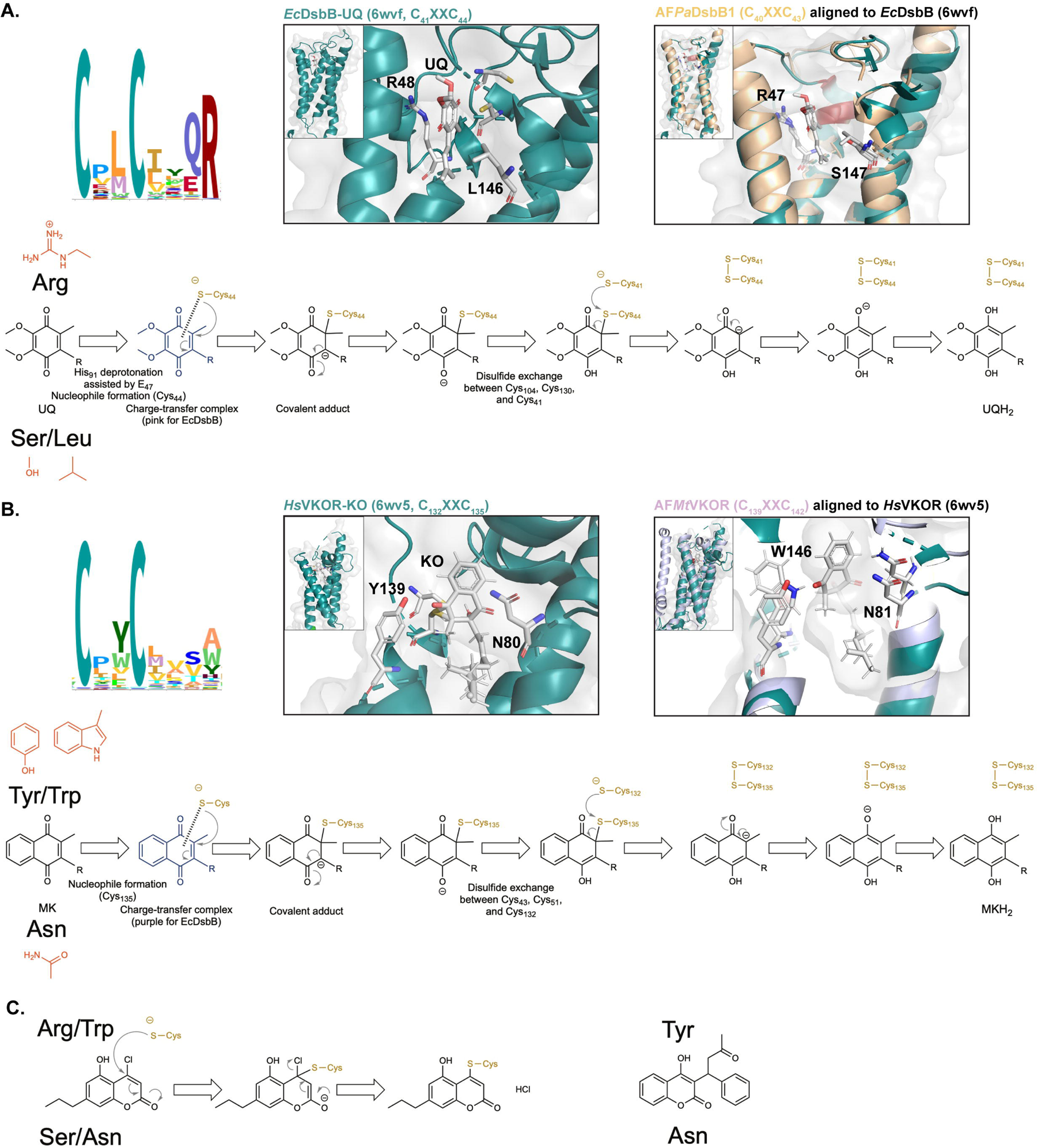
Role of CXXC+4 residue in both DsbB and VKOR enzymes is conserved and stabilizes the quinone intermediates. See text for details. HMM logo of the Pfam seed models for DsbB (PF02600) and VKOR (PF07884) visualized with Skylign (https://www.ebi.ac.uk/interpro/)^44^. A) Structural alignment of *Ec*DsbB-UQ (6wvf) with the AlphaFold^36^ predicted structure of *Pa*DsbB1 (Q02TM7). B) Structural alignment of *Hs*VKOR-KO (6wv5) with the AlphaFold^36^ predicted structure of *Mt*VKOR (I6X5W1). Alignments were done using PyMOL (The PyMOL Molecular Graphics System, Version 2.0 Schrödinger, LLC). Residues important for quinone and warfarin-like molecules are highlighted. Quinone intermediates are shown below alignments, residues that stabilize these intermediates are shown at the top and bottom but they do not mean the structural position. Models were generated with ChemDraw 22.2.0^45^ using models from Ref.^25,46^. C) Proposed mechanism of binding of DsbB and VKOR enzymes to compound W2 (left) based on *Hs*VKOR with warfarin (right) as previously reported^23^.

In contrast to the DsbB proteins, vertebrate VKOR enzymes use two conserved residues N80 and Y139 which form two hydrogen bonds to the 1,4-diketone groups of the naphthoquinone ring of vitamin K quinone (or epoxide), and these same residues help with warfarin interaction (Figure 6BC)^23^. These hydrogen bonds are thought to stabilize the transition states. Substituting Y139 in *Hs*VKOR by another hydrophobic residue like F or W produces a functional enzyme even though *Hs*VKOR_Y139F_ displayed more oxidized form than wildtype or Y139W. However, similar changes in *Mt*VKOR, W146F, and W146Y were more detrimental to its activity. This was consistent with the redox states observed for *Mt*VKOR_W146Y_ which showed reduced and partially oxidized states, while only oxidized is observed in wildtype. Oxidized *Hs*VKOR is not functional^25,41,43^ however, a third of *Hs*VKOR_Y139F_ is found in the oxidized state in *E. coli* and this strain complemented β-Gal^dbs^, motility, and anaerobic growth like the wildtype. It would be interesting to test whether this mutant behaves in this way in humans. In addition, *Mt*VKOR_W146F_ is resistant to all except compound W6 whereas *Hs*VKOR_Y139F_ confers resistance to all analogs. Our data suggests that the mechanism of quinone binding between bacterial and human VKOR enzymes proceeds differently, mycobacterial VKOR is unable to work with other hydrophobic CXXC+4 residues while human VKORc1 tolerates a variety of changes.

We propose that the CXXC+4 residue in DsbB and VKOR enzymes performs a parallel mechanism of stabilization of the quinone intermediates (Figure 6). Thus, R47 in *Pa*DsbB1, R48 in *Ec*DsbB, Y139 in *Hs*VKOR, or W146 in *Mt*VKOR may help to stabilize the benzoquinone (in ubiquinone) or naphthoquinone (in menaquinone/vitamin K quinone or epoxide) ring either through charge or hydrogen bonds (Figure 6). While DsbB and VKOR enzymes may have evolved convergently, our studies highlight a conserved role of the CXXC+4 residue in *de novo* disulfide-generating enzymes. Possibly the position of this residue and its role has been conserved due to the distance between the quinone-binding cysteine and the size of the benzoquinone/naphtoquinone rings.

Lastly, *Hs*VKOR_Y139R_ and *Mt*VKOR_W146R_ mutants were significantly less expressed, while *Mt*VKOR_N81D_ mutant displayed a three-fold increase. What is the reason for these differences? One possibility is that the presence of positive charges in the extracytoplasmic space destabilizes the proteins as we have shown here. However, *Hs*VKOR is already expressed in an *E. coli* strain that can stabilize the excess of positive charges. Another possibility is that during the biogenesis of these membrane proteins, the quinone has to be attached when inserted into the membrane. Thus, since Y139R and W146R are not interacting properly with the quinone they get degraded and this would imply the reverse, that N81D may have more stably attached the quinone. To investigate the stabilization of these mutants, one could select suppressor mutants with more stable *Hs*VKOR_Y139R_ or *Mt*VKOR_W146R_ proteins using the β-Gal^dbs^ phenotype, selecting for white colonies and identifying the genes involved.

Even though the DSB formation pathway was discovered over 30 years ago^5,6^, the mechanism of *de novo* disulfide-generating enzymes has not been fully elucidated^13,25,26,43^. Using bacteria and small molecules could contribute to our understanding of the intermediate states required in disulfide generation by DsbB and VKOR enzymes, and lead to new drugs that can target these mechanisms for both classes of enzymes.

### Significance

The results of this work indicate that disulfide-generating enzymes while acting with two different mechanisms of *de novo* disulfide bond formation can be inhibited by the same class of drugs. Our structure-activity studies set the basis for designing novel anticoagulants or antibacterials using coumarin as a scaffold. Our work also emphasizes a conserved role of the fourth residue after the catalytic motif in *de novo* disulfide-generating enzymes. The nature of this residue in disulfide-generating enzymes can guide the design of novel coumarin analogs.

## Acknowledgments

We thank Laura McPartland for the initial substructure search to find compounds W11-W17 in chemical libraries. We thank Shangfeng Li and Baojian Pei at Bioduro-Sundia for their chemical synthesis efforts. We also thank Clay Fuqua for his helpful comments and suggestions for this manuscript. D.C. was supported by Indiana University’s Cox Research Scholar program. This work was supported by Indiana University Bloomington and by the Cystic Fibrosis Foundation Pilot and Feasibility Award 004846I222 (to C.L.).

## Author’s contributions

Conceptualization, C.L.; Methodology, C.L.; Investigation, D.C., G.N.A., A.M.S.; D., K.R., and C. L.; Resources, L.Z.; Writing – Original Draft, C.L.; Writing – Review & Editing, C.L.; Visualization, C.L.; Supervision and Project Administration, C.L.; Funding Acquisition, C.L.

## STAR Methods

### Bacterial strains and growth conditions

The strains and plasmids used in this study are listed in Table 1. The primers used in this study are listed in Table 2.

**Table 1.** List of strains.

| ID | Genotype | Reference |
| --- | --- | --- |
| *E. coli* host strains |  |  |
| HK325 | HK295 $\Delta dsbB$ λMalFLacZ102, Km<sup>r</sup> | 38 |
| FSH231 | HK295 $\Delta dsbB$ λMalFLacZ102 YidC<sub>T362I</sub> HslVC<sub>160Y</sub> Km<sup>r</sup>, Cm<sup>r</sup> | 30 |
| *HsVKOR* variants |  |  |
| LL238 | HK295 $\Delta dsbB$ λMalFLacZ102 pDSW204-6XHis-*HsVKOR*<sub>$\Delta A31AR$</sub> | This study |
| LL234 | HK295 $\Delta dsbB$ λMalFLacZ102 pDSW204-6XHis-*HsVKOR*<sub>$\Delta A31AR$</sub><br>W10R D36N R37E | This study |
| LL218 | HK295 $\Delta dsbB$ λMalFLacZ102 pDSW204-6XHis-*HsVKOR*<sub>$\Delta A31AR$</sub><br>W10R | This study |
| LL219 | HK295 $\Delta dsbB$ λMalFLacZ102 pDSW204-6XHis-*HsVKOR*<sub>$\Delta A31AR$</sub> D36N<br>R37E | This study |
| LL239 | HK295 $\Delta dsbB$ λMalFLacZ102 YidC<sub>T362I</sub> HslVC<sub>160Y</sub> pDSW204-6XHis-*HsVKOR*<sub>$\Delta A31AR$</sub> | This study |
| LL144 | HK295 $\Delta dsbB$ λMalFLacZ102 YidC<sub>T362I</sub> HslVC<sub>160Y</sub> pDSW204-6XHis-*HsVKOR*<sub>$\Delta A31AR$</sub> W10R D36N R37E | This study |
| LL235 | HK295 $\Delta dsbB$ λMalFLacZ102 YidC<sub>T362I</sub> HslVC<sub>160Y</sub> pDSW204-6XHis-*HsVKOR*<sub>$\Delta A31AR$</sub> W10R | This study |
| LL236 | HK295 $\Delta dsbB$ λMalFLacZ102 YidC<sub>T362I</sub> HslVC<sub>160Y</sub> pDSW204-6XHis-*HsVKOR*<sub>$\Delta A31AR$</sub> D36N R37E | This study |
| LL351 | HK295 $\Delta dsbB$ λMalFLacZ102 YidC<sub>T362I</sub> HslVC<sub>160Y</sub> pDSW204-6XHis-*HsVKOR*<sub>$\Delta R33 \Delta R37$</sub> | This study |
| LL340 | HK295 $\Delta dsbB$ λMalFLacZ102 YidC<sub>T362I</sub> HslVC<sub>160Y</sub> pDSW204-6XHis-*HsVKOR*<sub>$\Delta R33 \Delta R37$</sub> W10R | This study |
| LL426 | HK295 $\Delta dsbB$ λMalFLacZ102 YidC<sub>T362I</sub> HslVC<sub>160Y</sub> pDSW204-6XHis-*HsVKOR*<sub>$\Delta R33 \Delta D36 \Delta R37$</sub> | This study |
| LL428 | HK295 $\Delta dsbB$ λMalFLacZ102 YidC<sub>T362I</sub> HslVC<sub>160Y</sub> pDSW204-6XHis-*HsVKOR*<sub>R33G R37G</sub> | This study |
| LL427 | HK295 $\Delta dsbB$ λMalFLacZ102 YidC<sub>T362I</sub> HslVC<sub>160Y</sub> pDSW204-6XHis-*HsVKOR*<sub>$\Delta R33$</sub> R37N | This study |
| β-Gal strains for drug testing | | |
| CL499 | HK295 λMalFLacZ102 pTrc99a | 27 |
| CL379 | HK295 $\Delta dsbB$ λMalFLacZ102 pTrc99a | 27 |
| CL523 | HK295 $\Delta dsbB$ λMalFLacZ102 pDWS206-*PaDsbB1* | 28 |
| CL382 | HK295 $\Delta dsbB$ λMalFLacZ102 pTrc99a-6XHis-*MtVKOR* | 28 |
| CL377 | HK295 $\Delta dsbB$ λMalFLacZ102 pDWS206-*PaDsbB2*-6XHis | 27 |
| FSH250 | HK295 $\Delta dsbB$ λMalFLacZ102 YidC<sub>T362I</sub> HslVC<sub>160Y</sub> pTrc99a-*RnVKOR*-$\Delta A31AR$ | 30 |
| CXXC+4 mutants |  |  |
| LL395 | HK325 $\Delta dsbB$ λMalFLacZ102 pDWS206-*EcDsbB* | This study |
| LL396 | HK325 $\Delta dsbB$ λMalFLacZ102 pDWS206-*EcDsbB*<sub>R48Y</sub> | This study |
| LL318 | HK325 $\Delta dsbB$ λMalFLacZ102 pDWS206-*PaDsbB1*-6XHis | This study |
| LL353 | HK325 $\Delta dsbB$ λMalFLacZ102 pDWS206-*PaDsbB1*<sub>R47Y</sub>-6XHis | This study |
| LL393 | HK325 $\Delta dsbB$ λMalFLacZ102 pDWS206-*PaDsbB1*<sub>S147N</sub>-6XHis | This study |
| LL250 | HK325 $\Delta dsbB$ λMalFLacZ102 pTrc99a-6XHis-*MtVKOR*<sub>N81D</sub> | This study |
| LL251 | HK325 $\Delta dsbB$ λMalFLacZ102 pTrc99a-6XHis-*MtVKOR*<sub>N81Y</sub> | This study |
| LL241 | HK325 $\Delta dsbB$ λMalFLacZ102 pTrc99a-6XHis-*MtVKOR*<sup>-N81I</sup> | This study |
| LL242 | HK325 $\Delta dsbB$ λMalFLacZ102 pTrc99a-6XHis-*MtVKOR*<sup>-N81S</sup> | This study |
| LL282 | HK325 $\Delta dsbB$ λMalFLacZ102 pTrc99a-6XHis-*MtVKOR*<sup>-W146Y</sup> | This study |
| LL450 | HK325 $\Delta dsbB$ λMalFLacZ102 pTrc99a-6XHis-*MtVKOR*<sup>-W146R</sup> | This study |
| LL414 | HK325 $\Delta dsbB$ λMalFLacZ102 pTrc99a-6XHis-*MtVKOR*<sup>-W146F</sup> | This study |
| LL236 | HK295 $\Delta dsbB$ λMalFLacZ102 YidC<sub>T362I</sub> HslVC<sub>160Y</sub> pDSW204-6XHis-*HsVKOR*<sub>ΔA31AR D36N R37E</sub> | This study |
| LL451 | HK295 $\Delta dsbB$ λMalFLacZ102 YidC<sub>T362I</sub> HslVC<sub>160Y</sub> pDSW204-6XHis-*HsVKOR*<sub>ΔA31AR D36N R37E Y139W</sub> | This study |
| LL392 | HK295 $\Delta dsbB$ λMalFLacZ102 YidC<sub>T362I</sub> HslVC<sub>160Y</sub> pDSW204-6XHis-*HsVKOR*<sub>ΔA31AR D36N R37E Y139R</sub> | This study |
| LL413 | HK295 $\Delta dsbB$ λMalFLacZ102 YidC<sub>T362I</sub> HslVC<sub>160Y</sub> pDSW204-6XHis-*HsVKOR*<sub>ΔA31AR D36N R37E Y139F</sub> | This study |
| Plasmids |  |  |
| pTrc99a | Expression vector, Amp<sup>r</sup> | 47 |
| pDSW204 | Promoter down mutation in –35 of pTrc99a, Amp<sup>r</sup> | 33 |
| pDSW206 | Promoter down mutation in –10 and –35 of pTrc99a, Amp<sup>r</sup> | 33 |
| pRD33 | pTrc99a-6XHis-*MtVKOR*, Amp<sup>r</sup> | 12 |
| pLEM6 | pDWS206-*PaDsbB1* | 28 |
| pCL23 | pDWS204-*EcDsbB* | 48 |
| PL174 | pDSW204-6XHis-*HsVKOR*<sub>ΔA31AR</sub> | This study |
| PL101 | pDSW204-6XHis-*HsVKOR*<sub>ΔA31AR W10R D36N R37E</sub> | This study |
| PL157 | pDSW204-6XHis-*HsVKOR*<sub>ΔA31AR W10R</sub> | This study |
| PL162 | pDSW204-6XHis-*HsVKOR*<sub>ΔA31AR D36N R37E</sub> | This study |
| PL228 | pDSW204-6XHis-*HsVKOR*<sub>ΔR33 ΔR37</sub> | This study |
| PL216 | pDSW204-6XHis-*HsVKOR*<sub>ΔR33 ΔR37 W10R</sub> | This study |
| PL267 | pDSW204-6XHis-*HsVKOR*<sub>ΔR33 ΔD36 ΔR37</sub> | This study |
| PL273 | pDSW204-6XHis-*HsVKOR*<sub>R33G R37G</sub> | This study |
| PL269 | pDSW204-6XHis-*HsVKOR*<sub>ΔR33 R37N</sub> | This study |
| PL245 | pDSW204-6XHis-*HsVKOR*<sub>ΔA31AR D36N R37E Y139R</sub> | This study |
| PL262 | pDSW204-6XHis-*HsVKOR*<sub>ΔA31AR D36N R37E Y139F</sub> | This study |
| PL285 | pDSW204-6XHis-*HsVKOR*<sub>ΔA31AR D36N R37E Y139W</sub> | This study |
| PL254 | pDWS206-*EcDsbB* | This study |
| PL255 | pDWS206-*EcDsbB*<sub>R48Y</sub> | This study |
| PL207 | pDWS206-*PaDsbB1*-6XHis | This study |
| PL226 | pDWS206-*PaDsbB1*<sub>R47Y</sub>-6XHis | This study |
| PL246 | pDWS206-*PaDsbB1*<sub>S147N</sub>-6XHis | This study |
| PL197 | pTrc99a-6XHis-*MtVKOR*<sup>-W146Y</sup> | This study |
| PL179 | pTrc99a-6XHis-*MtVKOR*<sup>-N81D</sup> | This study |
| PL180 | pTrc99a-6XHis-*MtVKOR*<sup>-N81Y</sup> | This study |
| PL176 | pTrc99a-6XHis-*MtVKOR*<sup>-N81I</sup> | This study |
| PL177 | pTrc99a-6XHis-*MtVKOR*<sup>-N81S</sup> | This study |
| PL263 | pTrc99a-6XHis-*MtVKOR*<sup>-W146F</sup> | This study |
| PL284 | pTrc99a-6XHis-*MtVKOR*<sup>-W146R</sup> | This study |

**Table 2.**
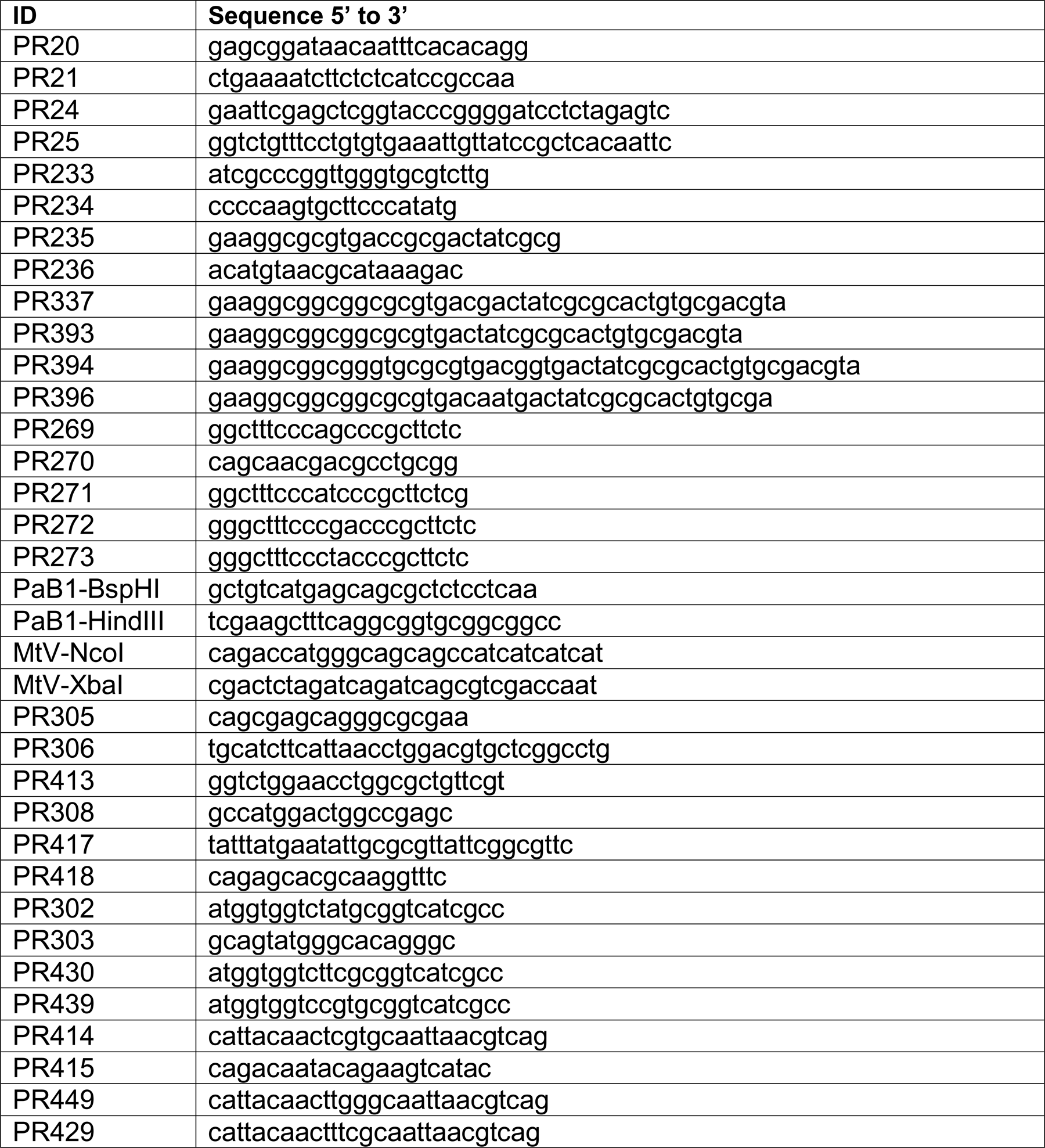
List of primers.

To clone 6X-His tagged *Hs*VKOR_ΔA31AR W10R D36N R37E_, an *E. coli* codon-optimized gene was synthesized containing the desired mutations in human VKORc1 isoform 1 Q9BQB, (gblock17 Table 3, Integrated DNA Technologies). To generate PL101, a PCR product of pDSW204 was amplified with primers PR24-PR25 and then ligated to gblock17 using HiFi Assembly (NEBuilder, New England Biolabs). The right construct was verified by PCR with primers 20 and 21 and sequenced using primer PR21. This vector was then used to construct all the mutations in human VKORc1 used in this study. Derivatives were obtained by site-directed mutagenesis using primers and KLD mix (NEBase Changer, New England Biolabs). Site-directed mutagenesis was used to make *Hs*VKOR derivatives of PL101. First, PL157 was obtained with primers PR235-PR236 and PL162 with PR233-234 using in both cases as template PL101 plasmid. Second, PL174 was constructed with primers PR233-PR234 using as template PL157 plasmid. Third, PL216 was made with primers PR236-PR337 using as template PL101 plasmid. Fourth, PL228 was obtained with primers PR236-PR337, and PL267 with PR236-PR393 using as template PL162 plasmid. Finally, PL273 was made with primers PR236-PR394, and PL269 with primers PR236-PR396 using as template PL162 plasmid.

**Table 3.**
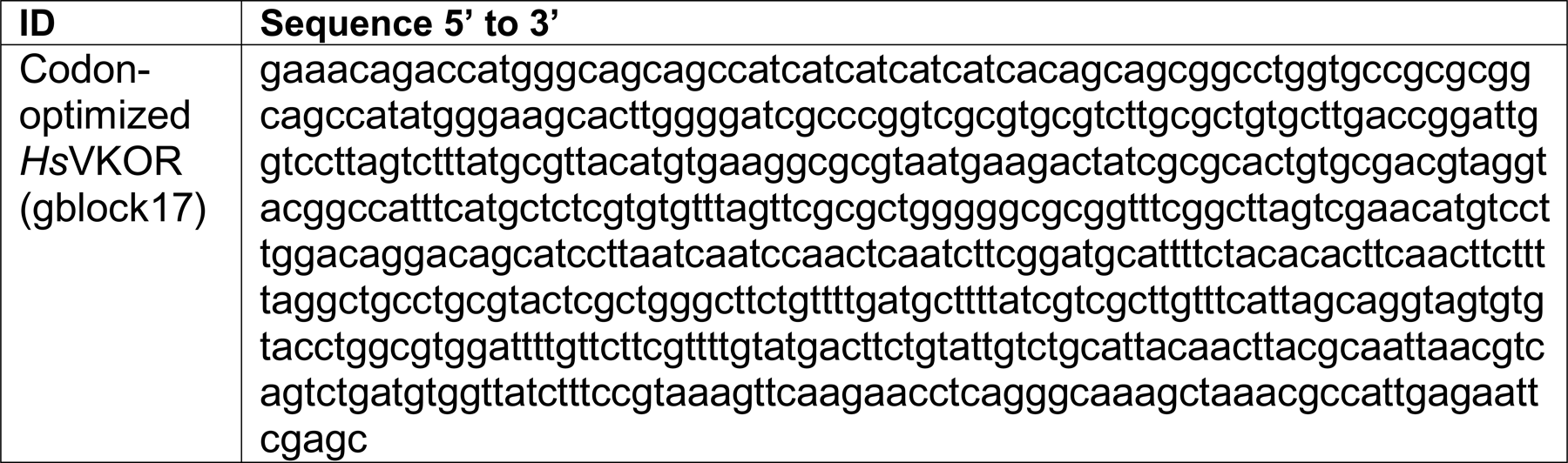
List of synthetic genes.

For the mutant library construction, a mutagenic PCR of the *PadsbB1* and *Mtvkor* genes using primer pairs PaB1-BspHI/PaB1-HindIII and MtV-NcoI/MtV-XbaI was generated using the first seven mutagenic conditions of Diversify mutagenesis kit (Clontech) that on average generates 2–5.8 mutations/kb. The amplification conditions used were 94°C (30 s) as denaturing temperature, 55 °C (30 s) as annealing, and 68 °C (30 s) as extension repeated for 25 cycles. The products were reamplified using Taq platinum (Thermo Fisher Scientific) to produce more product. PCR products of all reactions were then mixed, column-purified, digested with respective restriction enzymes, and ligated to a digested pDSW206 (*PadsbB1*) or pTrc99a (*Mtvkor*) plasmid. 1 μL of the ligation reaction was transformed into highly competent XL1-Blue cells (Agilent Technologies).

A sample of the colonies obtained after selection on ampicillin plates was collected for plasmid preparation used to confirm the efficiency of ligation by PCR and digestion. Given that 9 of 10 colonies did have the expected insert, the rest of the ligation reaction was transformed into DH10β highly competent cells (New England Biolabs). The transformation yielded 50-70,000 colonies, which were scraped up and resuspended in M63 glucose for plasmid preparation. Plasmid preparations were frozen at 20 °C until use. Plasmids were then transformed into HK325 electrocompetent cells to generate libraries of approximately the same number of colonies as the plasmid library. Colonies were then scrapped and resuspended in NZ media with 25% glycerol. Aliquots were stored at -80 °C until use. A frozen aliquot was then thawed and plated on selective M63 0.2% glucose media supplemented with antibiotics, 120 μg/mL of X-Gal, and either 2.5 μM of IPTG/10 μM Bromindione for *Mtvkor* or no IPTG/8 μM of PA5 for *PadsbB1*. The *Mt*VKOR mutant library yielded 33 white colonies. Plasmids were purified and retransformed into HK325 to confirm resistance to Bromindione. Plasmids were then sequenced with primer PR21 which gave the gene and promoter sequence. Point mutations N81S, N81I, N81D, and N81Y were introduced with PR270 in combination with either PR269, PR271, PR272, or PR273, respectively into a fresh backbone (pRD33) to confirm resistance to Bromindione. The *Pa*DsbB1 mutant library yielded only mutations mapping to the promoter region or *lacI* gene likely increasing the expression of *PadsbB1*.

The CXXC+4 mutations were generated by site-directed mutagenesis (KLD enzyme mix, New England Biolabs) on the plasmids carrying wildtype genes using the following primers: PR305-PR306 (*Pa*DsbB1_R47Y_), PR413-PR308 (*Pa*DsbB1_S147N_), PR417-PR418 (*Ec*DsbB_R48Y_), PR302-PR303 (*Mt*VKOR_W146Y_), PR439-PR303 (*Mt*VKOR_W146R_), PR430-PR303 (*Mt*VKOR_W146F_), PR449-PR415 (*Hs*VKOR_Y139W_), PR414-PR415 (*Hs*VKOR_Y139R_), and PR429-PR415 (*Hs*VKOR_Y139F_).

All *E. coli* strains were grown in NZ or M63 0.2% glucose either broth or agar media at 30-37°C when indicated. The antibiotic concentrations used were 100 μg/mL carbenicillin and 40 μg/mL kanamycin.

### Compound synthesis and resupply

Bromindione (sc-396742) was purchased from ChemCruz (USA). Compound 12 (EN300-173996, purity 95%), was purchased from Enamine (Ukraine). Compounds PA4 (JFD02470SC) and PA5 (RJC03767SC) were purchased from Maybridge, Ltd (UK). Compound MT8 (5162526) was purchased from Hit2Lead (USA). Compounds W1-6 were designed by Cristina Landeta and synthesized by Bioduro-Sundia (USA-China, see supplementary material for the synthesis methods). Compounds W7-10 were designed and synthesized by Lifan Zeng (IU Chemical Genomics, see supplementary material for the synthesis methods and spectra). Compounds W11 (STK925033), W12 (STK368402), W15 (STL372192), W16 (STL372635), W17 (STL457156) were purchased through MolPort from Vitas-M Laboratory Ltd (USA). Compound W13 (AR-683/43306491) was purchased through MolPort with Specs (USA) and W14 (AX8101833) with Oxchem Corporation (USA). All purchased compounds were analyzed by mass spectrometry (LCMS) to verify their molecular weights and to confirm their purity (over 90%). All resupplied molecules were dissolved in DMSO.

### Agar drug testing

Drug testing was performed as previously described with slight modifications^27,28^. A liquid dispenser (BioTek) fitted with a small-bore tubing cartridge was used to dispense 50 μL aliquots of hot agar medium to 384-well tissue culture-treated plates (BD Falcon #353289). Agar medium was made with M63 medium containing 0.2% glucose and 0.9% agar, supplemented with kanamycin (40 μg/mL), carbenicillin (100 μg/mL), IPTG (2.5 μM for CL382, 25 μM for LL236, 5 μM for CL377, 150 μM IPTG for FSH250, and no IPTG addition for CL325, LL395 and LL318), and X-Gal (120 μg/mL). To prevent agar solidification in the tubing the medium was maintained in a 60 °C oven and the tubing was pre-warmed by washing with sterile hot water immediately before loading the agar medium. After the agar solidified, the plates were stored overnight in a humidified sealed container at 4 °C for no more than 2 days. Molecules were added by pipetting 1 μL of serial dilutions (DMSO as solvent) into the agar’s surface. Final concentrations ranged often from 1000 μM to 0.25 μM (final DMSO concentration: 0.6%). Then, 10 μL of diluted bacteria (OD600 of 0.05) were dispensed with a liquid dispenser (BioTek). 384-well plates were sealed with a breathable film and incubated in humidity boxes at 30°C for 24 h. Plates were then stored for 2 days at 4°C to determine the minimal concentration to induce β-Gal^dbs^ and produce a blue color as described before^27,28^.

### Motility

Swarming assays were done in M63 0.2% glucose and 0.3% agar supplemented with antibiotics. No induction with IPTG was used for LL395 and LL318, while 2.5 μM and 10 μM IPTG were used for CL382 and LL236, respectively. Bacteria were stabbed into plates and incubated at 30°C. Halos were measured after 48 h.

### β-Gal quantification

β-Gal assays were done by determining the velocity of hydrolysis of o-nitrophenyl-β-galactoside (ONPG, Sigma) in microtiter plates using a protocol previously described with slight modifications^27,48,49^. Briefly, cultures were grown in M63 0.2% glucose medium with proper antibiotics at 37°C overnight. Cultures were then diluted to an OD600 of 0.01 into fresh M63 medium containing 0.2% glucose, 0.2% maltose (freshly made), proper antibiotics, and IPTG (2.5 μM for CL382, 25 μM for LL236 and no IPTG addition for LL395, LL318, CL499 and CL379). 200 μL of diluted bacteria were transferred to a 96-well plate which was sealed with a breathable film. The growth plate was incubated for 18 h at 30°C 700 rpm in an Incu-mixer MP (Benchmark). After growth, absorbance at 600 nm was read in a Synergy H1 (BioTek) plate reader. Then, 100 μL of bacteria from the growth plate were transferred to the assay plate. Note that no cell lysis step was performed. The reaction was started by adding 100 μL of the ONPG buffer to the cells (a mixture of 8 mL of Z-buffer with 4 mL of 4 mg/mL ONPG). The absorbance at λ420nm was measured every minute for 1-2 h to follow the kinetics of ONPG hydrolysis in a Synergy H1 plate reader (BioTek). The velocity of the reaction was calculated by performing linear regression using GraphPad Prism software. The slopes were then used together with OD600 and the following constants 1.81 (CF1), 2.45 (CF2), and 3.05 (CF3) to calculate Miller Units.

### Anaerobic growth

Anaerobic growth was determined as previously described^35^. Briefly, aerobic cultures were grown in M63 0.2% glucose medium containing proper antibiotics at 37°C overnight. Absorbance at λ600nm was determined and used to calculate the inoculum to reach an OD600 of 0.01 (∼9x10^6^ CFU/mL). Bacterial cultures were then diluted into anaerobic M63 medium 0.2% glucose containing 100 mM potassium nitrate. Media was degassed by transferring into a Coy anaerobic chamber to equilibrate for at least 24 h before use. The anaerobic chamber contained 85% nitrogen, 10% hydrogen, and 5% carbon dioxide. Strains CL499, CL379, LL395, LL318, and CL382 did not require the addition of IPTG, while LL236 required 2.5 μM IPTG. Mutants in the CXXC+4 residue required more IPTG: 25 µM was used for LL353; 5 µM was used for LL396, LL414, and LL450; and 15 µM was used for LL392. Bacteria were enumerated at inoculation and after 24 h of growth at 37°C. NZ aerobic plates were used for the enumeration of CFU. For drug testing assays, overnight aerobic cultures were diluted 1:50 into M63 0.2% glucose supplemented with antibiotics and incubated at 37°C shaking at 250 rpm. When cells reached log-phase after ∼3-4h (OD600 of 0.5), cultures were then diluted 1:10 and used to inoculate anaerobic media to an OD600 of 0.0001 (∼1x10^5^ CFU/mL). Bacteria were also enumerated at inoculation and after 24 h of growth. IPTG was used to induce the expression of DsbB or VKOR proteins when grown anaerobically. Strains LL395, LL318, and CL382 did not require IPTG, while LL236 required 2.5 μM IPTG. Mutants in the CXXC+4 residue required more IPTG to grow, 25 µM was used for LL353; 5 µM for LL396, LL414 and LL450; and 15 µM for LL392.

### Western blotting and protein abundance

*E. coli* strains were grown in M63 0.2% glucose media to an OD600 of 0.5-0.8 at 37°C and shaken at 250 rpm. Cells were then precipitated with trichloroacetic acid (TCA, Sigma) and reduced with Dithiothreitol (DTT, Sigma). The total protein concentration in each sample was determined by Pierce^TM^ BCA assay (ThermoFisher Scientific) prior to the addition of DTT. A 10-50 μg-aliquot of the total protein was diluted with Laemmli loading buffer and subjected to SDS-PAGE (4-20% acrylamide, Bio-Rad). Proteins were then semi-dry transferred to a PVDF membrane (Millipore). Western blotting was used to detect DsbB or VKOR using 1:10,000 dilution of α-His antibody (Santa Cruz Biotechnology). α-His antibody specificity was evaluated using CL379 strain (no His-tagged protein) and no non-specific bands were detected. Anti-RpoA (4RA2; BioLegend) was used as a loading control. Chemiluminescent substrate (ECL, Bio-Rad) was used to detect proteins using ChemiDoc™ MP (Bio-Rad) detection system. The adjusted total band volume was determined for both His and RpoA images using ImageLab Software (Bio-Rad). The resulting arbitrary units given for each band in the His and RpoA images were then normalized to the volume obtained in the wildtype protein band. The relative band volume of His was then divided by the relative band volume of RpoA to obtain the relative protein abundance plotted in the figures of this study.

### In vivo redox states of DsbB and VKOR

*E. coli* strains were diluted to an OD600 of 0.02 in M63 0.2% glucose media supplemented with 2.5 µM (CL382 and variants), 1 mM (LL318 and variants) or 25 µM IPTG (LL236 and variants). Diluted cultures were grown at 37°C and shaken at 250 rpm to mid-log phase, approximately OD600 of 0.5-0.8. Cells were then precipitated with trichloroacetic acid (TCA), washed with acetone and resuspended in Tris-HCl pH 8 1% SDS and immediately alkylated with 12.5 mM of MalPEG-2k (α-[3-(3-Maleimido-1-oxopropyl)amino]propyl-ω-methoxy, polyoxyethylene, NOF corporation) as described previously^27^. For the reduced controls, TCA-precipitated samples were treated with 100 mM Dithiothreitol (DTT, Sigma) for 30 min at room temperature. After, reduction, one control was again TCA precipitated and alkylated with 12.5 mM MalPEG-2k. The total protein concentration in each sample was determined by Pierce^TM^ BCA assay (ThermoFisher Scientific) prior to the addition of DTT or MalPEG-2k. A 10-50 μg-aliquot of the total protein was diluted with non-reducing Laemmli loading buffer and subjected to SDS-PAGE (4-20% acrylamide, Bio-Rad), proteins were then semi-dry transferred to a PVDF membrane (Millipore). Western blotting was used to detect DsbB or VKOR using 1:10,000 dilution of α-His antibody (Santa Cruz Biotechnology). Chemiluminescent substrate (ECL, Bio-Rad) was used to detect proteins through ChemiDoc (Bio-Rad) detection system or Amersham Hyperfilm^TM^ (GE Healthcare). A picture of the film was taken with an iPhone camera on a white filter LED transilluminator. Band volume was determined using Fiji ImageJ2 v2.14.0/1.54f (USA, http://imagej.net/)

### Statistical analysis

Comparisons of the mean between mutants and their respective wildtype counterpart in Figure 4 were done using Ordinary one-way ANOVA multiple comparisons with GraphPad Prism (USA). Significant differences were indicated in graphs using GP style: p-value ≤0.0001 (****), 0.0002 (***), 0.021 (**), and 0.0332 (*). Non-significant p-values 0.1234 (ns) were indicated as absence of legend.

