## Supplementary Figures and Material for "Warfarin analogs target disulfide bond-forming enzymes and suggest a residue important for quinone and coumarin binding"

Running Title: Targeting *de novo* disulfide bond formation

Dariana Chavez[1#], Gwendolyn Nita Amarquaye[1#], Adrian Mejia-Santana[1], Dyotima[1], Kayley Ryan[1], Lifan Zeng[2], Cristina Landeta[1*]

Author's affiliations:

[1] Department of Biology. Indiana University. Bloomington, USA.

[2] Indiana University Chemical Genomics Core Facility. Department of Biochemistry and Molecular Biology, School of Medicine, Indiana University. Indianapolis, USA.

[#] Authors contributed equally to this work

**Supplementary Figure 1.** Spacing is also important in addition to removal of two positive charges for *Hs*VKOR to complement the *E. coli* Δ*dsbB* mutant. Swarming motility was done by stabbing soft minimal M63 glucose plates (0.3% agar) supplemented with 10 μM IPTG. Motility halos of variants were measured after 48 h incubation at 30ºC. Data represents average±SD of at least three independent experiments.

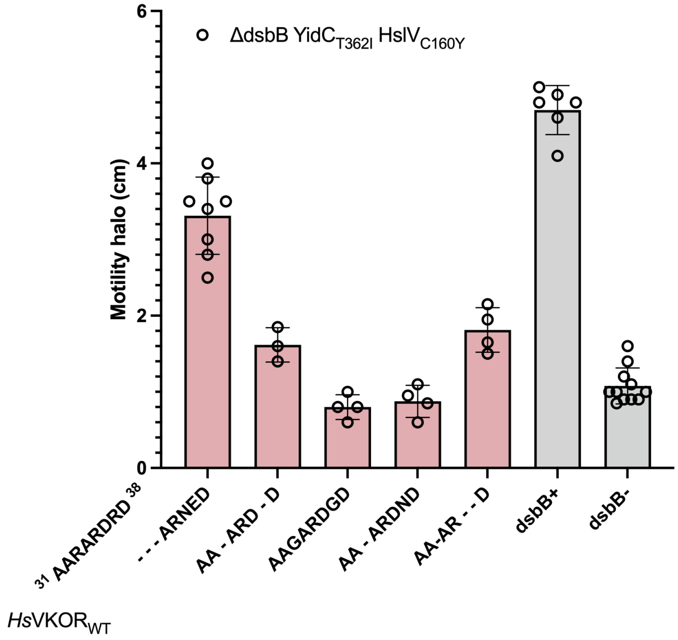

**Supplementary Figure 2.** Overexpression of *Ec*DsbB$_{R48Y}$, *Mt*VKOR$_{W146Y/F}$, *Hs*VKOR$_{Y139W}$ allows complementation of the *E. coli* Δ*dsbB* mutant. A) *E. coli* β-Gal$^{dbs}$ Δ*dsbB* strains expressing DsbB or VKOR variants were grown on X-Gal M63 0.2% glucose media supplemented with various concentrations of IPTG at 30ºC for 24 h growth. A representative image is shown. The red dotted square indicates strains that grow poorly at those concentrations of IPTG. B) Swarming motility was done by stabbing soft minimal M63 glucose plates (0.3% agar) supplemented with various concentrations of IPTG. Halos were measured after 48 h incubation at 30ºC. Data represents average±SD of at least two independent experiments. C) Anaerobic growth of *E. coli* strains expressing DsbB or VKOR variants. Strains were grown aerobically to mid-log phase in M63 0.2% glucose. Cells were then diluted to an OD600 of 0.0001 (~1x10$^5$ CFU/mL) into 200 μL of anaerobic M63 0.2% glucose medium containing 100 mM potassium nitrate, and various concentrations of IPTG. Bacteria were enumerated after 24 h of anaerobic incubation at 37ºC.

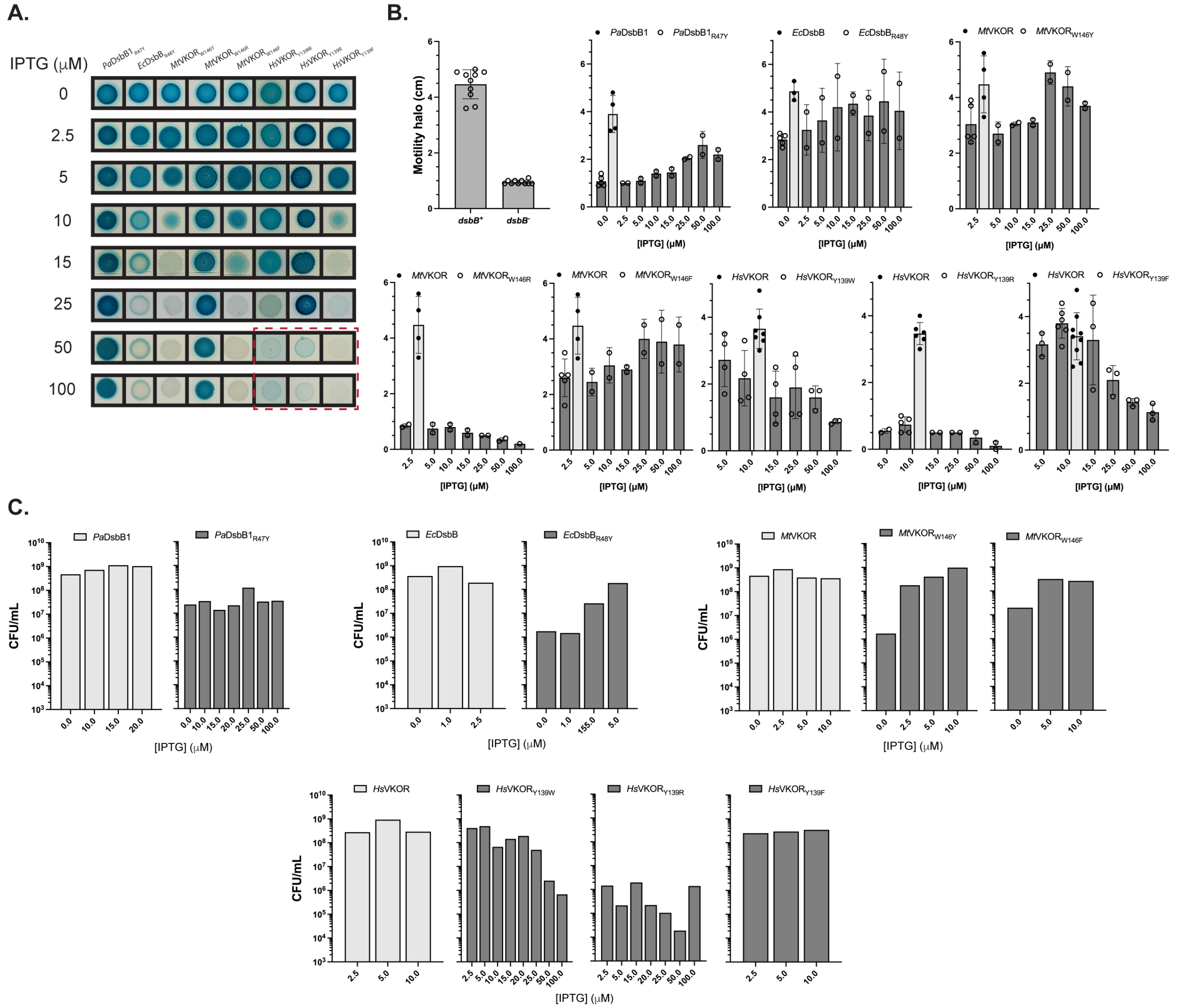

**Supplementary Figure 3.** CXXC+4 mutants are sensitized to 4-chloro-coumarin analogs with substitutions at position 3. Strains were grown aerobically to mid-log phase in M63 0.2% glucose. Cells were then diluted to an OD600 of 0.0001 (~1x$10^5$ CFU/mL, gray dotted line) into 200 µL of anaerobic M63 0.2% glucose medium containing 100 mM potassium nitrate and various concentrations of IPTG: 2.5 µM for *Mt*VKOR$_{W146Y}$, *Hs*VKOR and *Hs*VKOR$_{Y139W/F}$; 5 µM for *Ec*DsbB$_{R48Y}$ and *Mt*VKOR$_{W146R/F}$; 15 µM for *Hs*VKOR$_{Y139R}$; 25 µM for *Pa*DsbB$_{R47Y}$ and no addition of IPTG for *Ec*DsbB, *Pa*DsbB, and *Mt*VKOR. DMSO-diluted drugs were added to one final concentration indicated on the left with a 0.25% final DMSO concentration. Bacteria were enumerated after 24 h of anaerobic incubation at 37ºC. The results represent the average of at least three independent experiments.

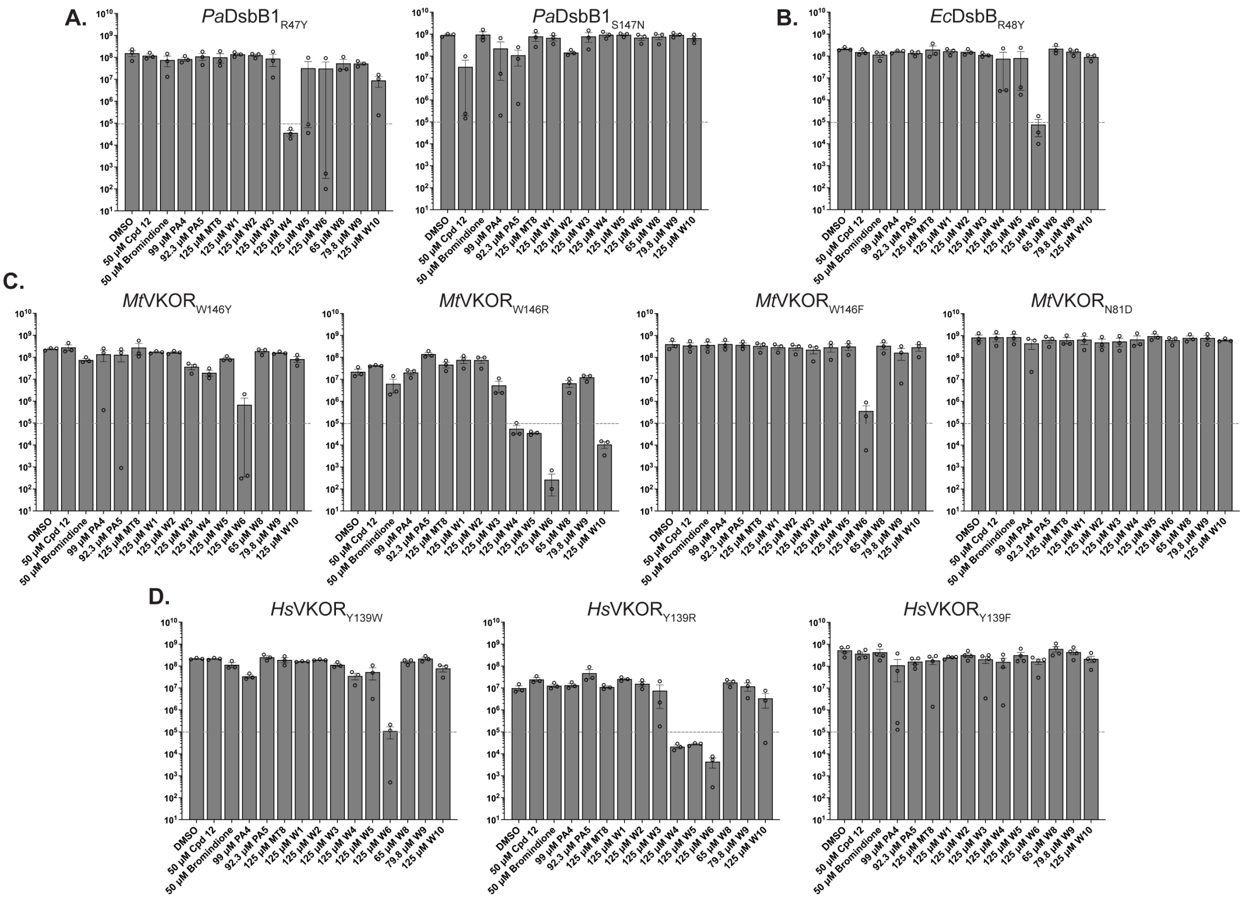

Full western blot images unedited:

Figure 1d

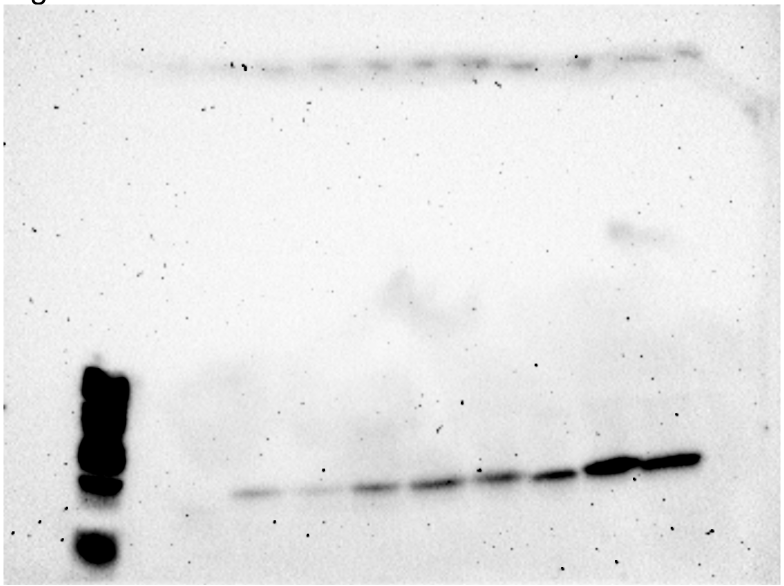

Figure 3b:

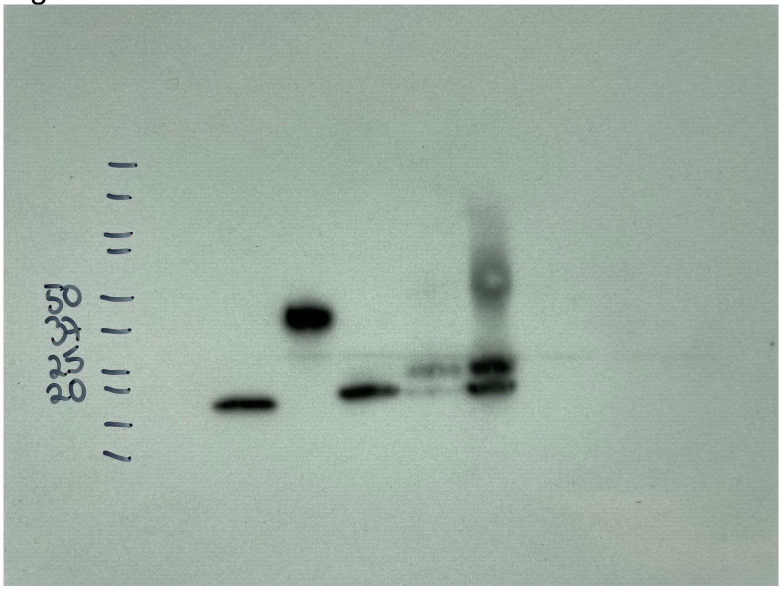

Figure 4f left:

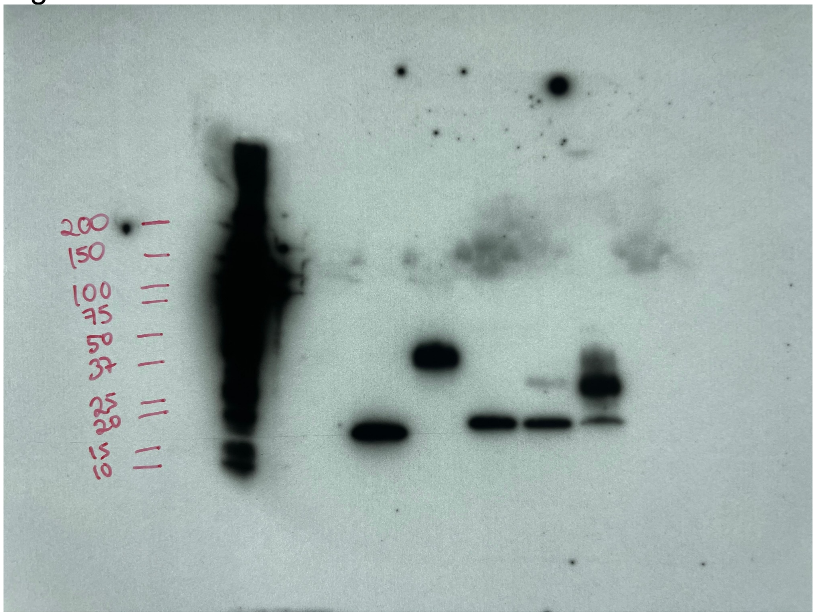


Figure 4f center:

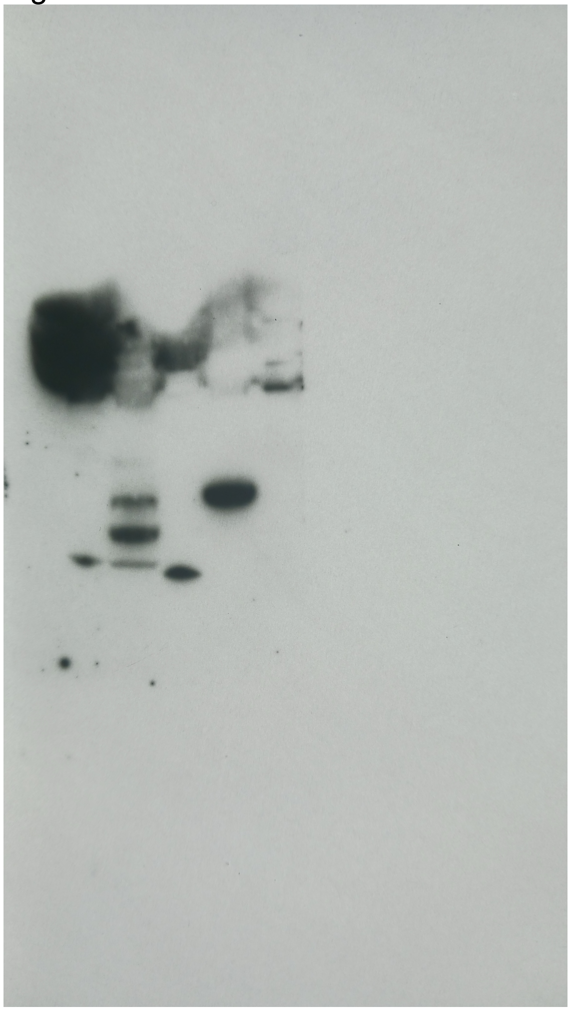

Figure 4f right:

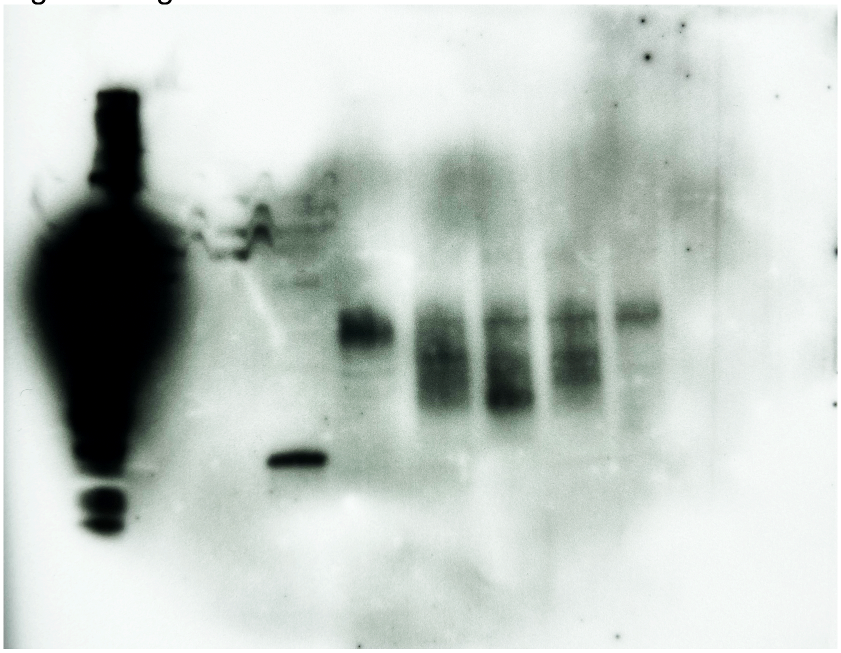

Additional supplementary information include:

1. HNMR and HPLC spectra of compounds W1 to W10

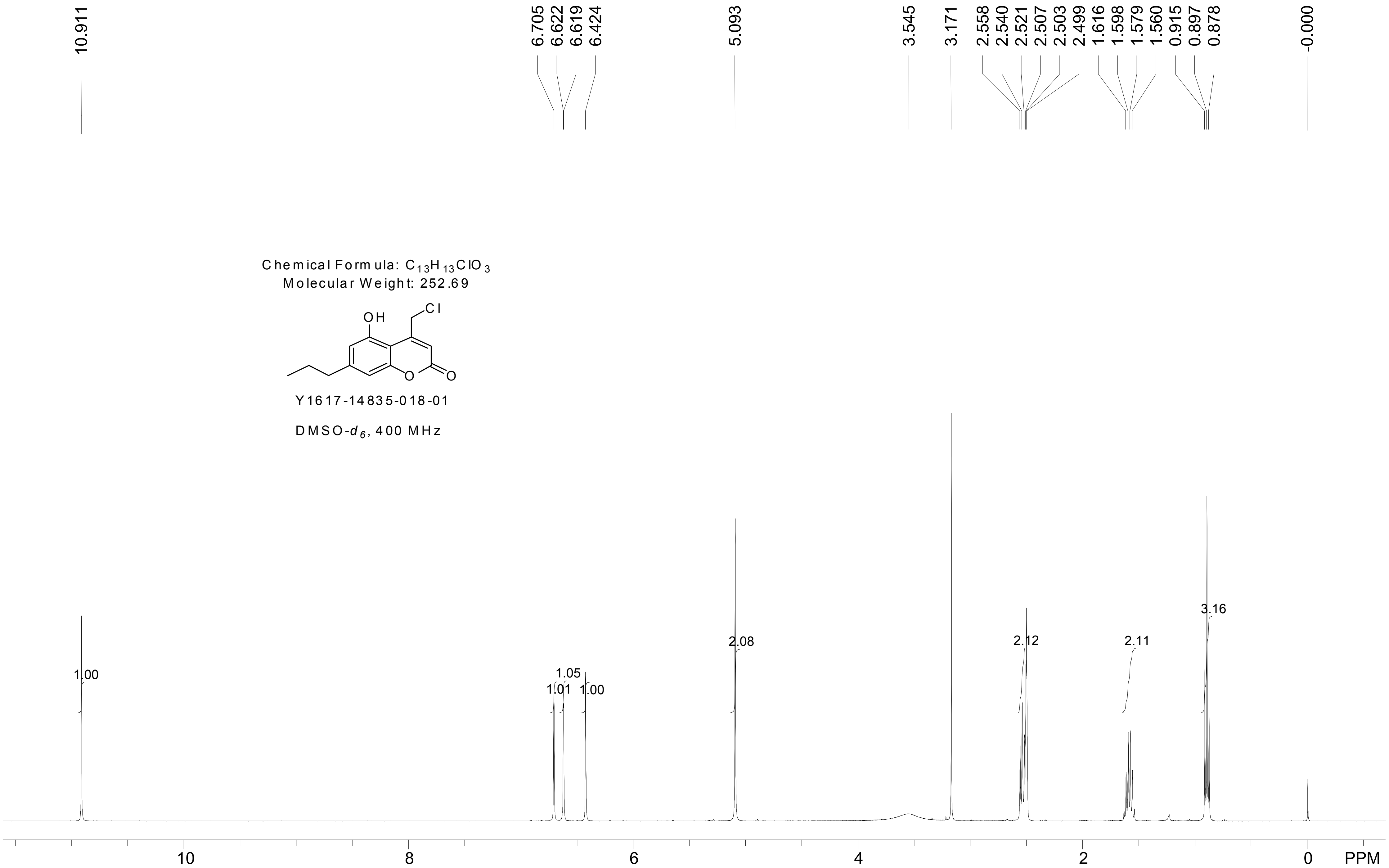

10.911
6.705
6.622
6.619
6.424
5.093
3.545
3.171
2.558
2.540
2.521
2.507
2.503
2.499
1.616
1.598
1.579
1.560
0.915
0.897
0.878
-0.000
Chemical Formula: $C_{13}H_{13}ClO_3$
Molecular Weight: 252.69
OH
Cl
O
O
Y1617-14835-018-01
DMSO-$d_6$, 400 MHz
1.00
1.01
1.05
1.00
2.08
2.12
2.11
3.16
10
8
6
4
2
0
PPM

### Sundia Meditech Co.Ltd

### QC Report

388 Jialilue Road
Zhangjiang Hightech [illegible]
Shanghai, China
Post code : 201203

```
Sample Name      : Y1617-14835-018-01
Data File        : D:\DATA-LCMS-5\2020\03-20\05\1\Y1617-14835-018-01-.D
Injection Date   : Thu, 5. Mar. 2020                Inj. Vol.   : 2ul
Acq Operator     : lcms-5                           Location    : Vial 32
Acq. Method      : D:\data-lcms-5\2020\03-20\05\1\P-20-95-TFA.M
Sample Info.     : column: ZORBAX C18 5um 4.6*50mm ;  mobile phase:B(ACN),
                   A(0.05%TFA); gradient(B%):as Acq.Method above
```

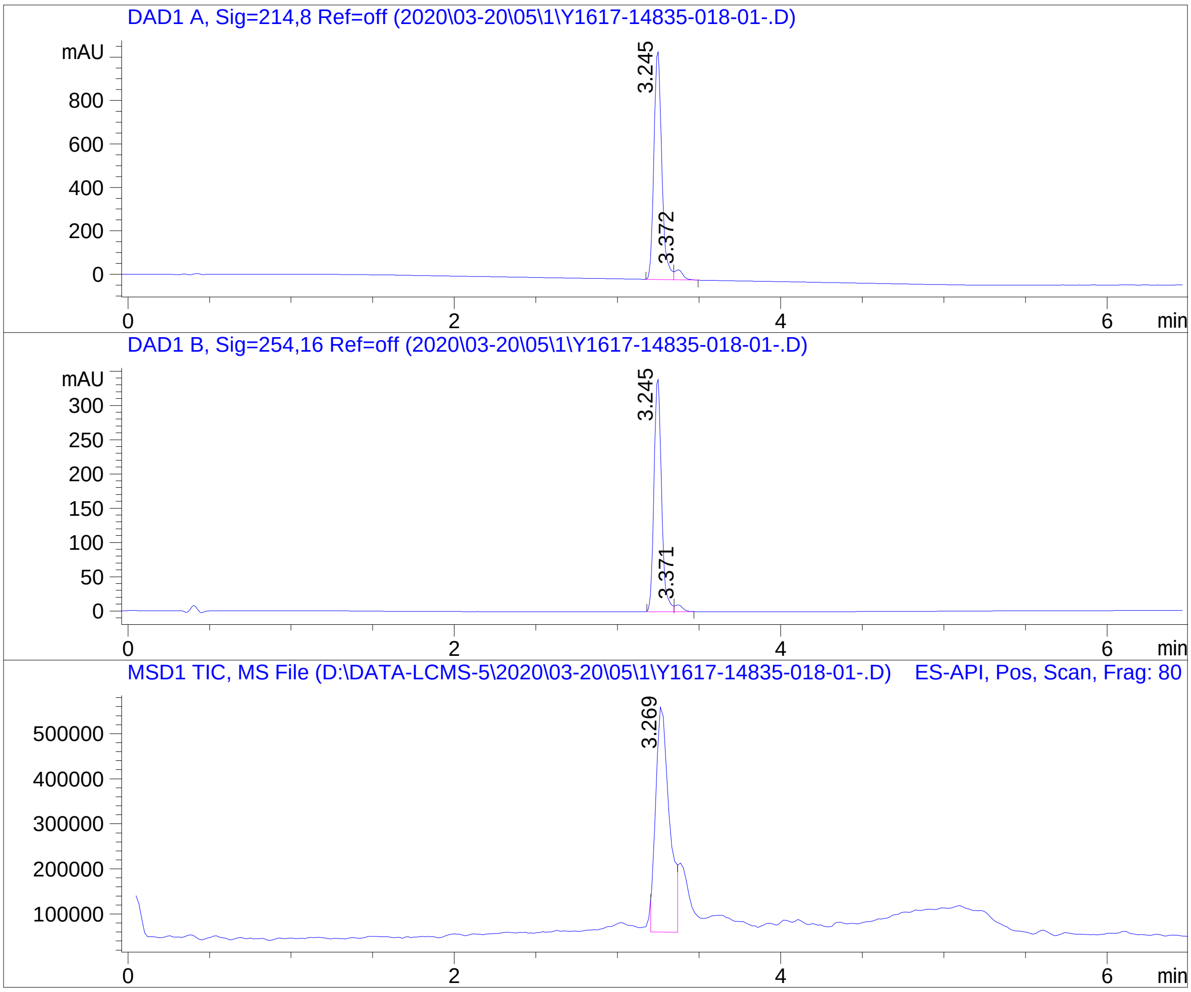


=====================================================================

Integration Results

=====================================================================

Signal 1: DAD1 A, Sig=214,8 Ref=off

| # | R.T. | Type | Height | Height% | Width | Area | Area % |
|---|---|---|---|---|---|---|---|
| 1 | 3.245 | BV | 1051.217 | 95.823 | 0.051 | 3538.250 | 95.617 |
| 2 | 3.372 | VB | 45.820 | 4.177 | 0.053 | 162.172 | 4.383 |

Signal 2: DAD1 B, Sig=254,16 Ref=off

| # | R.T. | Type | Height | Height% | Width | Area | Area % |
|---|---|---|---|---|---|---|---|
| 1 | 3.245 | BV | 340.364 | 97.212 | 0.050 | 1096.499 | 97.060 |
| 2 | 3.371 | VB | 9.763 | 2.788 | 0.052 | 33.213 | 2.940 |

Signal 3: MSD1 TIC, MS File

| # | R.T. | Type | Height | Height% | Width | Area | Area % |
|---|---|---|---|---|---|---|---|
| 1 | 3.269 | FM | 511211.750 | 100.000 | 0.095 | 2.912e+006 | 100.000 |

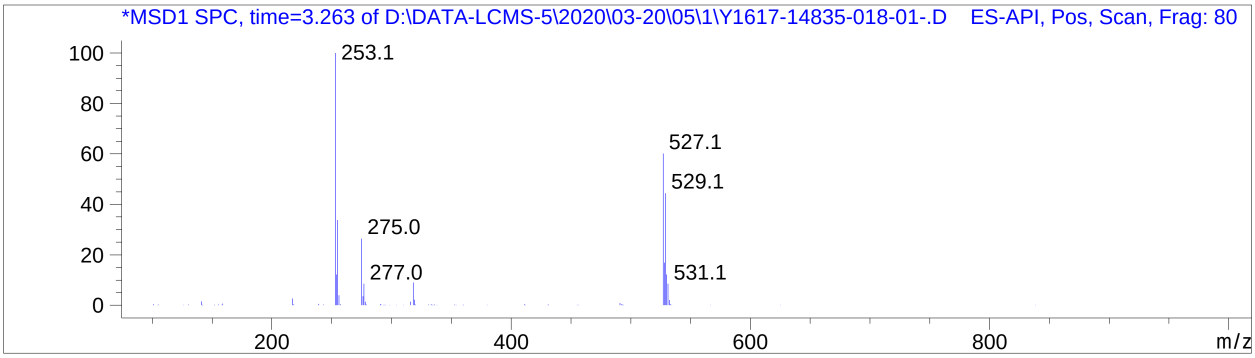


*** End of Report ***

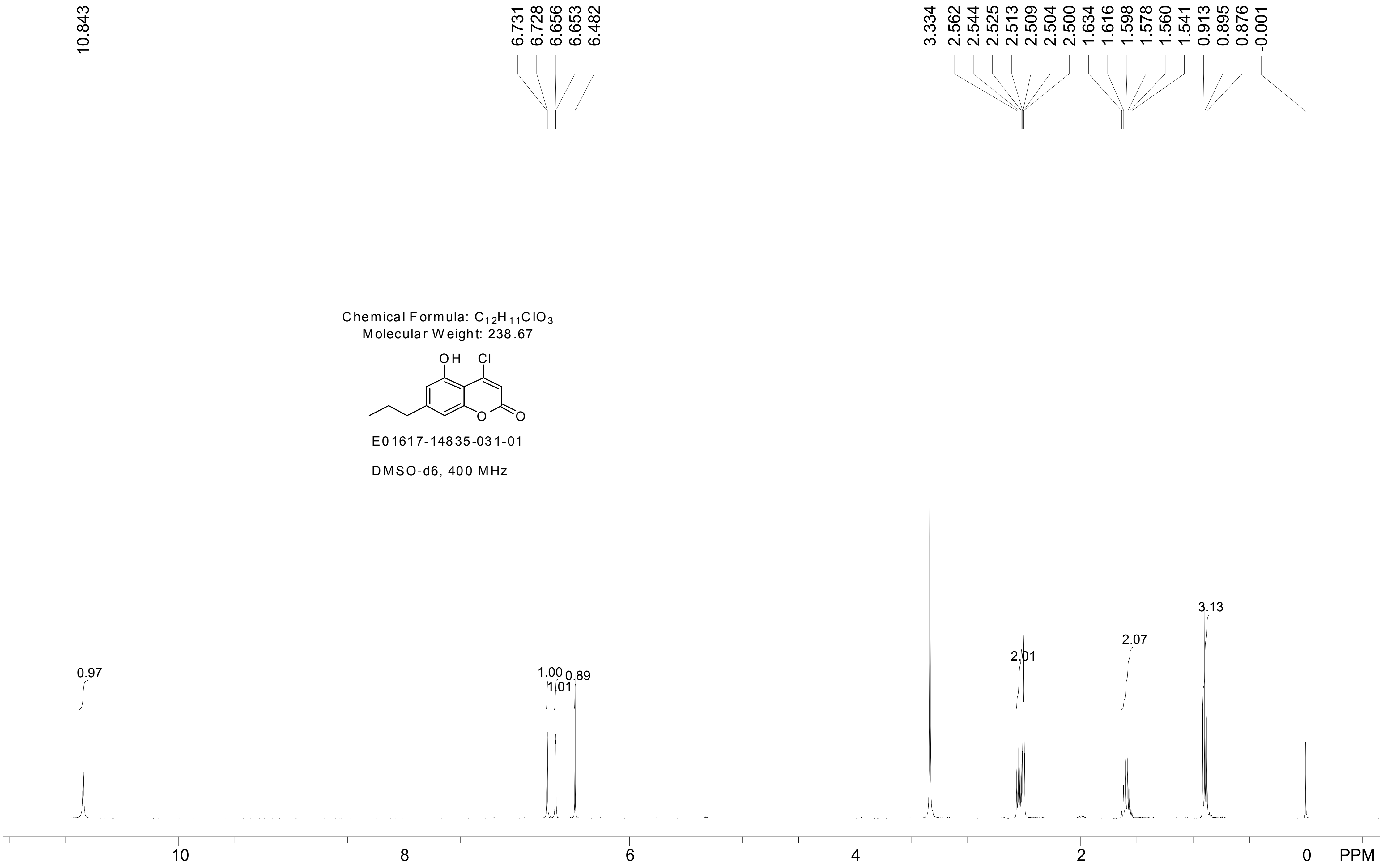

10.843
6.731
6.728
6.656
6.653
6.482
3.334
2.562
2.544
2.525
2.513
2.509
2.504
2.500
1.634
1.616
1.598
1.578
1.560
1.541
0.913
0.895
0.876
-0.001
Chemical Formula: $C_{12}H_{11}ClO_3$
Molecular Weight: 238.67
OH
Cl
O
O
E01617-14835-031-01
DMSO-d6, 400 MHz
0.97
1.00
1.01
0.89
2.01
2.07
3.13
10
8
6
4
2
0
PPM

Sundia Meditech Co.Ltd
388 Jialilue Road
Zhangjiang Hightech Park
Shanghai, China
Post code : 201203

### QC Report

Sample Name      : E01617-14835-031-01
Data File        : D:\DATA-LCMS-5\2020\06-20\15\1\E01617-14835-031-01-.D
Injection Date   : Mon, 15. Jun. 2020          Inj. Vol.   : 5ul
Acq Operator     : lcms-5                      Location    : Vial 5
Acq. Method      : D:\data-lcms-5\2020\06-20\15\1\P-30-70-TFA.M
Method Info.     : column:EC  C18 4um 4.6*50mm ;mobile phase:B(ACN), A(0.05% TFA); gradient(B%):as Acq.Method above

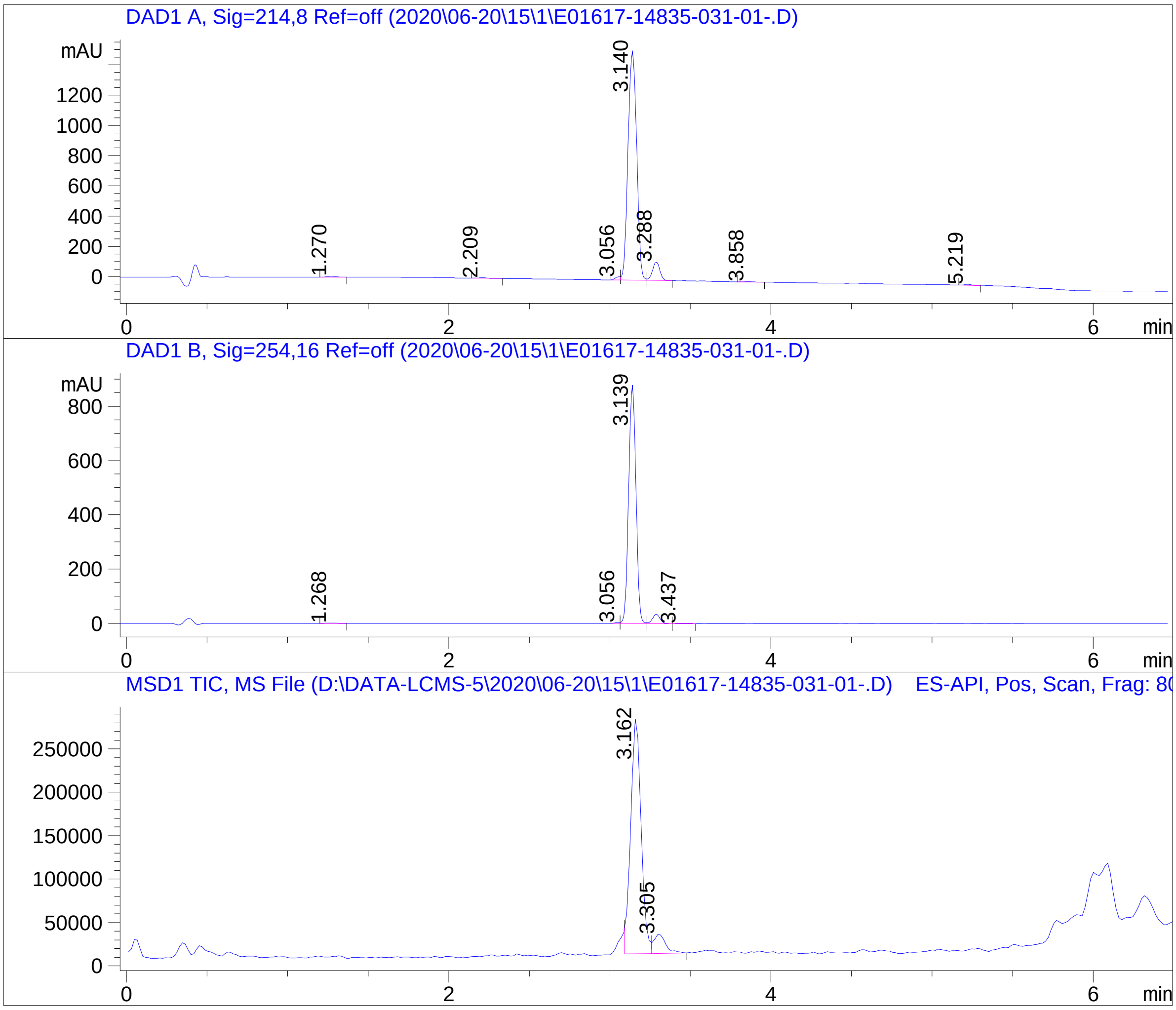


=====================================================================

Integration Results

=====================================================================

Signal  1: DAD1 A, Sig=214,8 Ref=off

| # | R.T. | Type | Height | Height% | Width | Area | Area % |
|---|---|---|---|---|---|---|---|
| 1 | 1.270 | BB | 4.814 | 0.288 | 0.053 | 16.046 | 0.261 |
| 2 | 2.209 | BB | 3.147 | 0.188 | 0.064 | 12.539 | 0.204 |
| 3 | 3.056 | MF | 20.668 | 1.235 | 0.038 | 47.627 | 0.775 |
| 4 | 3.140 | FM | 1514.989 | 90.548 | 0.062 | 5654.296 | 91.972 |
| 5 | 3.288 | VB | 121.377 | 7.254 | 0.051 | 388.959 | 6.327 |
| 6 | 3.858 | BB | 2.885 | 0.172 | 0.047 | 8.708 | 0.142 |
| 7 | 5.219 | BB | 5.261 | 0.314 | 0.056 | 19.693 | 0.320 |

| # | R.T. | Type | Height | Height% | Width | Area | Area % |
|---|---|---|---|---|---|---|---|

Signal 2: DAD1 B, Sig=254,16 Ref=off

| # | R.T. | Type | Height | Height% | Width | Area | Area % |
|---|---|---|---|---|---|---|---|
| 1 | 1.268 | BB | 1.345 | 0.146 | 0.059 | 5.214 | 0.180 |
| 2 | 3.056 | MF | 4.217 | 0.457 | 0.039 | 9.779 | 0.337 |
| 3 | 3.139 | FM | 881.458 | 95.594 | 0.052 | 2773.314 | 95.558 |
| 4 | 3.288 | VB | 34.041 | 3.692 | 0.052 | 110.031 | 3.791 |
| 5 | 3.437 | BB | 1.024 | 0.111 | 0.058 | 3.889 | 0.134 |

Signal 3: MSD1 TIC, MS File

| # | R.T. | Type | Height | Height% | Width | Area | Area % |
|---|---|---|---|---|---|---|---|
| 1 | 3.162 | FM | 276712.188 | 92.543 | 0.074 | 1.229e+006 | 91.535 |
| 2 | 3.305 | VB | 22297.127 | 7.457 | 0.085 | 113608.313 | 8.465 |

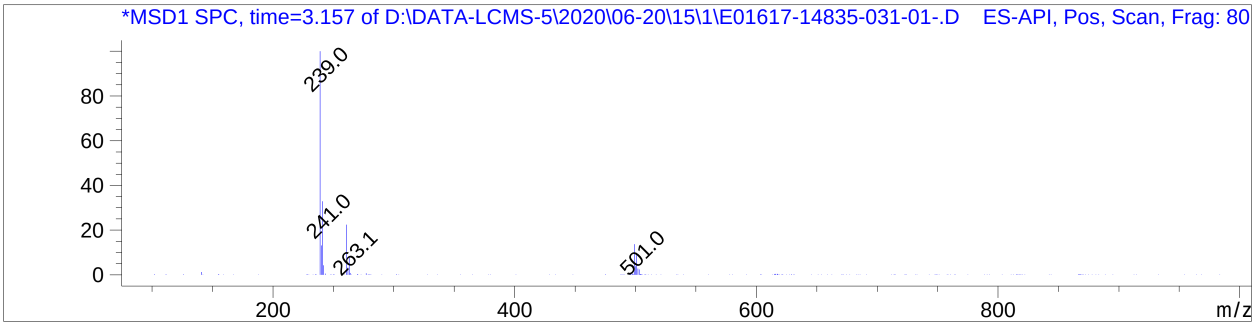


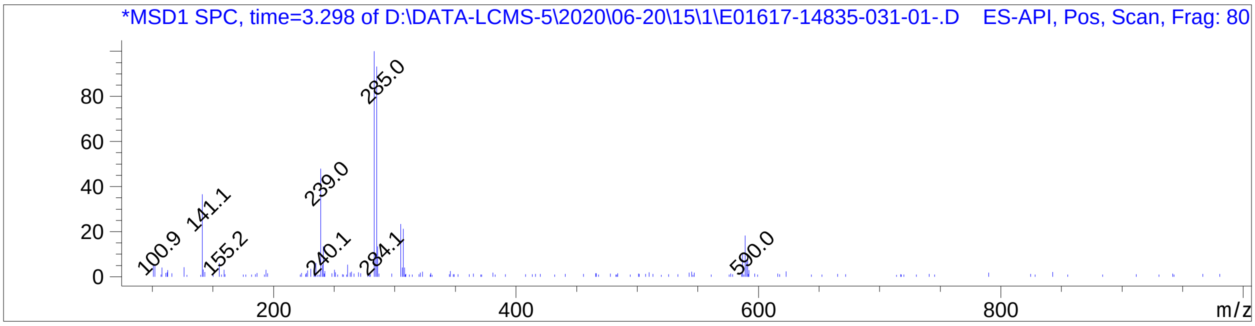


==========================================================================

*** End of Report ***

Chemical Formula: $C_{14}H_{16}O_3$
Molecular Weight: 232.28

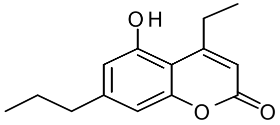


Y1617-14835-017-01

DMSO-d6, 400 MHz

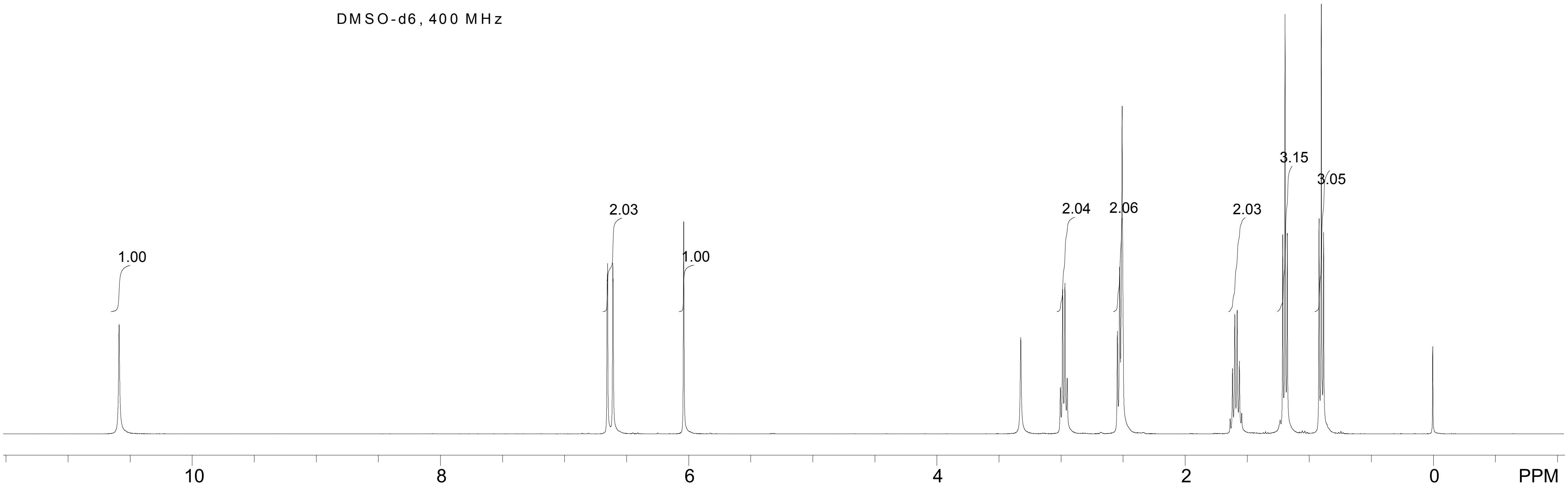

Sundia Meditech Co.Ltd

### QC Report

388 Jialilue Road
Zhangjiang Hightech Park
Shanghai, China
Post code : 201203

Sample Name : Y1617-14835-017-01
Data File : D:\DATA-LCMS-13\2020\03-20\06\2\Y1617-14835-017-01.D
Injection Date : Fri, 6. Mar. 2020 Inj. Vol. : 3ul
Acq Operator : LCMS-13 Location : Vial 91
Acq. Method : D:\data-lcms-13\2020\03-20\06\2\P-05-95.M
Method Info. : column:Ecllpse XDB-C18 5um 4.6*150mm ;mobile phase:B(ACN), A(0.02%NH4AC); gradient(B%):as Acq.Method above

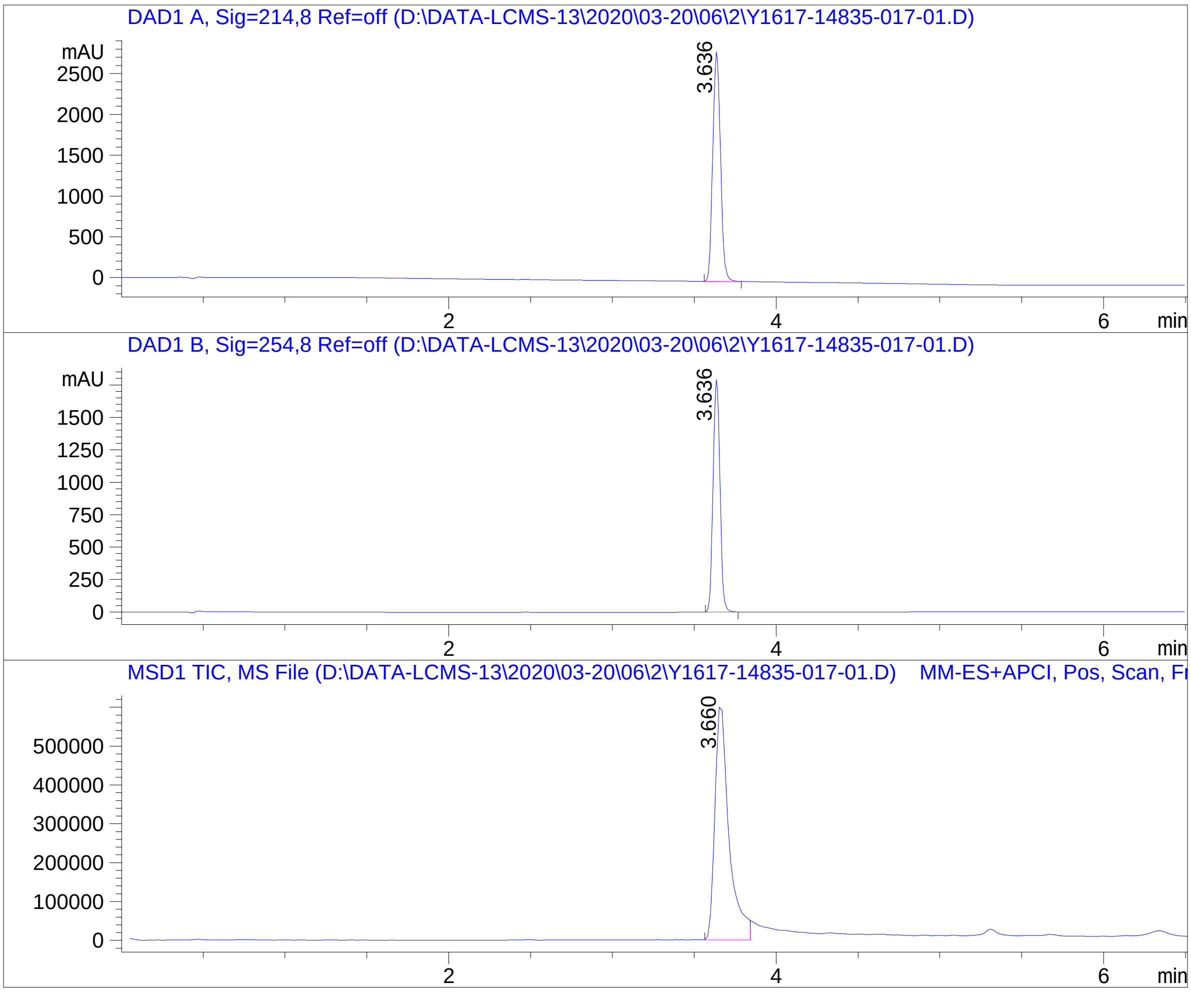


==================================================================

Integration Results

==================================================================

Signal 1: DAD1 A, Sig=214,8 Ref=off

| # | R.T. | Type | Height | Height% | Width | Area | Area % |
|---|---|---|---|---|---|---|---|
| 1 | 3.636 | BB | 2816.121 | 100.000 | 0.052 | 9264.238 | 100.000 |

Signal 2: DAD1 B, Sig=254,8 Ref=off

| # | R.T. | Type | Height | Height% | Width | Area | Area % |
|---|---|---|---|---|---|---|---|
| 1 | 3.636 | BB | 1794.671 | 100.000 | 0.046 | 5261.366 | 100.000 |

Signal 3: MSD1 TIC, MS File

| # | R.T. | Type | Height | Height% | Width | Area | Area % |
|---|---|---|---|---|---|---|---|
| 1 | 3.660 | MM | 621897.625 | 100.000 | 0.097 | 3.633e+006 | 100.000 |

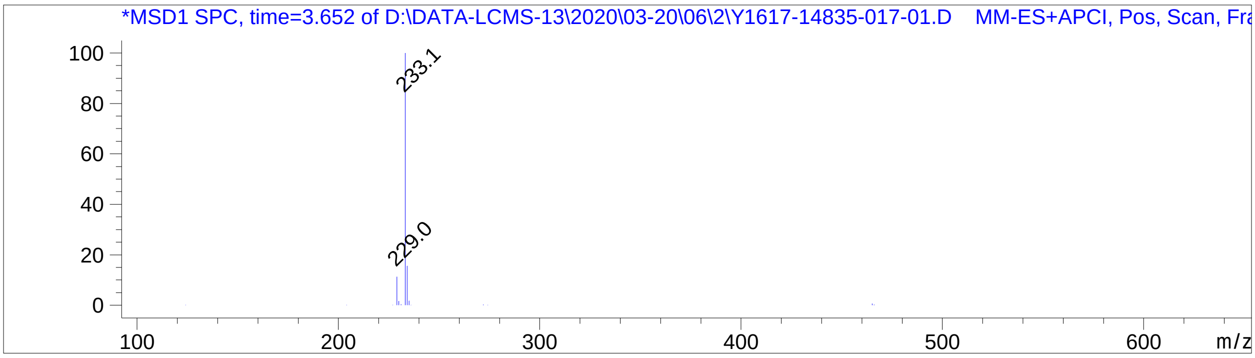


*** End of Report ***

388 Jialilue Road
Zhangjiang Hightech Par
Shanghai, China
Post code : 201203

### Sundia Meditech Co.Ltd
### QC Report

Sample Name : E03594-ENB227306-044-01
Data File : D:\DATA-LCMS5\2022\202211\25\1\E03594-ENB227306-044-01.D
Injection Date : Fri, 25. Nov. 2022 Inj. Vol. : 2ul
Acq Operator : lcms-5 Location : Vial 24
Acq. Method : D:\data-lcms5\2022\202211\25\1\C2-P-70-95-15MIN-TF
Sample Info.: column: ZORBAX SB-C8 ;column size: 4.6*150mm,3.5um ; mobile phase:B(ACN),A(0.05%TFA); gradient(B%):as Acq. Method above

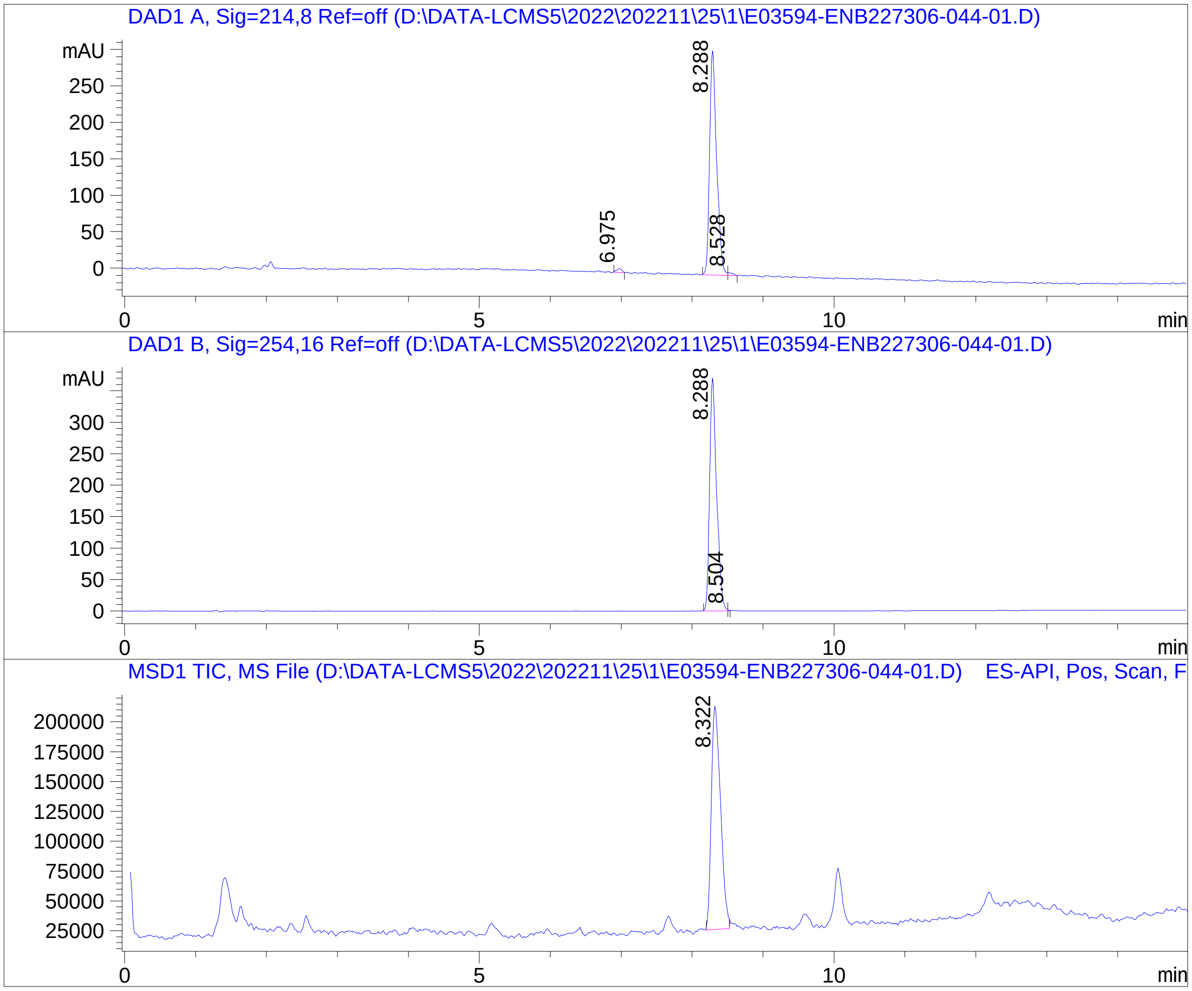


#### Integration Results

Signal 1: DAD1 A, Sig=214,8 Ref=off

| # | R.T. | Type | Height | Height% | Width | Area | Area % |
|---|---|---|---|---|---|---|---|
| 1 | 6.975 | MM | 5.041 | 1.594 | 0.083 | 25.183 | 1.167 |
| 2 | 8.288 | MF | 307.394 | 97.233 | 0.115 | 2114.688 | 98.017 |
| 3 | 8.528 | FM | 3.707 | 1.173 | 0.079 | 17.603 | 0.816 |

| # | R.T. | Type | Height | Height% | Width | Area | Area % |
|---|---|---|---|---|---|---|---|

Signal 2: DAD1 B, Sig=254,16 Ref=off

| # | R.T. | Type | Height | Height% | Width | Area | Area % |
|---|---|---|---|---|---|---|---|
| 1 | 8.288 | MF | 369.577 | 99.548 | 0.114 | 2517.349 | 99.919 |
| 2 | 8.504 | FM | 1.678 | 0.452 | 0.020 | 2.046 | 0.081 |

Signal 3: MSD1 TIC, MS File

| # | R.T. | Type | Height | Height% | Width | Area | Area % |
|---|---|---|---|---|---|---|---|
| 1 | 8.322 | MF | 188447.359 | 100.000 | 0.150 | 1.692e+006 | 100.000 |

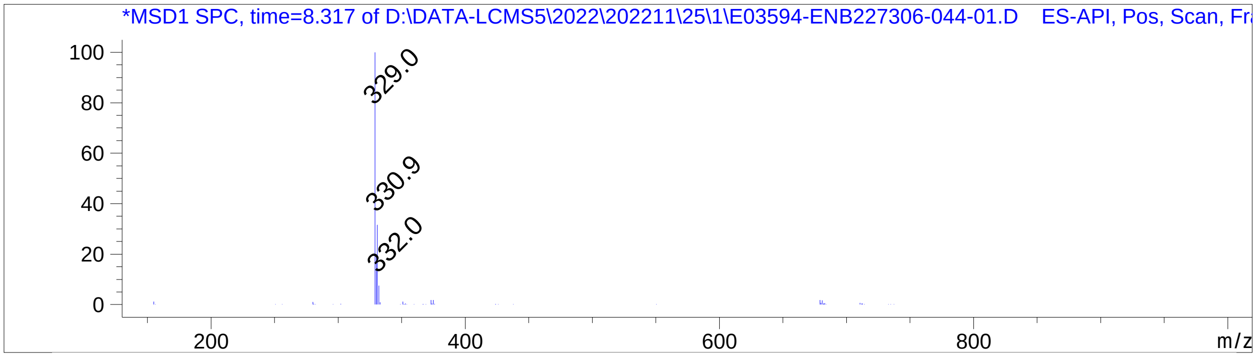


*** End of Report ***

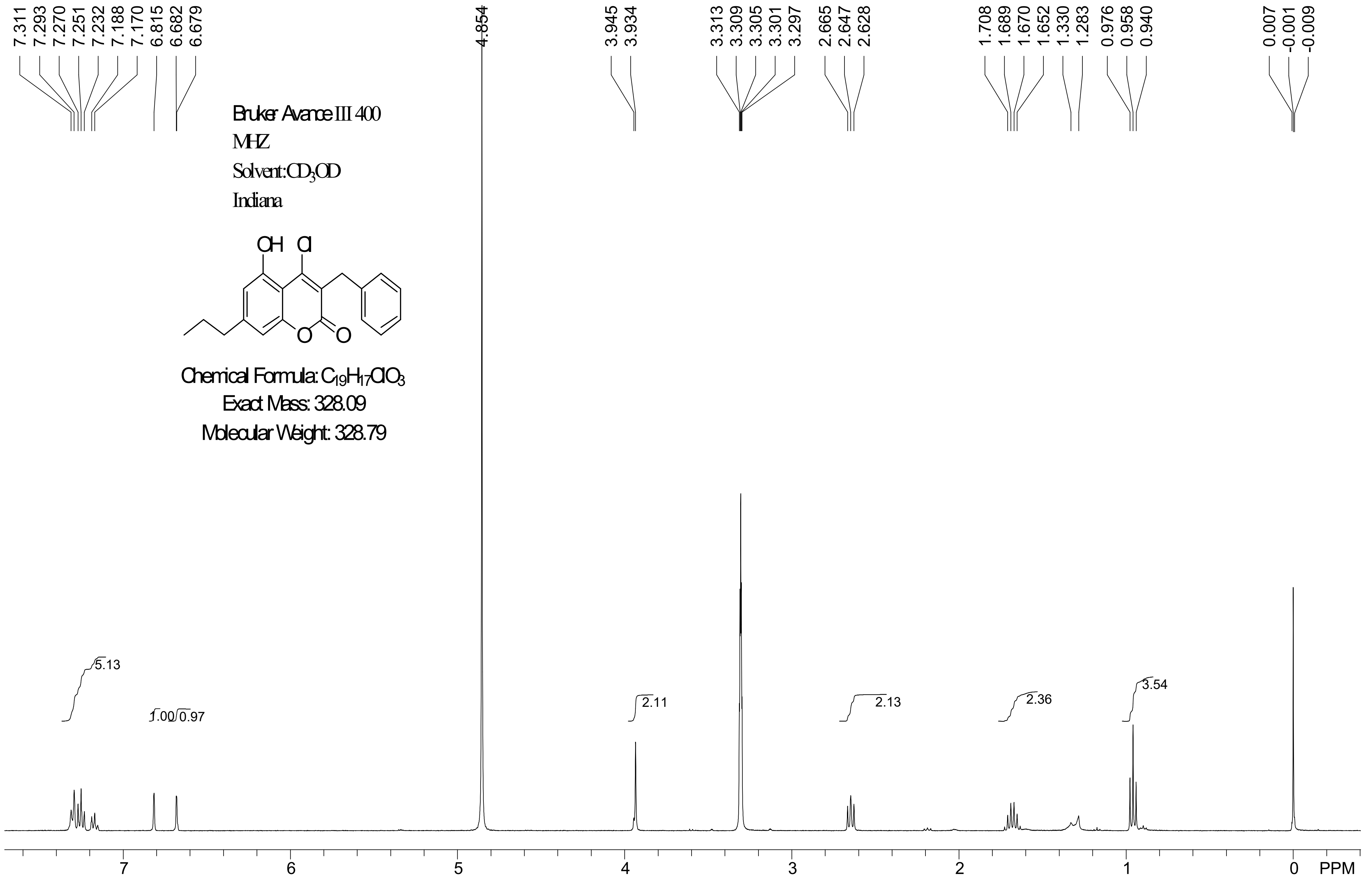
7.311 7.293 7.270 7.251 7.232 7.188 7.170 6.815 6.682 6.679
4.854
3.945 3.934
3.313 3.309 3.305 3.301 3.297 2.665 2.647 2.628
1.708 1.689 1.670 1.652 1.330 1.283 0.976 0.958 0.940
0.007 -0.001 -0.009
Bruker Avance III 400
MHZ
Solvent:CD3OD
Indiana
OH Cl
O O
Chemical Formula: C19H17ClO3
Exact Mass: 328.09
Molecular Weight: 328.79
5.13
1.00 0.97
2.11
2.13
2.36
3.54
7 6 5 4 3 2 1 0 PPM

388 Jialilue Road
Zhangjiang Hightech Park
Shanghai , China
Post code :201203

### Sundia Meditech Co.Ltd
### QC Report

| | |
|---|---|
| Sample Name | : E03594-ENB227306-060-01 |
| Data File | : D:\data-RPHPLC2\2022\202212\13\E03594-ENB227306-060-01----.lcd |
| Injection Date | : 12/13/2022 |
| Inj. Vol. | : 0.2 ul |
| Vial# | : 19 |
| Acq. Operator | : LCMS-26 |
| Acq. Method | : D:\methods\A\1.2ml\A-6.5min-70-95.lcm |
| Oven Temperature | : 40 C |
| Column | : Waters Sunfire C18 3.5um, 50*4.6mm |
| Mobile Phase A | : 0.02%NH4AC aq. |
| Mobile Phase B | : ACN |
| Total Flow | : 1.2ml/min |

Chromatogram

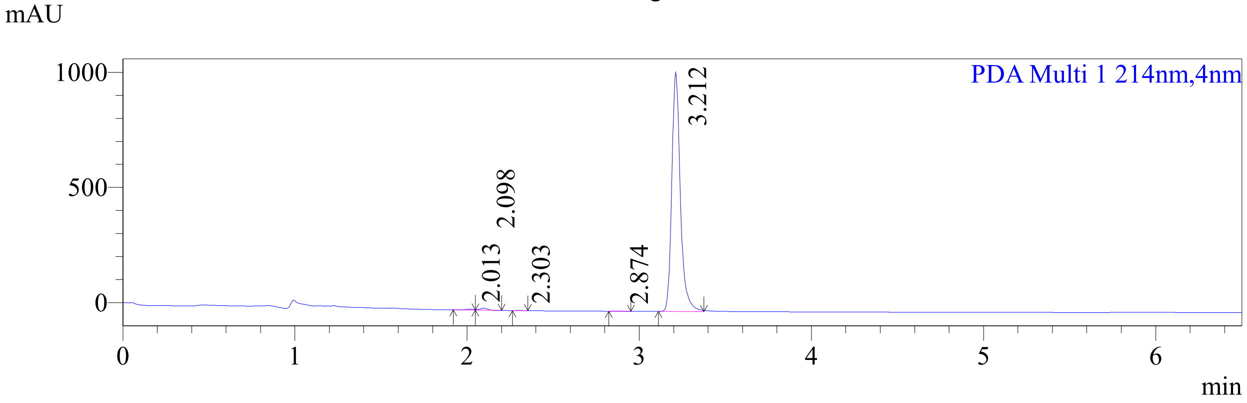


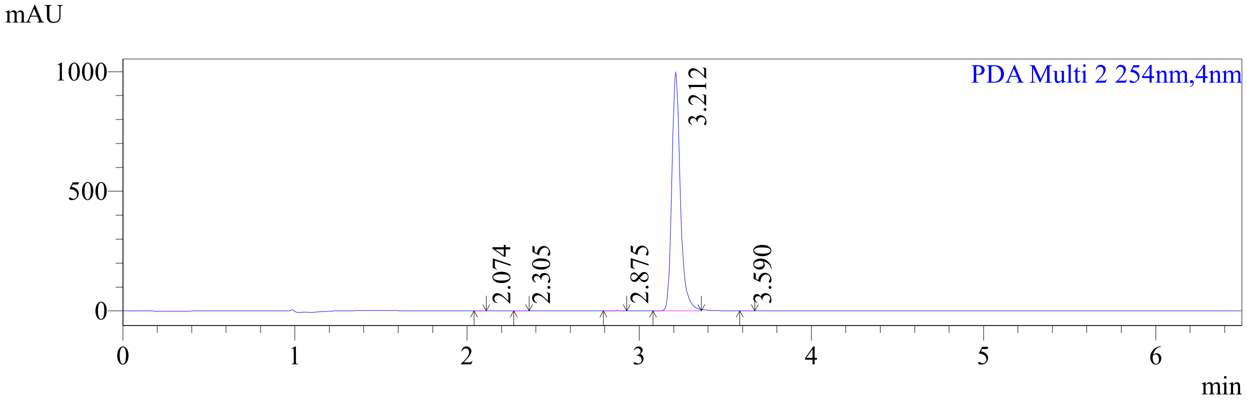


Peak Table

PDA Ch1 214nm

| # | R.T. | Mark | Height | Area | Area% | Tailing Factor | Resolution |
|---|---|---|---|---|---|---|---|
| 1 | 2.013 | | 4206.08 | 14440.14 | 0.40 | -- | -- |
| 2 | 2.098 | V | 9437.64 | 29388.09 | 0.81 | -- | 0.91 |
| 3 | 2.303 | | 1003.07 | 2392.05 | 0.07 | 1.08 | 2.57 |
| 4 | 2.874 | M | 1036.17 | 3388.98 | 0.09 | 1.39 | 7.03 |
| 5 | 3.212 | M | 1027698.02 | 3589440.28 | 98.64 | 1.28 | 3.69 |

PDA Ch2 254nm

| # | R.T. | Mark | Height | Area | Area% | Tailing Factor | Resolution |
|---|---|---|---|---|---|---|---|
| 1 | 2.074 | M | 498.30 | 1096.14 | 0.03 | 1.04 | -- |
| 2 | 2.305 | M | 566.27 | 1395.98 | 0.04 | 1.16 | 3.42 |
| 3 | 2.875 | M | 1198.17 | 3896.56 | 0.11 | -- | 7.02 |

| # | R.T. | Mark | Height | Area | Area% | Tailing Factor | Resolution |
|---|---|---|---|---|---|---|---|
| 4 | 3.212 | M | 986949.87 | 3448762.97 | 99.77 | 1.28 | 3.65 |
| 5 | 3.590 | M | 373.17 | 1531.04 | 0.04 | -- | 0.20 |

### Sundia Meditech Co.Ltd
### QC Report

388 Jialilue Road
Zhangjiang Hightech Park
Shanghai, China
Post code : 201203

| | | | | |
|---|---|---|---|---|
| Sample Name | : E03594-ENB227306-060-01 | | | |
| Data File | : D:\DATA-LCMS13\2022\202212\13\1\E03594-ENB227306-060-0-> | | | |
| Injection Date | : Tue, 13. Dec. 2022 | Inj. Vol. | : | 0.30000001ul |
| Acq Operator | : LCMS-13 | Location | : | Vial 41 |
| Acq. Method | : D:\data-lcms13\2022\202212\13\1\P-70-95-2ML-TFA.M | | | |
| Method Info. | : column:Diamonsil Plus,5um 4.6*30mm ;mobile phase:B(ACN), A(0.05%TFA aq.); gradient(B%):as Acq.Method above | | | |

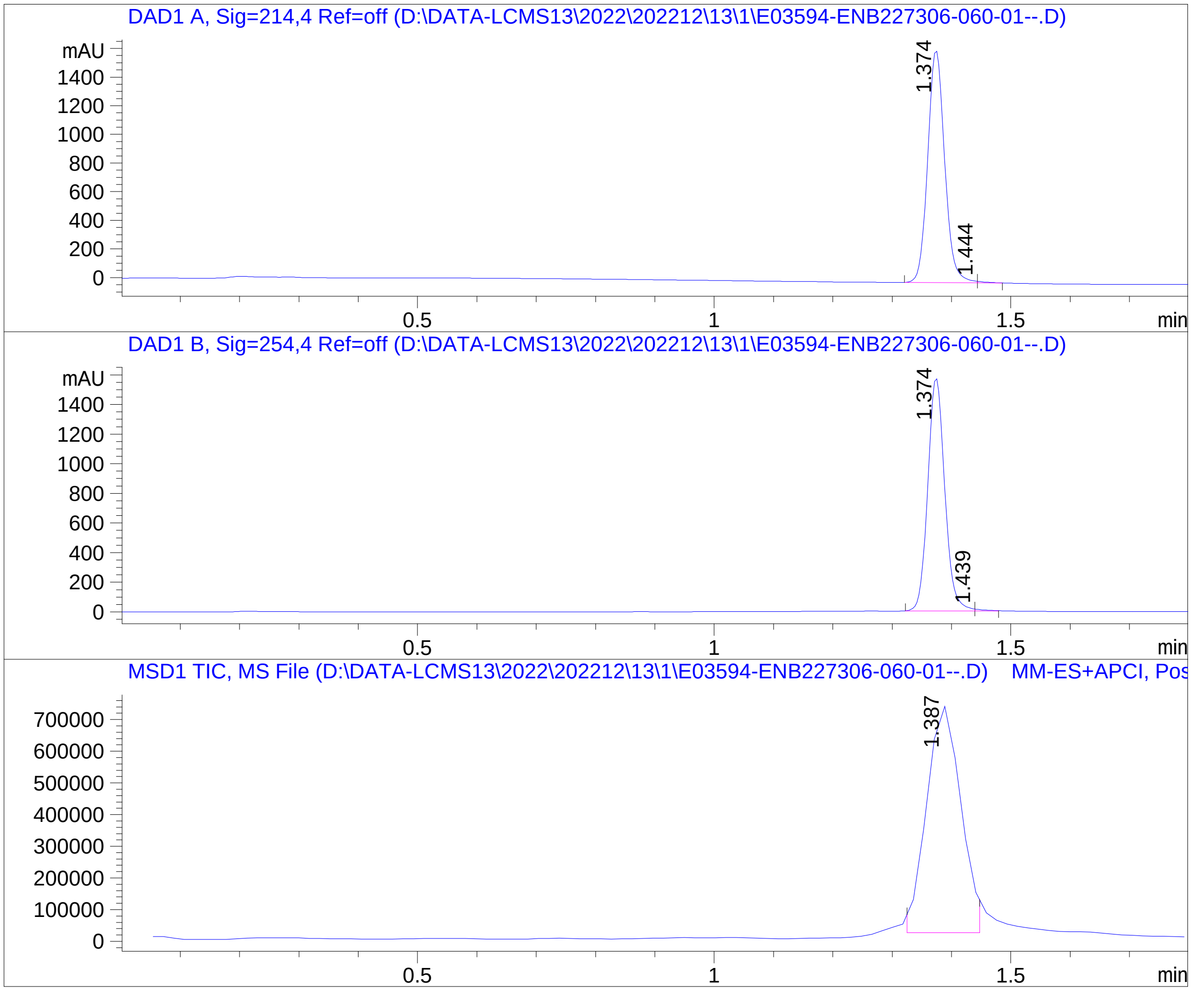


=====================================================================

Integration Results

=====================================================================

Signal 1: DAD1 A, Sig=214,4 Ref=off

| # | R.T. | Type | Height | Height% | Width | Area | Area % |
|---|---|---|---|---|---|---|---|
| 1 | 1.374 | MF | 1627.19 | 99.35 | 0.03 | 3219.91 | 99.68 |
| 2 | 1.444 | FM | 10.60 | 0.65 | 0.02 | 10.26 | 0.32 |

Signal 2: DAD1 B, Sig=254,4 Ref=off

| # | R.T. | Type | Height | Height% | Width | Area | Area % |
|---|---|---|---|---|---|---|---|
| 1 | 1.374 | MF | 1576.79 | 99.23 | 0.03 | 3126.26 | 99.61 |
| 2 | 1.439 | FM | 12.26 | 0.77 | 0.02 | 12.14 | 0.39 |

Signal 3: MSD1 TIC, MS File

| # | R.T. | Type | Height | Height% | Width | Area | Area % |
|---|---|---|---|---|---|---|---|
| 1 | 1.387 | MM | 727398.44 | 100.00 | 0.07 | 2839274.00 | 100.00 |

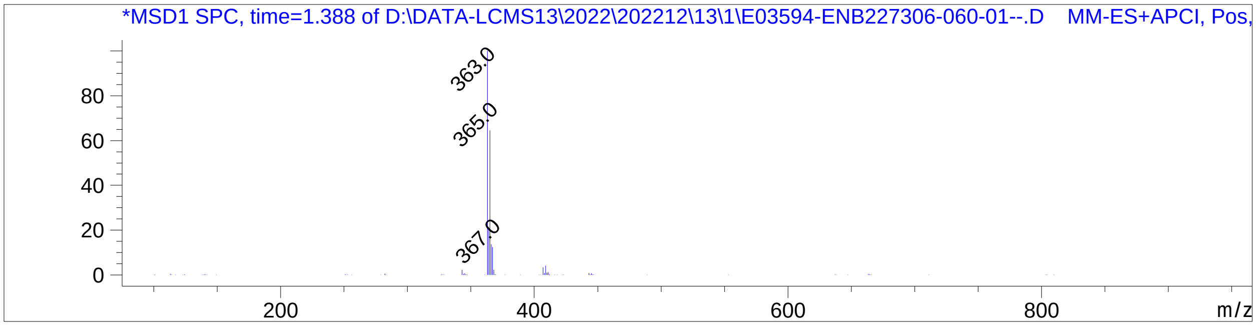


================================================================================
*** End of Report ***

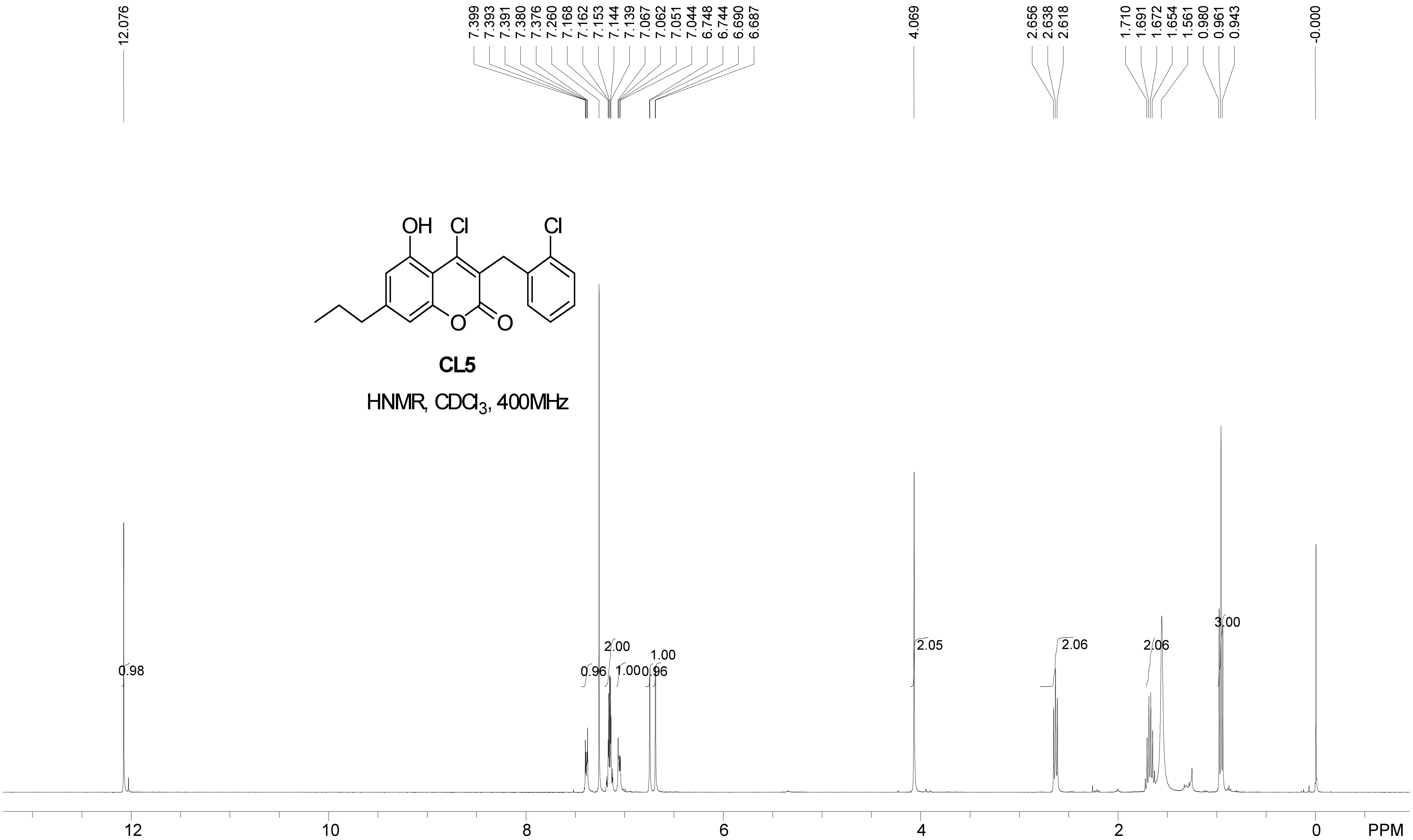
12.076
7.399 7.393 7.391 7.380 7.376 7.260 7.168 7.162 7.153 7.144 7.139 7.067 7.062 7.051 7.044 6.748 6.744 6.690 6.687
4.069
2.656 2.638 2.618
1.710 1.691 1.672 1.654 1.561 0.980 0.961 0.943
-0.000
OH Cl Cl
O O
CL5
HNMR, $CDCl_3$, 400MHz
0.98
0.96
2.00
1.00
1.00
0.96
2.05
2.06
2.06
3.00
12 10 8 6 4 2 0 PPM

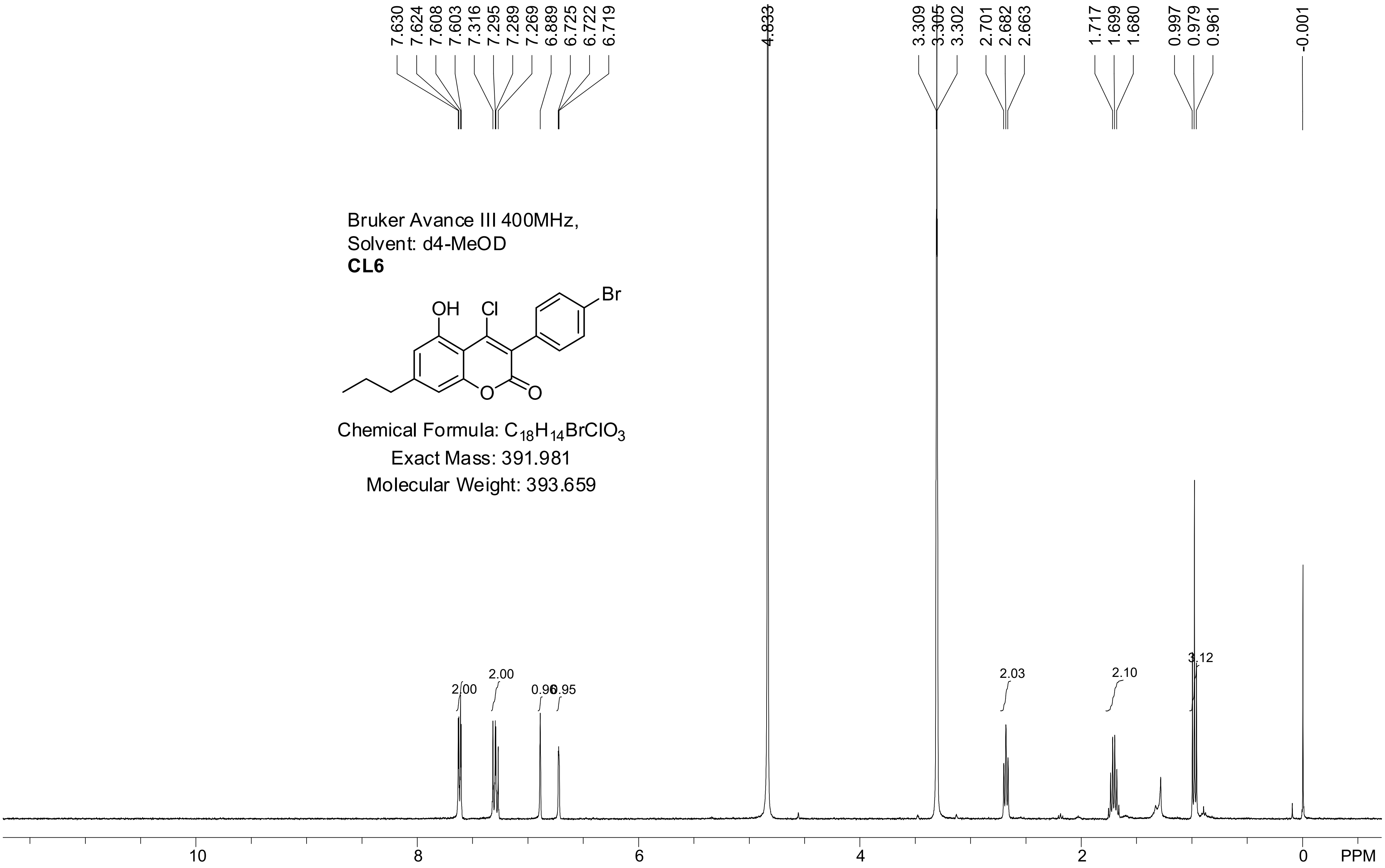


Bruker Avance III 400MHz,
Solvent: d4-MeOD
**CL6**

Chemical Formula: $C_{18}H_{14}BrClO_3$
Exact Mass: 391.981
Molecular Weight: 393.659

Sundia Meditech Co.Ltd

### QC Report

388 Jialilue Road
Zhangjiang Hightech Par
Shanghai, China
Post code : 201203

```
Sample Name      : E03722-ENB227271-047-001
Data File        : D:\DATA-LCMS5\2022\202211\25\1\E03722-ENB227271-047-00->
Injection Date   : Fri, 25. Nov. 2022            Inj. Vol.  : 2ul
Acq Operator     : lcms-5                        Location   : Vial 23
Acq. Method      : D:\data-lcms5\2022\202211\25\1\C2-P-70-95-15MIN-TF
Sample Info.:      column: ZORBAX   SB-C8  ;column  size: 4.6*150mm,3.5um ;
                   mobile phase:B(ACN),A(0.05%TFA); gradient(B%):as Acq.
                   Method above
```

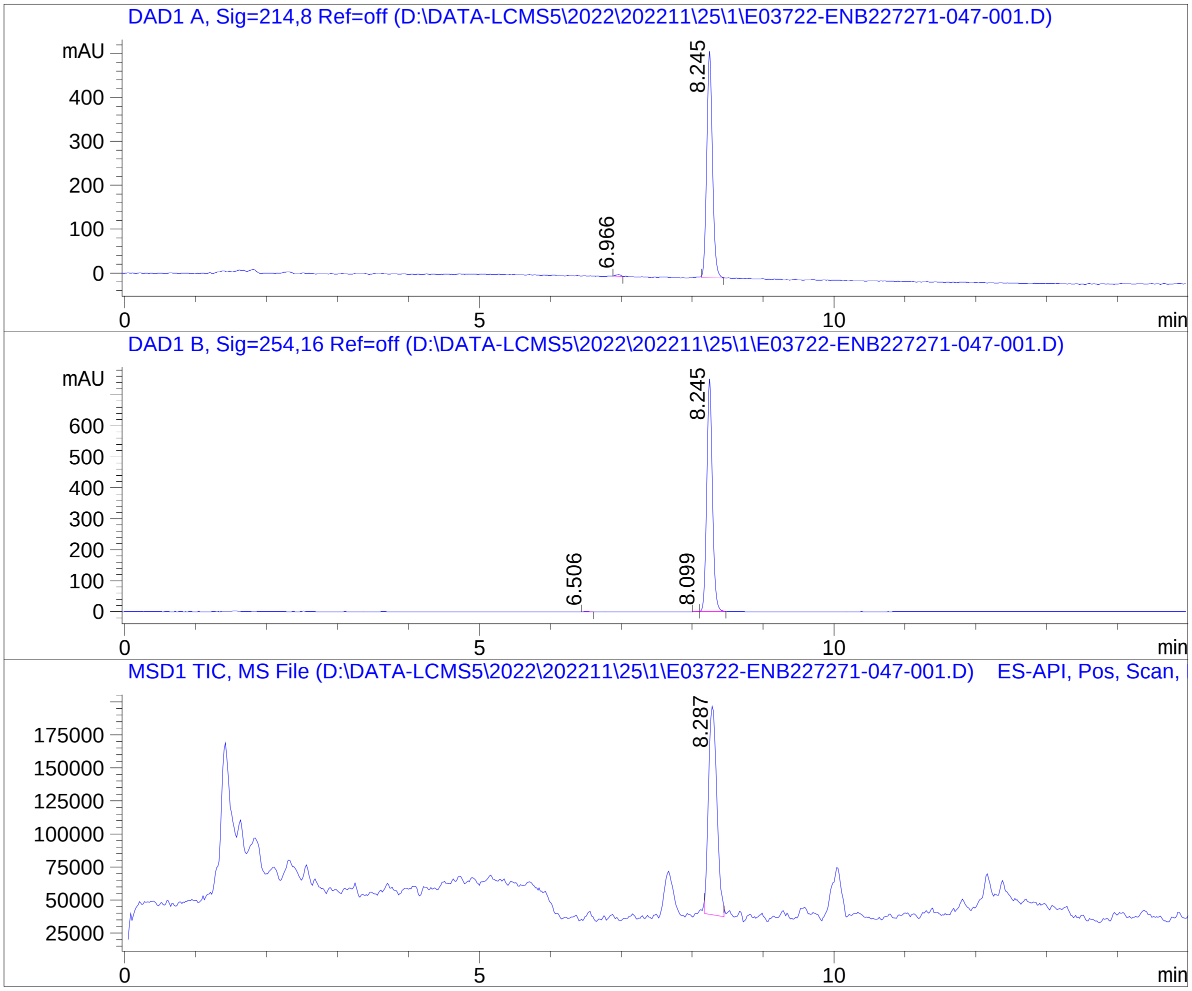


=====================================================================

Integration Results

=====================================================================

Signal 1: DAD1 A, Sig=214,8 Ref=off

| # | R.T. | Type | Height | Height% | Width | Area | Area % |
|---|---|---|---|---|---|---|---|
| 1 | 6.966 | MM | 3.710 | 0.713 | 0.081 | 17.998 | 0.614 |
| 2 | 8.245 | BB | 516.318 | 99.287 | 0.088 | 2911.518 | 99.386 |

Signal 2: DAD1 B, Sig=254,16 Ref=off

| # | R.T. | Type | Height | Height% | Width | Area | Area % |
|---|---|---|---|---|---|---|---|
| 1 | 6.506 | MM | 1.008 | 0.133 | 0.096 | 5.811 | 0.138 |
| 2 | 8.099 | MF | 1.473 | 0.195 | 0.051 | 4.527 | 0.108 |
| 3 | 8.245 | FM | 753.035 | 99.672 | 0.093 | 4192.384 | 99.754 |

Signal 3: MSD1 TIC, MS File

| # | R.T. | Type | Height | Height% | Width | Area | Area % |
|---|---|---|---|---|---|---|---|
| 1 | 8.287 | MM | 158636.406 | 100.000 | 0.137 | 1.303e+006 | 100.000 |

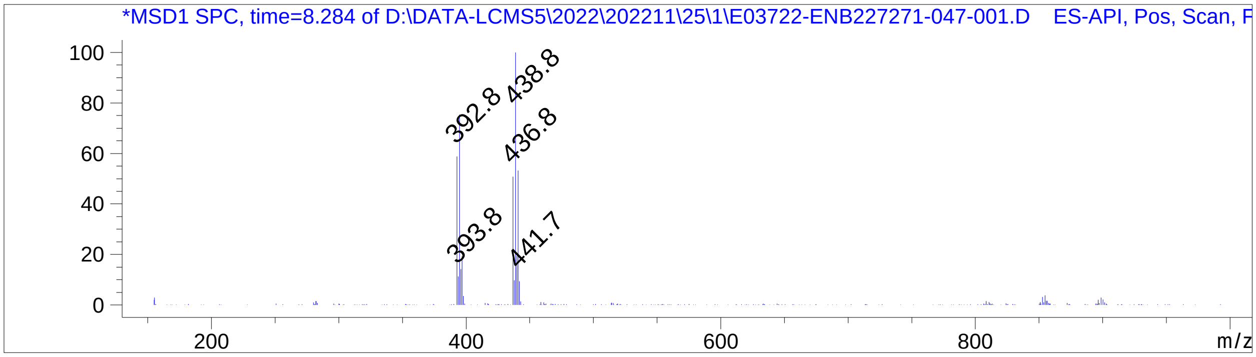


=================================================================

*** End of Report ***

### Qualitative Analysis Report

| | | | |
|---|---|---|---|
| **Data File** | 20220824 ZL1 20 ul.d | **Sample Name** | |
| **Sample Type** | Sample | **Position** | P2-F1 |
| **Instrument Name** | LC QTOF | **User Name** | |
| **Acq Method** | Small Molecule Training_0.6mLPerMin_7Min.m | **Acquired Time** | 8/24/2022 2:26:09 PM (UTC-04:00) |
| **IRM Calibration Status** | Success | **DA Method** | default.m |
| **Comment** | | | |
| **Sample Group** | | **Info.** | |
| **Stream Name** | LC 1 | **Acquisition Time (Local)** | 8/24/2022 2:26:09 PM (UTC-04:00) |
| **Acquisition SW Version** | 6200 series TOF/6500 series Q-TOF B.08.00 (B8058.0) | **QTOF Driver Version** | 8.00.00 |
| **QTOF Firmware Version** | 20.698 | **Tune Mass Range Max.** | 3200 |

#### Chromatograms

**Fragmentor Voltage** 220 **Collision Energy** 0 **Ionization Mode** ESI

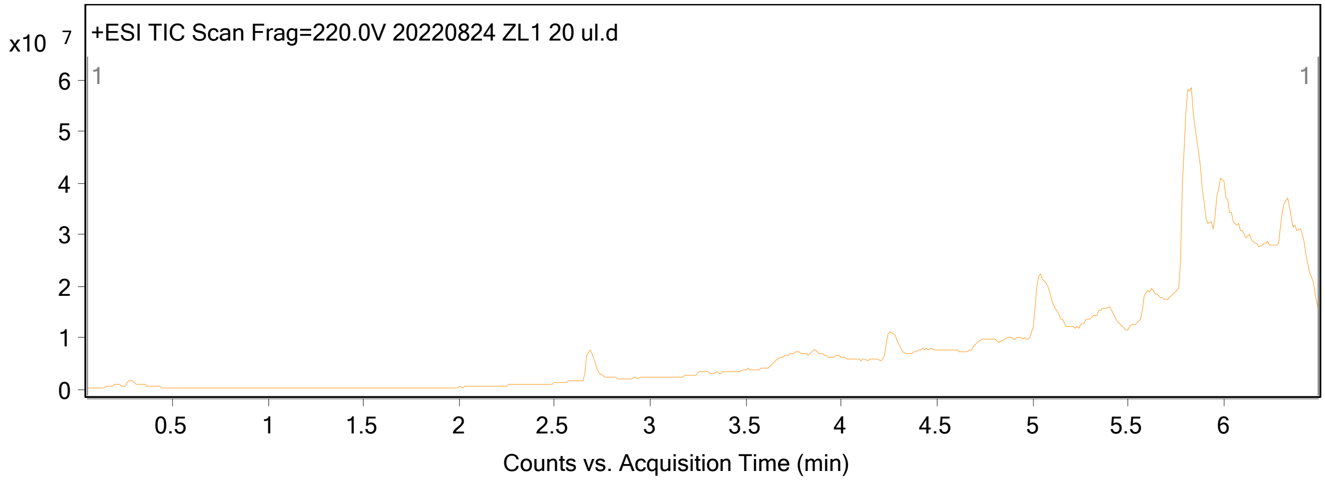

### Qualitative Analysis Report

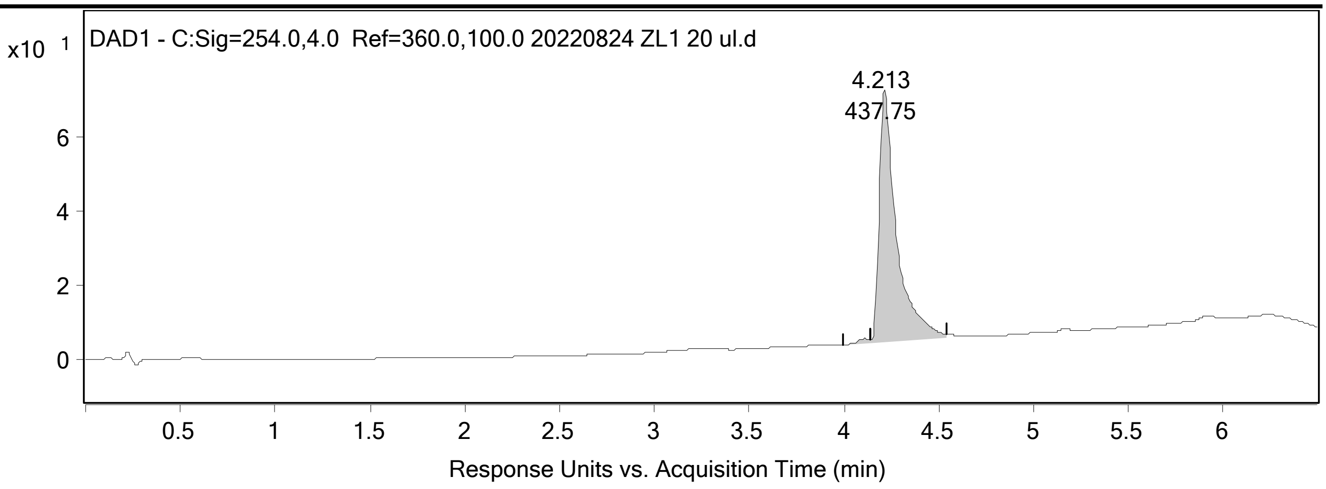


**Integration Peak List**

| Peak | Start | RT | End | Height | Area | Area % |
|---|---|---|---|---|---|---|
| 1 | 3.993 | 4.107 | 4.14 | 1.5 | 6.4 | 1.46 |
| 2 | 4.14 | 4.213 | 4.54 | 67.7 | 437.75 | 100 |

#### Spectra

| Fragmentor Voltage | Collision Energy | Ionization Mode |
|---|---|---|
| 220 | 0 | ESI |

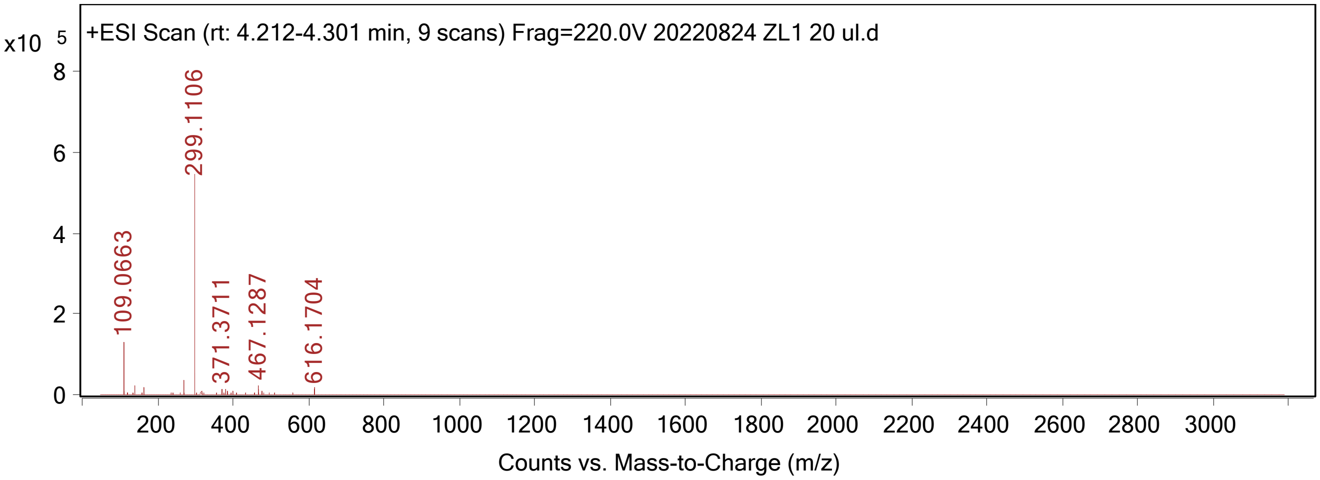


**Peak List**

| *m/z* | z | Abund |
|---|---|---|
| 109.0663 | 1 | 132829.47 |
| 137.0641 | | 23591.36 |
| 163.0658 | | 20465.17 |
| 268.2999 | | 36770.89 |
| 299.1106 | 1 | 557872.44 |
| 300.1139 | 1 | 55106.45 |
| 371.3711 | | 14502.39 |
| 382.4855 | 2 | 13468.01 |
| 467.1287 | 2 | 24927.58 |

| | | |
|---|---|---|
| 616.1704 | 2 | 16712.19 |

--- End Of Report ---

ZL2
PROTON DMSO {C:\Bruker\TopSpin3.2} core_user 2

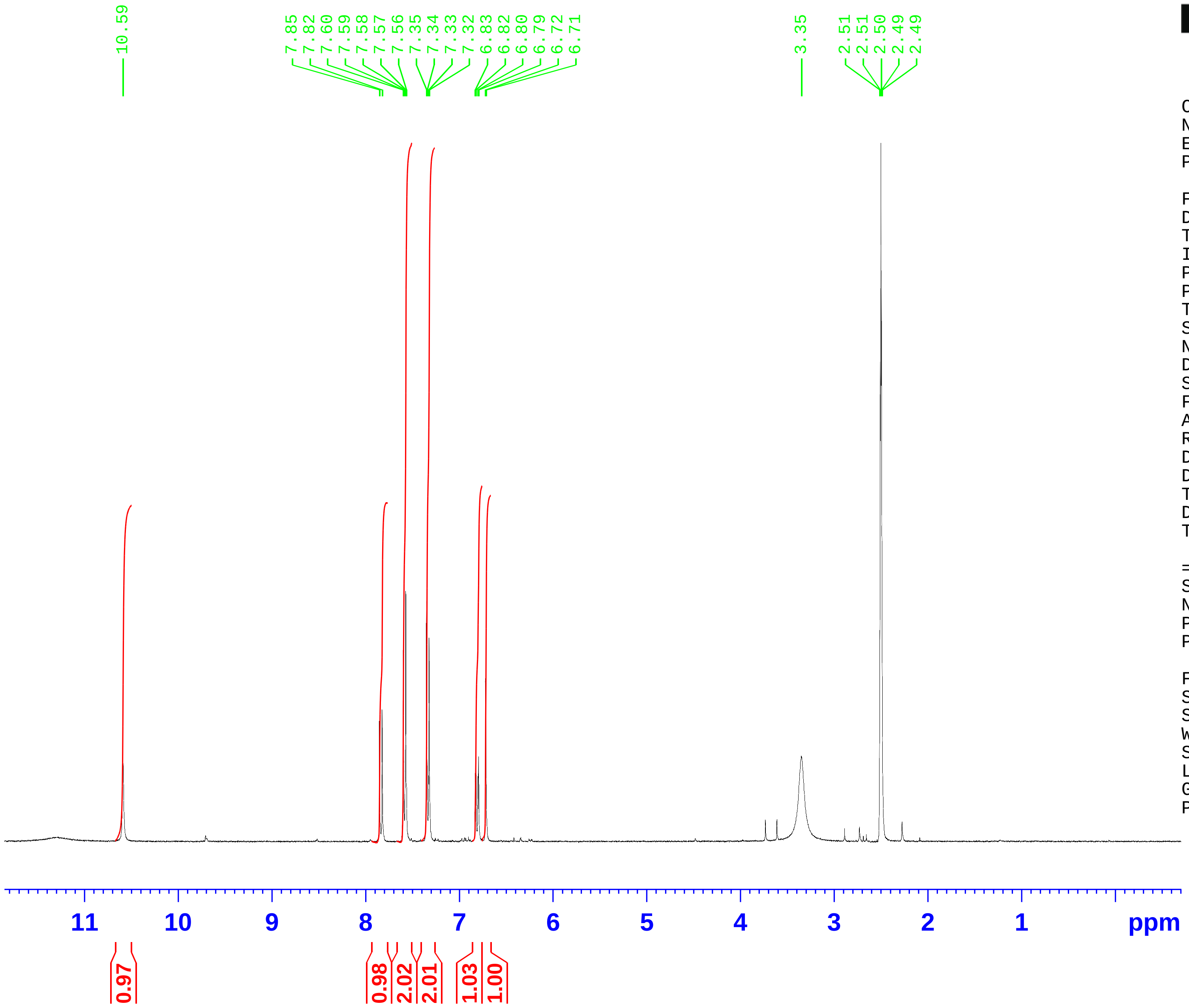


Current Data Parameters
NAME Aug19-2022
EXPNO 10
PROCNO 1

F2 - Acquisition Parameters
Date_ 20220819
Time 15.39
INSTRUM FOURIER300
PROBHD 5 mm DUL 13C-1
PULPROG zg30
TD 65536
SOLVENT DMSO
NS 16
DS 2
SWH 6103.516 Hz
FIDRES 0.093132 Hz
AQ 5.3687091 sec
RG 162.587
DW 81.920 usec
DE 6.50 usec
TE 294.3 K
D1 1.00000000 sec
TD0 1

======== CHANNEL f1 ========
SFO1 300.1818537 MHz
NUC1 1H
P1 14.20 usec
PLW1 9.00000000 W

F2 - Processing parameters
SI 65536
SF 300.1800001 MHz
WDW EM
SSB 0
LB 0.30 Hz
GB 0
PC 1.00

### Qualitative Analysis Report

| | | | |
|---|---|---|---|
| **Data File** | 20220818 ZL2.d | **Sample Name** | |
| **Sample Type** | Sample | **Position** | P2-C1 |
| **Instrument Name** | LC QTOF | **User Name** | |
| **Acq Method** | Small Molecule Training_0.6mLPerMin_7Min_30P of B.m | **Acquired Time** | 8/18/2022 4:47:51 PM (UTC-04:00) |
| **IRM Calibration Status** | Success | **DA Method** | default.m |
| **Comment** | | | |
| **Sample Group** | | **Info.** | |
| **Stream Name** | LC 1 | **Acquisition Time (Local)** | 8/18/2022 4:47:51 PM (UTC-04:00) |
| **Acquisition SW Version** | 6200 series TOF/6500 series Q-TOF B.08.00 (B8058.0) | **QTOF Driver Version** | 8.00.00 |
| **QTOF Firmware Version** | 20.698 | **Tune Mass Range Max.** | 3200 |

#### Chromatograms

**Fragmentor Voltage** 220 **Collision Energy** 0 **Ionization Mode** ESI

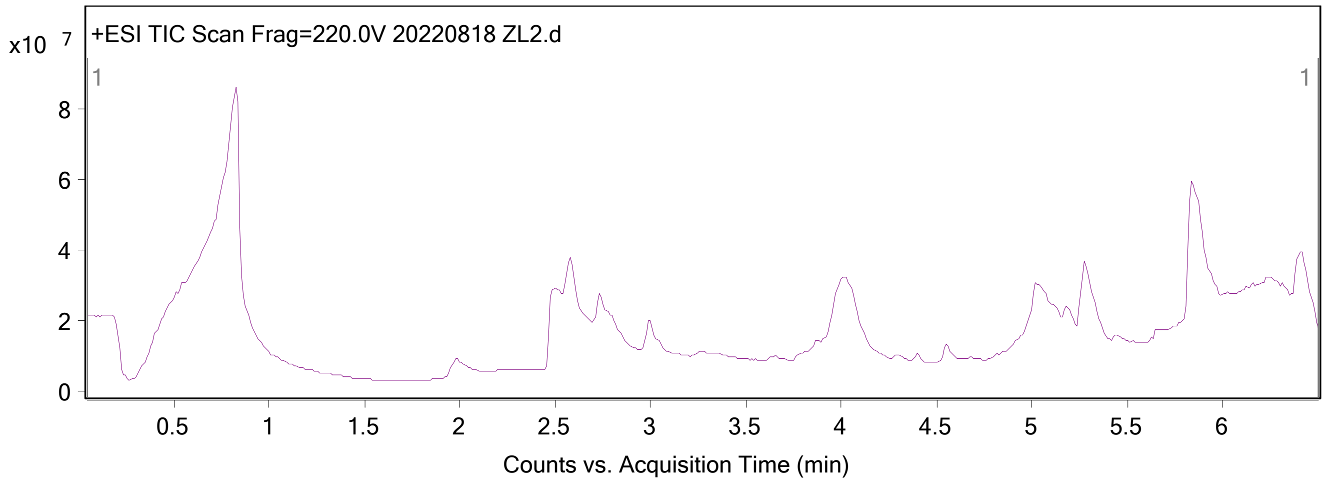

### Qualitative Analysis Report

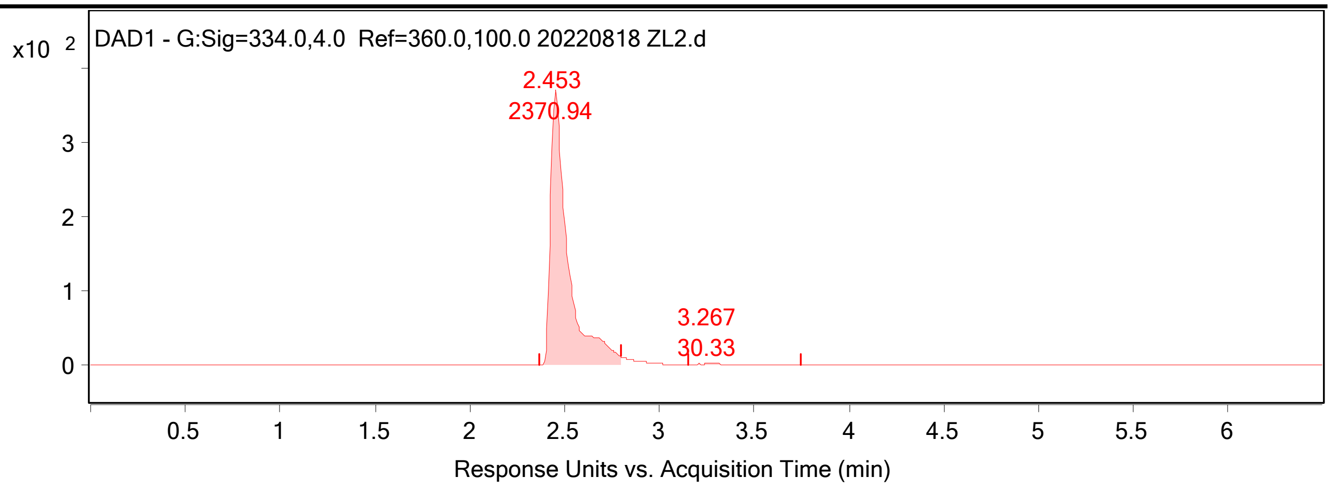


**Integration Peak List**

| Peak | Start | RT | End | Height | Area | Area % |
|---|---|---|---|---|---|---|
| 1 | 2.367 | 2.453 | 2.793 | 369.17 | 2370.94 | 100 |
| 2 | 3.153 | 3.267 | 3.747 | 2.63 | 30.33 | 1.28 |

#### Spectra

| Fragmentor Voltage | Collision Energy | Ionization Mode |
|---|---|---|
| 220 | 0 | ESI |

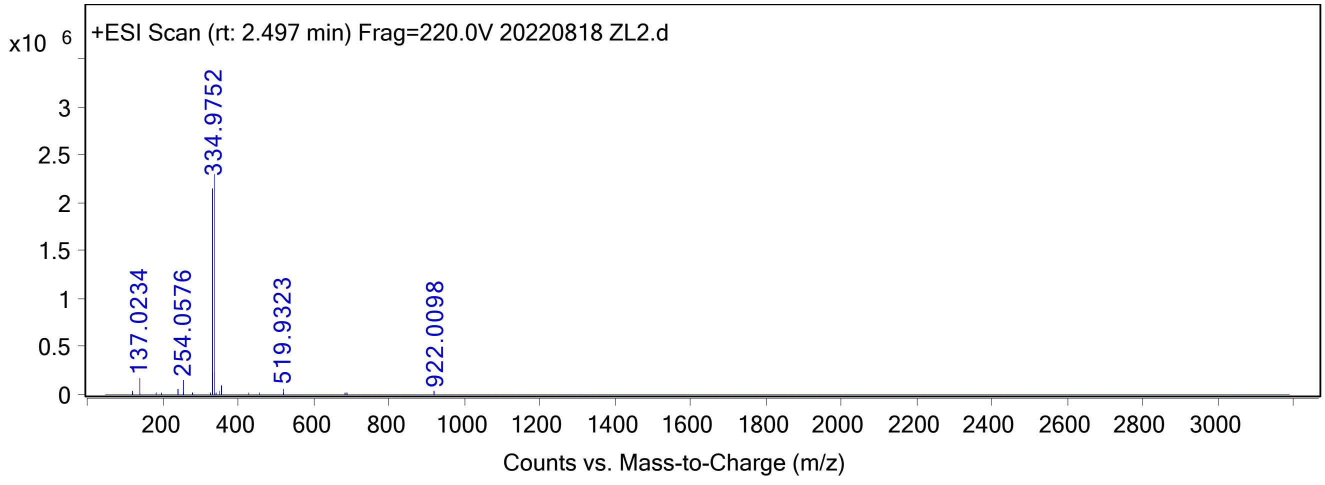


**Peak List**

| *m/z* | z | Abund |
|---|---|---|
| 137.0234 | | 182898.71 |
| 254.0576 | | 147626 |
| 332.9769 | 1 | 2210423.32 |
| 333.9798 | 1 | 230598.75 |
| 334.9752 | 1 | 2296122.76 |
| 335.978 | 1 | 235560.86 |
| 354.9578 | | 100308.95 |
| 356.956 | | 104969.98 |
| 518.9334 | 2 | 57522.26 |

| | | |
|---|---|---|
| 519.9323 | 2 | 61013.51 |

--- End Of Report ---

ZL3
PROTON DMSO {C:\Bruker\TopSpin3.2} core_user 4

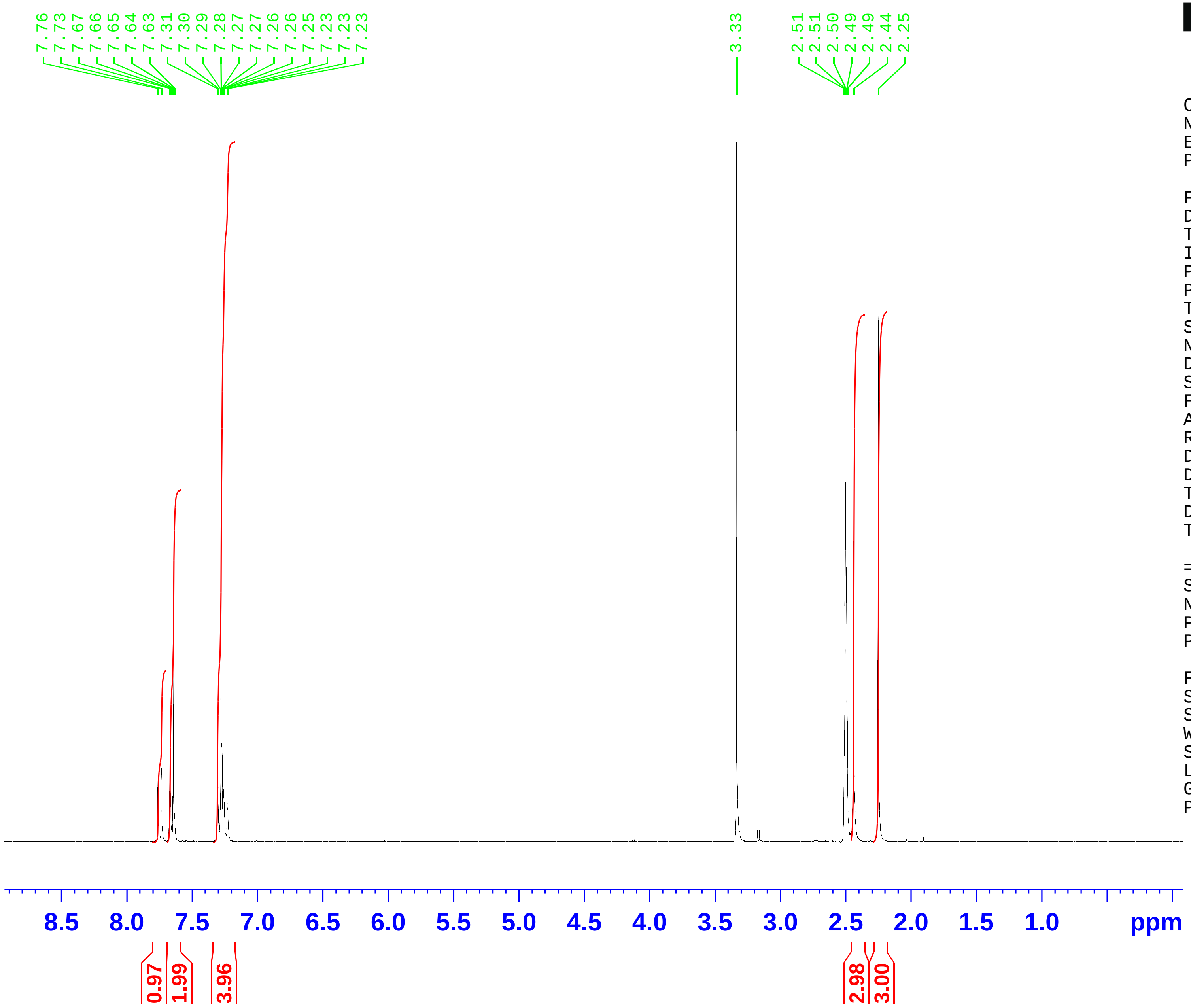


BRUKER

```
Current Data Parameters
NAME            Aug19-2022
EXPNO                   20
PROCNO                   1

F2 - Acquisition Parameters
Date_             20220819
Time                 15.45
INSTRUM         FOURIER300
PROBHD    5 mm DUL 13C-1
PULPROG               zg30
TD                   65536
SOLVENT               DMSO
NS                      16
DS                       2
SWH            6103.516 Hz
FIDRES         0.093132 Hz
AQ            5.3687091 sec
RG                 129.312
DW               81.920 usec
DE                 6.50 usec
TE                  294.4 K
D1           1.00000000 sec
TD0                      1

======== CHANNEL f1 ========
SFO1        300.1818537 MHz
NUC1                    1H
P1                14.20 usec
PLW1           9.00000000 W

F2 - Processing parameters
SI                   65536
SF          300.1800002 MHz
WDW                     EM
SSB      0
LB                 0.30 Hz
GB       0
PC                    1.00
```

### Qualitative Analysis Report

| | | | |
|---|---|---|---|
| **Data File** | 20220818 ZL3.d | **Sample Name** | |
| **Sample Type** | Sample | **Position** | P2-C2 |
| **Instrument Name** | LC QTOF | **User Name** | |
| **Acq Method** | Small Molecule Training_0.6mLPerMin_7Min_30P of B.m | **Acquired Time** | 8/18/2022 4:55:19 PM (UTC-04:00) |
| **IRM Calibration Status** | Success | **DA Method** | default.m |
| **Comment** | | | |
| **Sample Group** | | **Info.** | |
| **Stream Name** | LC 1 | **Acquisition Time (Local)** | 8/18/2022 4:55:19 PM (UTC-04:00) |
| **Acquisition SW Version** | 6200 series TOF/6500 series Q-TOF B.08.00 (B8058.0) | **QTOF Driver Version** | 8.00.00 |
| **QTOF Firmware Version** | 20.698 | **Tune Mass Range Max.** | 3200 |

#### Chromatograms

**Fragmentor Voltage** 220 **Collision Energy** 0 **Ionization Mode** ESI

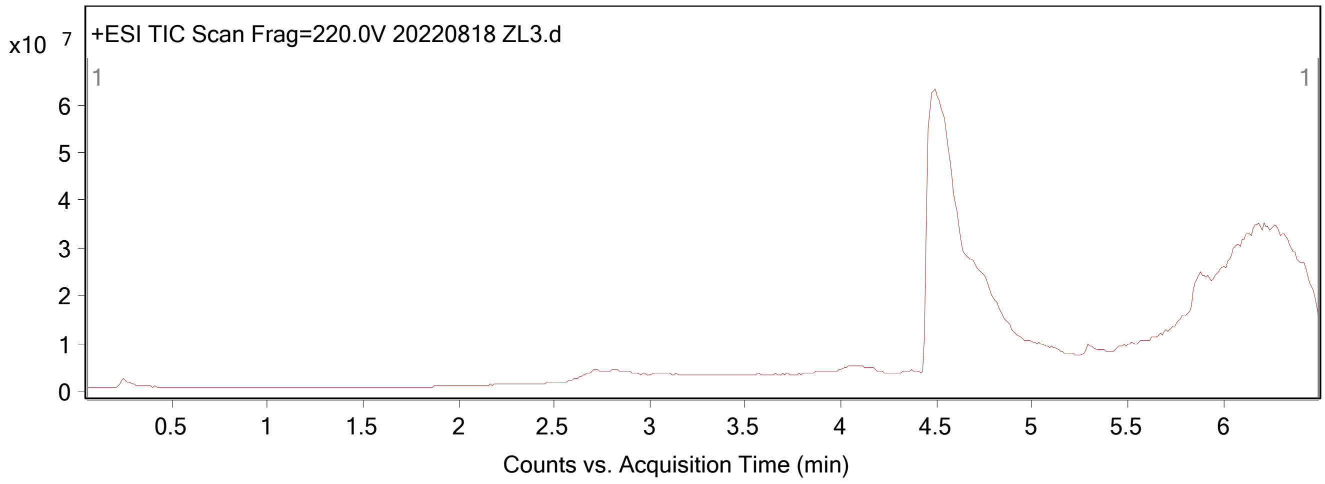

### Qualitative Analysis Report

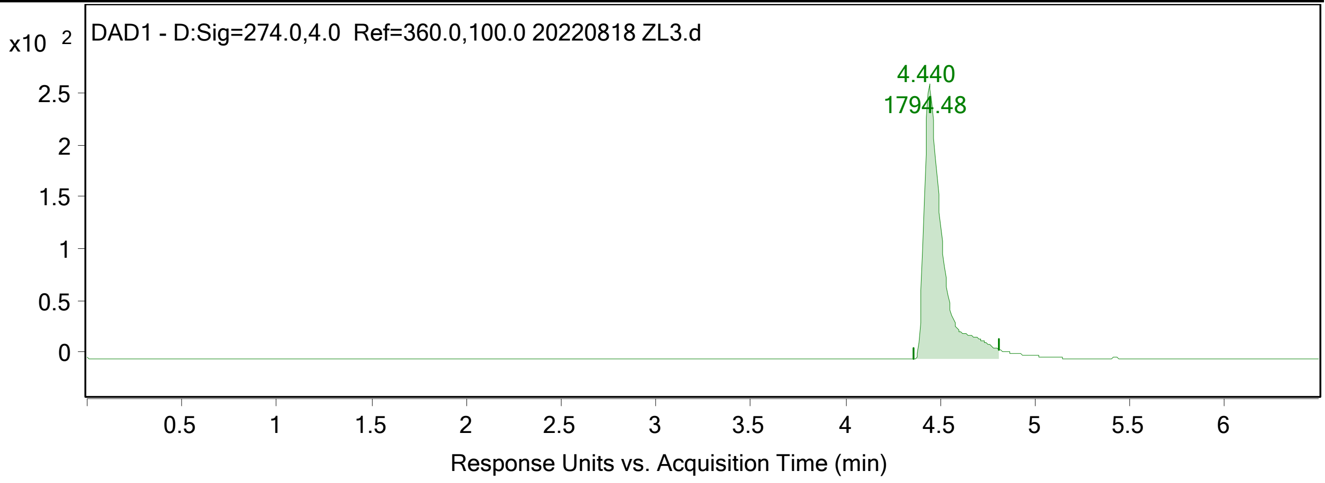


**Integration Peak List**

| Peak | Start | RT | End | Height | Area | Area % |
|---|---|---|---|---|---|---|
| 1 | 4.36 | 4.44 | 4.807 | 265.07 | 1794.48 | 100 |

#### Spectra

| Fragmentor Voltage | Collision Energy | Ionization Mode |
|---|---|---|
| 220 | 0 | ESI |

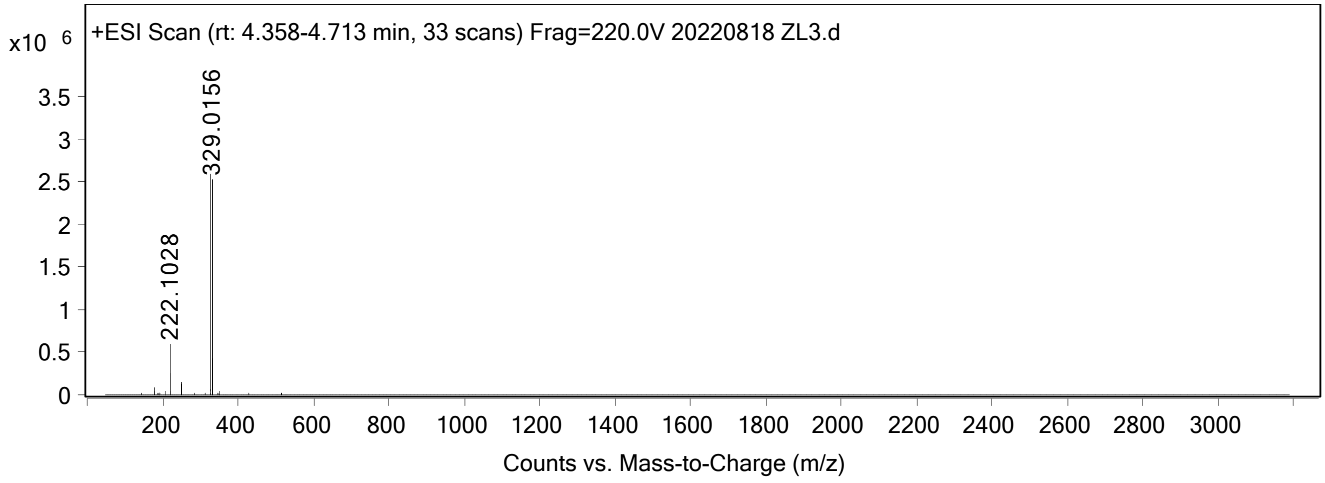


**Peak List**

| *m/z* | z | Abund |
|---|---|---|
| 178.0771 | | 98261.64 |
| 221.0956 | | 254636.19 |
| 222.1028 | 1 | 618397.51 |
| 223.106 | 1 | 52393.63 |
| 249.0898 | | 170119.28 |
| 250.0969 | | 124260.88 |
| 329.0156 | 1 | 2601066.55 |
| 330.0179 | 1 | 436239.48 |
| 331.0137 | 1 | 2603752.85 |
| 332.016 | 1 | 404439.9 |

--- End Of Report ---

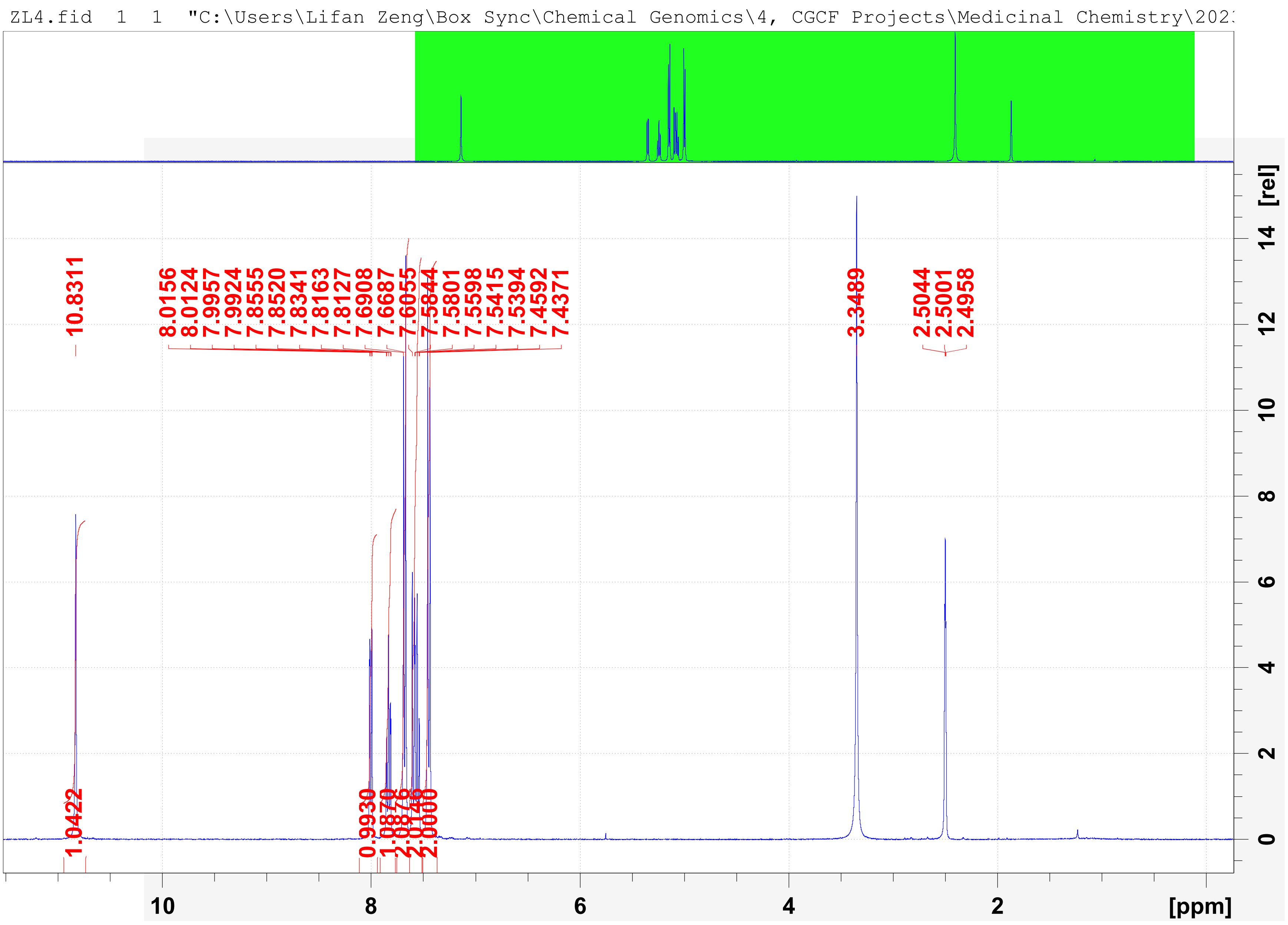
ZL4.fid 1 1 "C:\Users\Lifan Zeng\Box Sync\Chemical Genomics\4, CGCF Projects\Medicinal Chemistry\202
10.8311
8.0156
8.0124
7.9957
7.9924
7.8555
7.8520
7.8341
7.8163
7.8127
7.6908
7.6687
7.6055
7.5844
7.5801
7.5598
7.5415
7.5394
7.4592
7.4371
3.3489
2.5044
2.5001
2.4958
1.0422
0.9930
1.0870
2.0876
2.0146
2.0000
[rel]
[ppm]

### Qualitative Analysis Report

| | | | |
|---|---|---|---|
| **Data File** | 20221215 ZL4 110v.d | **Sample Name** | |
| **Sample Type** | Sample | **Position** | P2-B6 |
| **Instrument Name** | LC QTOF | **User Name** | |
| **Acq Method** | Small Molecule Training_0.6mLPerMin_7Min 110V.m | **Acquired Time** | 12/15/2022 4:10:17 PM (UTC-05:00) |
| **IRM Calibration Status** | Success | **DA Method** | default_small_molecule.m |
| **Comment** | | | |
| **Sample Group** | | **Info.** | |
| **Stream Name** | LC 1 | **Acquisition Time (Local)** | 12/15/2022 4:10:17 PM (UTC-04:00) |
| **Acquisition SW Version** | 6200 series TOF/6500 series Q-TOF B.08.00 (B8058.0) | **QTOF Driver Version** | 8.00.00 |
| **QTOF Firmware Version** | 20.698 | **Tune Mass Range Max.** | 3200 |

#### Chromatograms

**Fragmentor Voltage** 110 **Collision Energy** 0 **Ionization Mode** ESI

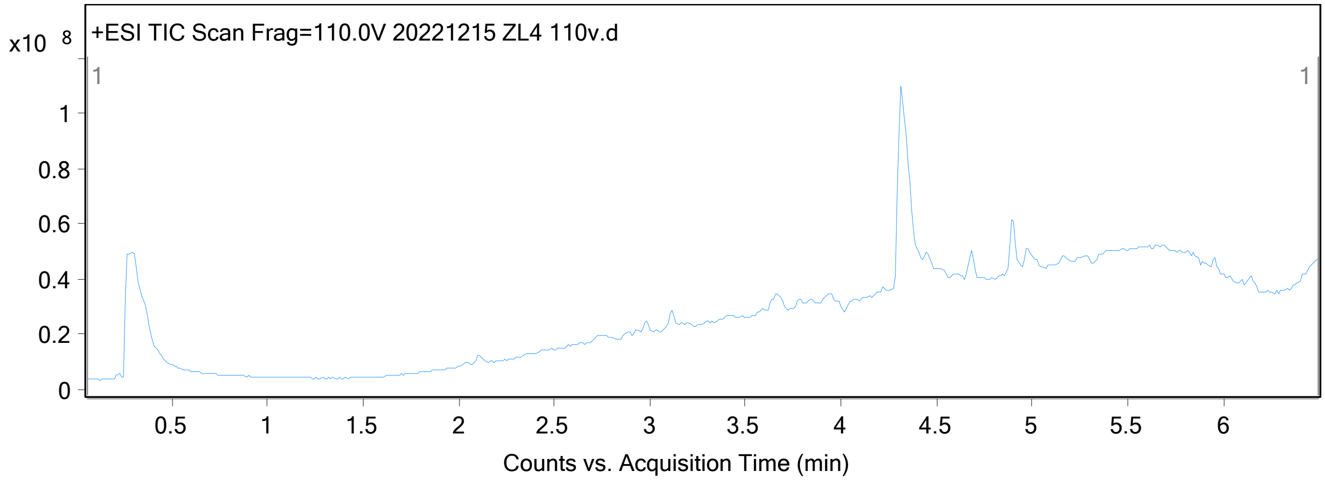

### Qualitative Analysis Report

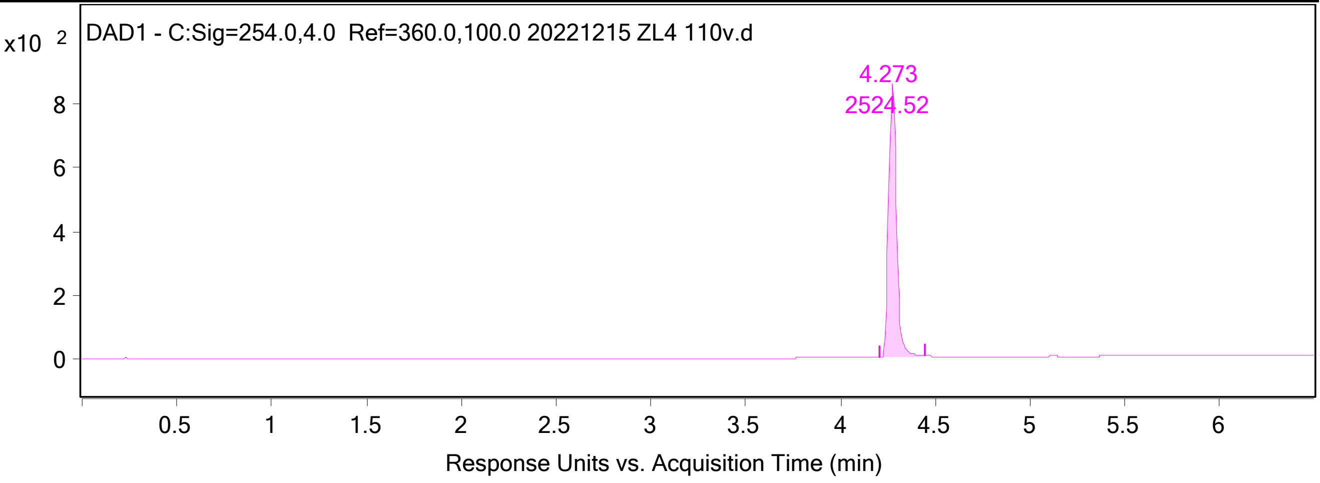


**Integration Peak List**

| Peak | Start | RT | End | Height | Area | Area % |
|---|---|---|---|---|---|---|
| 1 | 4.207 | 4.273 | 4.44 | 855.16 | 2524.52 | 100 |

#### Spectra

| Fragmentor Voltage | Collision Energy | Ionization Mode |
|---|---|---|
| 110 | 0 | ESI |

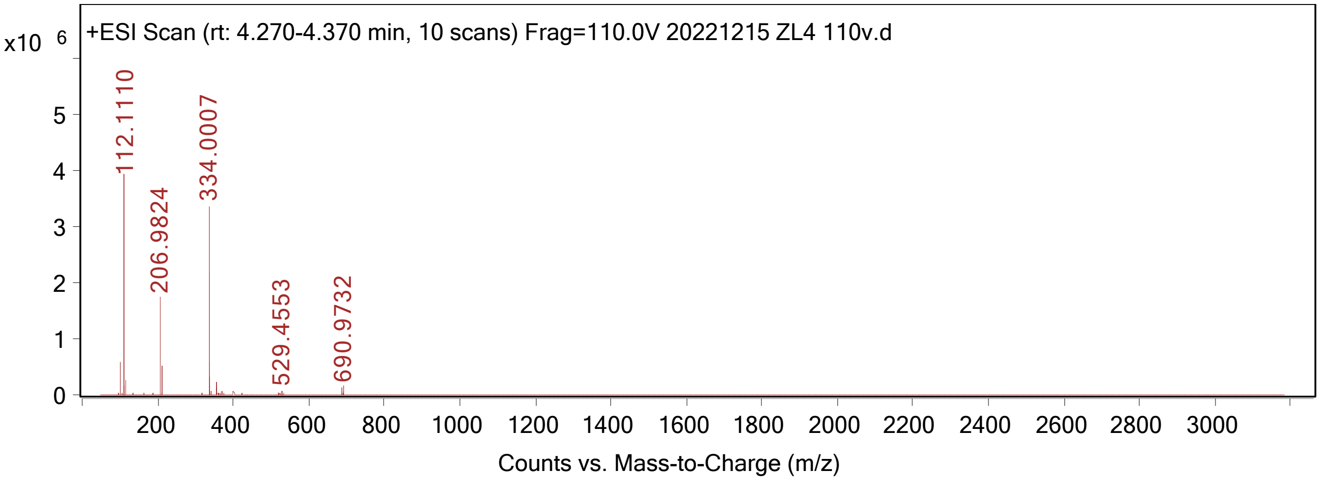


**Peak List**

| *m/z* | z | Abund |
|---|---|---|
| 102.1265 | | 594376.6 |
| 112.111 | 1 | 4062442.06 |
| 113.1142 | 1 | 276376.21 |
| 206.9824 | 1 | 1733305.16 |
| 208.9796 | 1 | 528066.91 |
| 334.0007 | 1 | 3442261.73 |
| 335.0036 | 1 | 569281.04 |
| 335.9976 | 1 | 2249844.14 |
| 337.0009 | 1 | 324610.14 |
| 337.9957 | 1 | 324381.43 |

--- End Of Report ---

### SYNTHESIS OF COMPOUNDS W1-W10

#### Synthesis of Compounds W1 (1), W2 (2) and W3 (3)

Bioduro-Sundia MediTech Company Ltd.

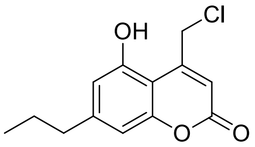


**1**

25 mg

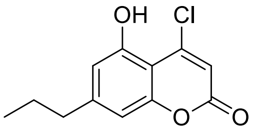


**2**

15 mg

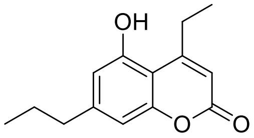


**3**

25 mg

Procedures:

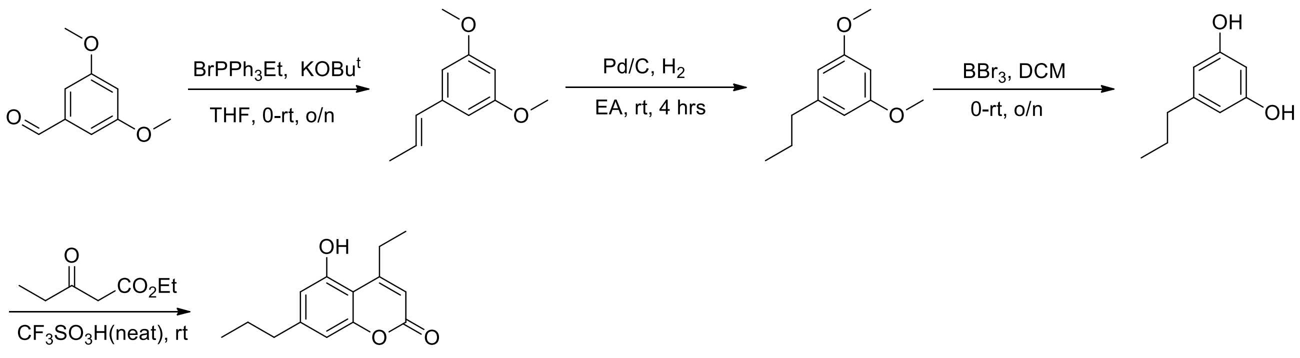


##### Step 1

To a solution of 3,5-dimethoxybenzaldehyde (1.66 g, 10.0 mmol) in THF (30 mL) was added Potassium tert-butanolate (1.68 g, 15.0 mmol) at 0$^o$C. The resulting mixture was stirred from 0$^o$C for 30 mins. Then ethyltriphenylphosphonium bromide was added and the mixture was stirred from 0$^o$C to room temperature overnight. The reaction was monitored by LC-MS and TLC. Then the mixture was filtered, and the filtrate was concentrated *in vacuum* to give a residue, which was purified by silica gel column (PE/EtOAc = 100/1 to 30/1 as eluent) to afford 1,3-dimethoxy-5-(prop-1-en-1-yl)benzene (1.45 g, yield: 81 %) as colorless oil.

##### Step 2

To a solution of 1,3-dimethoxy-5-(prop-1-en-1-yl)benzene (1.45 g, 8.10 mmol) in EtOAc (40 mL) was added Pd/C (145 mg, 10% wt). The resulting mixture was stirred at room temperature for 4 hrs. The reaction was monitored by LC-MS. Then the mixture was filtered and the filtrate was concentrated *in vacuum* to give 1,3-dimethoxy-5-propylbenzene (1.46 g, yield: 99%) as colorless oil.

##### Step 3

To a solution of 1,3-dimethoxy-5-propylbenzene (1.46 g, 8.10 mmol) in dry DCM (30 mL) was added $BB_{r3}$ (3.0 mL) at 0$^{o}$C dropwise. The resulting mixture was stirred from 0$^{o}$C to room temperature overnight. The reaction was monitored by LC-MS. Then the mixture was quenched by addition of MeOH (5.0 mL) slowly. The mixture was concentrated *in vacuum* to give a residue, which was purified by reverse-phase column (5-95% ACN in $H_2O$, 40 mins) to afford 5-propylbenzene-1,3-diol (1.18 g, yield: 96%) as yellow oil.

**Step 4**

To a mixture of 5-propylbenzene-1,3-diol (152 mg, 1.0 mmol) and ethyl 3-oxopentanoate (2.0 mL) was added $CF_3SO_2OH$ (2 drops). The resulting mixture was stirred at room temperature overnight. The reaction was monitored by LC-MS. Then MeOH (5 mL) was added and the mixture was concentrated in vacuum to give a residue, which was purified by prep-HPLC ($NH_4AC$ as additive) to afford 4-ethyl-5-hydroxy-7-propyl-2H-chromen-2-one (133.8 mg, yield: 58 %) as white solid.

$^1$HNMR (400 MHz, DMSO-*d6*): δ = 10.59 (brs, 1H), 6.66 (d, *J* = 1.2 Hz, 1H), 6.61 (d, *J* = 1.2 Hz, 1H), 6.04 (s, 1H), 2.98 (q, *J* = 7.6 Hz, 2H), 2.53 (t, *J* = 7.2 Hz, 2H), 1.65-1.53 (m, 2H), 1.19 (t, *J* = 7.6 Hz, 3H), 0.90 (t, *J* = 7.2 Hz, 3H). MS: m/z 233.1 (M+H$^+$).

Y1617-14835-017-01_HNMR(DMSO-d6, 40  Y1617-14835-017-01_LCMS.pdf

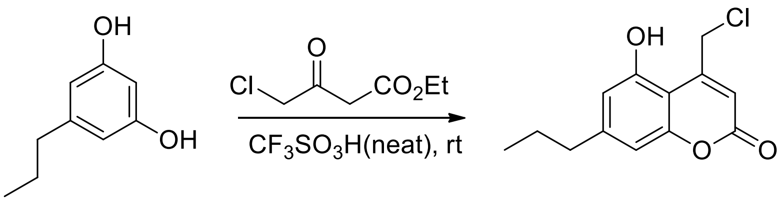


**4-(chloromethyl)-5-hydroxy-7-propyl-2H-chromen-2-one**

The title compound was prepared identical to the last step procedure of 4-ethyl-5-hydroxy-7-propyl-2H-chromen-2-one.

$^1$HNMR (400 MHz, DMSO-*d6*): δ = 10.91 (brs, 1H), 6.70 (s, 1H), 6.62 (d, *J* = 1.2 Hz, 1H), 6.42 (s, 1H), 5.09 (s, 2H), 2.54 (t, *J* = 7.2 Hz, 2H), 1.65-1.53 (m, 2H), 0.90 (t, *J* = 7.2 Hz, 3H). MS: m/z 253.1 (M+H$^+$).

Y1617-14835-018-01_HNMR(DMSO-d6, 40  Y1617-14835-018-01_LCMS.pdf

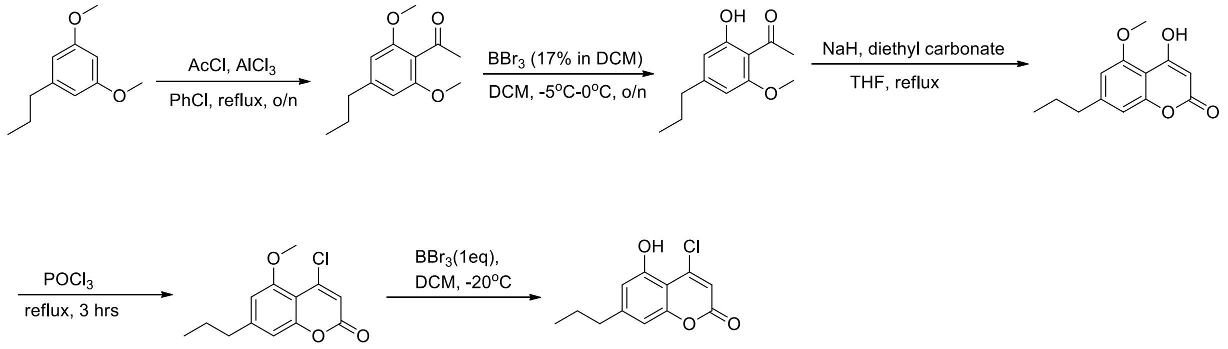


**Step 1**

To a solution of 1,3-dimethoxy-5-propylbenzene (460 mg, 2.56 mmol) in DCM (30 mL) was added acetyl chloride (300 mg, 3.83 mmol), followed by $AlCl_3$ (510 mg, 3.83 mmol) at 0°C. The resulting mixture was stirred at room temperature overnight. The reaction was monitored by LC-MS and TLC. Then the mixture was filtered and the filtrate was concentrated *in vacuum* to give a residue, which was purified by silica gel column (PE/EtOAc = 100/1 to 20/1) to afford 1-(2,6-dimethoxy-4-propylphenyl)ethanone (340 mg, yield: 60%) as yellow oil.

$^1$HNMR (400 MHz, DMSO-*d6*): δ = 6.37 (s, 2H), 3.79 (s, 6H), 2.56 (t, *J* = 8.0 Hz, 2H), 2.47 (s, 3H), 1.68-1.61 (m, 2H), 0.96 (t, *J* = 7.2 Hz, 3H).

E01617-14835-026-02_HNMR(CDCl3, 400

**Step 2**

To a solution of 1-(2,6-dimethoxy-4-propylphenyl)ethanone (340 mg, 1.53 mmol) in dry DCM (30 mL) was added BBr3 (2.0 mL, 17% in DCM) at 0°C dropwise. The resulting mixture was stirred from 0°C to room temperature overnight. The reaction was monitored by LC-MS. Then the reaction was quenched by addition of MeOH (5.0 mL) slowly. The mixture was concentrated *in vacuum* to give a residue, which was purified by silica gel column (PE/EtOAc = 100/1 to 20/1) to afford 1-(2-hydroxy-6-methoxy-4-propylphenyl)ethanone (242 mg, yield: 76%) as yellow oil.

**Step 3**

To a mixture of 1-(2-hydroxy-6-methoxy-4-propylphenyl)ethanone (192 mg, 0.92 mmol) in dry THF (20 mL) was added NaH (110 mg, 2.76 mmol). The mixture was stirred at 65°C for 10 mins and diethyl carbonate (218 mg, 1.84 mmol) was added drop wise. The resulting mixture was stirred at 65°C overnight. The reaction was monitored by LC-MS. Then MeOH (5 mL) was added to quench the reaction and the mixture was concentrated *in vacuum* to give a residue, which was purified by silica gel column (DCM to DCM/MeOH = 30/1) to afford 4-hydroxy-5-methoxy-7-propyl-2H-chromen-2-one (148 mg, yield: 58 %) as white solid.

**Step 4**

A mixture of 4-hydroxy-5-methoxy-7-propyl-2H-chromen-2-one (148 mg, 0.63 mmol) in $POCl_3$ (10 mL) was stirred at 100°C for 3 hrs. The reaction was monitored by LC-MS. Then $POCl_3$ was removed *in vacuum* to give a residue, which was purified by silica gel column (DCM) to afford 4-chloro-5-methoxy-7-propyl-2H-chromen-2-one (62 mg, yield: 39 %) as white solid.

$^1$HNMR (400 MHz, DMSO-*d6*): δ = 6.79 (s, 1H), 6.60 (s, 1H), 6.39 (s, 1H), 3.92 (s, 3H), 2.64 (t, *J* = 7.6 Hz, 2H), 1.74-1.63 (m, 2H), 0.97 (t, *J* = 7.2 Hz, 3H).

E01617-14835-030-01_HNMR(CDCl3, 400

**Step 5**

To a solution of 4-chloro-5-methoxy-7-propyl-2H-chromen-2-one (62 mg, 0.25 mmol) in dry DCM (30 mL) was added BBr3 (1.0 mL, 17% in DCM) at 0°C drop wise. The resulting mixture was stirred from 0°C to room temperature overnight. The reaction was monitored by LC-MS. Then the reaction was quenched by addition of MeOH (2.0 mL) slowly. The mixture was concentrated *in vacuum* to give a residue, which was purified by Prep-TLC (DCM/MeOH = 30/1) to afford 4-chloro-5-hydroxy-7-propyl-2H-chromen-2-one (43.4 mg, yield: 74%) as white solid.

$^1$HNMR (400 MHz, DMSO-*d6*): δ = 10.84 (brs, 1H), 6.73 (d, *J* = 1.2 Hz, 1H), 6.65 (d, *J* = 1.2 Hz, 1H), 6.48 (s, 1H), 2.54 (d, *J* = 7.2 Hz, 2H), 1.64-1.54 (m, 2H), 0.89 (t, *J* = 7.2 Hz, 3H). MS: m/z 239.0 ($M+H^+$).

E01617-14835-031-01_HNMR(DMSO-d6, 4

E01617-14835-031-01_LCMS.pdf

**Synthesis of Compound W4**

Bioduro-Sundia MediTech Company Ltd.

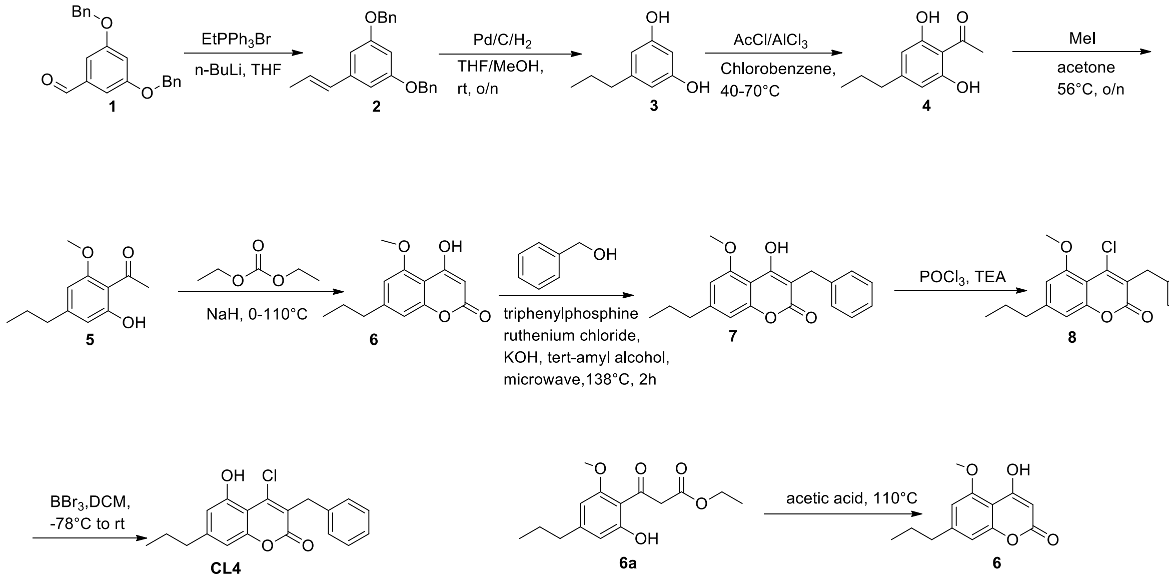

To a solution of ethyl triphenyl phosphanium bromide (32.06 g, 0.0863 mol) in THF (333 mL) was added n-butyllithium (5.53 g, 34 ml, 0.0863 mol) at 0 °C, and the reaction mixture was stirred for half an hour at 0 °C. Then 3,5-bis(benzyloxy)benzaldehyde **1** (25 g, 0.0785 mmol) was added. The reaction mixture was stirred at 25 °C for one hour. The reaction mixture was quenched with water (200 mL) and was extracted with EA (100 mL x 3). The combined organic layer was washed with water (100 mL) and brine (100 mL), dried over $Na_2SO_4$ and concentrated in vacuo. The residue was purified silica gel column chromatographed (PE/EA = 50/1) to afford compound **2** as yellow oil (13.01 g, yield: 48%). $^1$H NMR (300 MHz, $CDCl_3$): *δ* 7.58-7.37 (m, 10H), 6.70-6.57 (m, 3H), 6.50-5.80 (m, 2H), 5.12 (s, 4H), 2.00-1.92 (m, 3H). LC-MS: m/z 331.1 $[M+H]^+$.

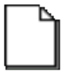

2_HNMR

To a solution of 1,3-bis(benzyloxy)-5-[(1E)-prop-1-en-1-yl]benzene **2** (13.009 g, 0.0394 mol) in THF (51 ml) and MeOH (173 ml) was added 10% Pd/C (2.6 g） at 25 °C. The mixture was stirred under hydrogen balloon for 15 hours at room temperature. The reaction solution was filtered and the filtrate was concentrated to get crude compound **3** (5.8 g, yield: 92%) as a yellow solid. $^1$H NMR (300 MHz, DMSO-$d_6$): *δ* 9.04 (s, 2H), 6.07 (s, 3H), 2.38 (t, *J* = 7.5 Hz, 2H), 1.58-1.50 (m, 2H), 0.89 (t, *J* = 7.2 Hz, 3H).

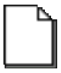

3_HNMR

To a solution of 5-propylbenzene-1,3-diol **3** (3.5 g, 0.023 mol), $AlCl_3$ (9.2 g 0.069 mol) in chlorobenzene (28 ml) was added AcCl（1.79 g 0.023 mol）at 40℃ and stirred for 30 minutes. Subsequently, the temperature was raised to 70 °C and the reaction mixture was stirred for an additional hour. The reaction was cooled to room temperature. The reaction mixture was quenched with water (50 mL) and 1M HCl (14 ml) was added until PH less than 1. The mixture was extracted with EA (50 mL x 3), the organics were washed with water (50 mL) and brine (50 mL), dried over $Na_2SO_4$. The organics were concentrated in vacuo and the residue was purified silica gel column chromatographed (PE/EA = 30/1) to afford compound **4** as a yellow solid (2.6 g, yield: 53%). LC-MS: m/z 195.0 $[M+H]^+$.

Compound 4_LCMS.pdf

A solution of 1-(2,6-dihydroxy-4-propylphenyl) ethenone **4** (1.79 g, 0.0092 mol), $K_2CO_3$ (1.27 g 0.0092 mol), $CH_3I$（1.31 g 0.0092 mol）in acetone (15.3 ml) was added into a glass tube and sealed. The reaction mixture was stirred for 22 hours at 56 ℃. The mixture was filtered. Then, the filtrate was concentrated in vacuo and the residue was purified silica gel column chromatographed (PE) to afford compound **5** as a yellow oil (1.173 g, yield: 59%). LC-MS: m/z 209.0 $[M+H]^+$.

Compound 5_LCMS.pdf

To a solution of 1-(2-hydroxy-6-methoxy-4-propylphenyl) ethanone **5** (1.335 g, 0.0064 mol) in diethyl carbonate (30 ml) was added NaH (1.54 g, 0.064 mol) at 0 ℃. Subsequently, the temperature was raised to 110 °C and the reaction mixture was stirred for 30 minutes. The reaction was cooled to room temperature. The reaction mixture was quenched with water (30 mL) and 1M HCl (12 ml) was added. The mixture was extracted with EA (50 mL x 2), the organics were washed with water (50 mL) and brine (50 mL), dried over $Na_2SO_4$. The organics were concentrated in vacuo and the residue was purified silica gel column chromatograph (PE/EA = 4/1) to afford compound **6** as a white solid (470 mg, yield: 30%). LC-MS: m/z 235.0 $[M+H]^+$.

Compound 6_LCMS.pdf

A solution of 4-hydroxy-5-methoxy-7-propylchromen-2-one **6** (200 mg, 0.8538 mmol), Benzyl alcohol (92.26 mg 0.8538 mmol), triphenylphosphine ruthenium chloride（40.94 mg 0.0426 mmol), KOH (9.59 mg, 0.01707 mmol) in tert-amyl alcohol (2 ml) was added into a microwavable glass tube. Subject the mixture to microwave irradiation at 138 °C for 2 hours. The reaction mixture was quenched with water (10 mL) and extracted with EA (10 mL x 2), the organics were washed with water (10 mL) and brine (10 mL), dried over $Na_2SO_4$. The organics were concentrated in vacuo and the residue was purified silica gel column chromatograph (PE/EA = 15/1) to afford compound **7** as a brown solid (118 mg, yield: 38%). LC-MS: m/z 325.0 $[M+H]^+$.

Compound 7_LCMS.pdf

To a solution of 3-benzyl-4-hydroxy-5-methoxy-7-propylchromen-2-one **7** (166 mg, 0.5118 mmol) in $POCl_3$ (3 ml) was added TEA (62.15 mg 0.6141 mmol). The reaction mixture was stirred for one hour at 70°C. The reaction mixture was quenched with $NaHCO_3$ (10 mL). The mixture was extracted with EA (10 mL x 3), the organics were washed with water (10 mL x 3) and brine (10 mL x 3), dried ($Na_2SO_4$). The organics were concentrated in vacuo and the residue was purified silica gel column chromatograph (PE/EA = 10/1) to afford compound **8** as a white solid (118 mg, yield: 64%). LC-MS: m/z 343.0 $[M+H]^+$.

Compound 6_LCMS.pdf

To a solution of 3-benzyl-4-chloro-5-methoxy-7-propylchromen-2-one **8** (118 mg, 0.344 mmol) in DCM (8.2 ml) was added $BBr_3$ (2M in DCM, 0.86 mL, 1.721 mmol) at -78°C. The reaction mixture was allowed to warm to room temperature and stirred for 4 hours. The reaction mixture was quenched with water (10 mL) and $NaHCO_3$ (6 mL). The mixture was extracted with EA (10 mL x 2), the organics were washed with water (10 mL) and brine (10 mL), dried over $Na_2SO_4$. The organics were concentrated in vacuo and the residue was purified by Prep-HPLC (0.1% $NH_4HCO_3$ in $H_2O$/MeCN) to afford CL4 as a white solid (50 mg, yield: 42%). $^1H$ NMR (400 MHz, $CD_3OD$): $\delta$ 7.32-7.17 (m, 5H), 6.81 (s, 1H), 6.68 (s, 1H), 3.94 (s, 2H), 2.65 (t, $J$ = 7.2 Hz, 2H), 1.71-1.65 (m, 2H), 0.96 (t, $J$ = 7.2 Hz, 3H). LC-MS: m/z 329.0 $[M+H]^+$.

CL4_HNMR CL4_LCMS

**Synthesis of Compound W5**

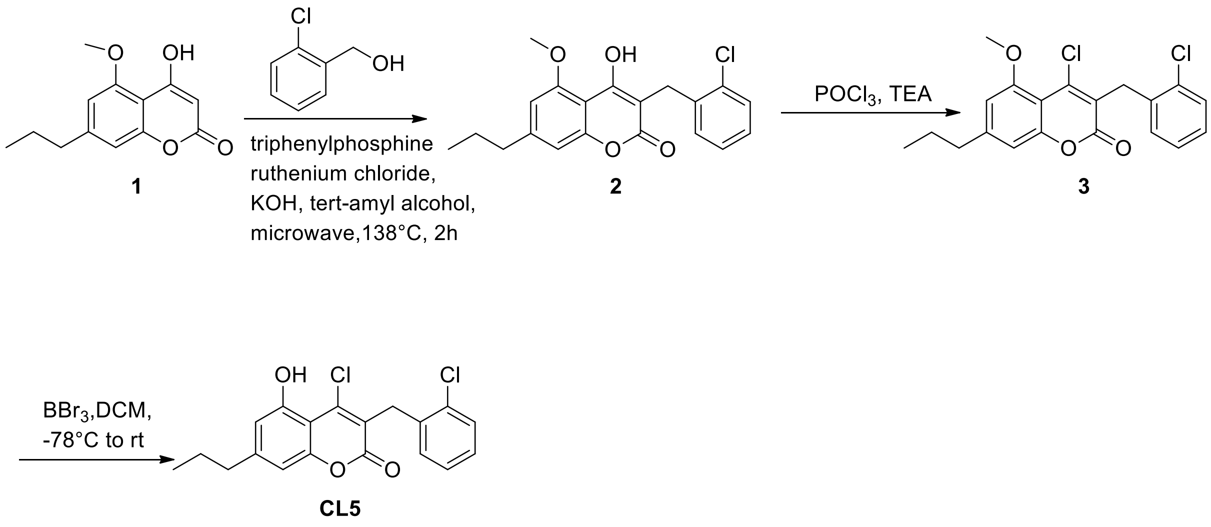


A solution of 4-hydroxy-5-methoxy-7-propylchromen-2-one **1** (620 mg, 0.265 mmol), 2-Chlorobenzyl alcohol (375 mg 0.265 mmol), triphenylphosphine ruthenium chloride（127 mg 0.0132 mmol), KOH (31 mg, 0.053 mmol) in tert-amyl alcohol (6.2 ml) was added into a microwavable glass tube. Subject the mixture to microwave irradiation at 138 °C for 2 hours. The reaction mixture was quenched with water (15 mL) and extracted with EA (10 mL x 2), the organics were washed with water (15 mL) and brine (15 mL), dried over $Na_2SO_4$. The organics were concentrated in vacuo and the residue was purified silica gel column chromatograph (PE/EA = 10/1) to afford compound **2** as a brown solid (329 mg, yield:35 %). $^1$H NMR (400 MHz, DMSO-$d_6$): $\delta$ 10.18 (s, 1H), 7.29-7.10 (m, 4H), 6.93-6.89 (m, 2H), 4.01 (s, 3H), 3.84 (s, 2H), 2.66 (t, $J$ = 7.2 Hz, 2H), 1.72-1.62 (m, 2H), 0.92 (t, $J$ = 7.2 Hz, 3H). LC-MS: m/z 359.0 $[M+H]^+$.

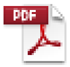

2-HNMR.pdf

To a solution of 3-[(2-chlorophenyl)methyl]-4-hydroxy-5-methoxy-7-propylchromen-2-one **2** (308 mg, 0.8584 mmol) in $POCl_3$ (5 ml) was added TEA (104 mg 1.0301 mmol). The reaction mixture was stirred for one hour at 70℃. The reaction mixture was quenched with $NaHCO_3$ (10 mL). The mixture was extracted with EA (10 mL x 3), the organics were washed with water (10 mL) and brine (10 mL) and dried over $Na_2SO_4$. The organics were concentrated in vacuo and the residue was purified silica gel column chromatograph (PE/EA = 10/1) to afford compound **3** as a white solid (178 mg, yield:52%). LC-MS: m/z 376.9 $[M+H]^+$.

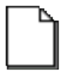

E03594-ENB2273
06-059-03.pdf

To a solution of 4-chloro-3-[(2-chlorophenyl)methyl]-5-methoxy-7-propylchromen-2-one **3** (178 mg, 0.4718 mmol) in DCM (13 ml) was added $BBr_3$ (2M in DCM, 1.2 mL, 2.359 mmol) at -78°C. The reaction mixture was allowed to warm to room temperature and stirred for 4 hours. The reaction mixture was quenched with water (10 mL) and $NaHCO_3$ (6 ml) was added. The mixture was extracted with EA (10 mL x 2), the organics were washed with water (10 mL x 3) and brine (10 mL x 3), dried ($Na_2SO_4$). The organics were concentrated in vacuo and purified by Prep-HPLC($NH_4HCO_3$) to afford CL5 as a white solid (20 mg, yield:12%). $^1$H NMR (400 MHz, $CDCl_3$): $\delta$ 12.06 (s, 1H), 7.40-7.37 (m, 1H), 7.18-7.12 (m, 2H), 7.07-7.04 (m, 1H), 6.74 (d, $J$ = 1.6 Hz, 1H), 6.69 (d, $J$ = 1.6 Hz, 1H), 4.07 (s, 2H), 2.64 (t, $J$ = 7.2 Hz, 2H), 1.71-1.63 (m, 2H), 0.96 (t, $J$ = 7.2 Hz, 3H). LC-MS: m/z 363.0 $[M+H]^+$.

CL5_HNMR CL5-HPLC.pdf CL5-LCMS.pdf

**Synthesis of Compound W6**

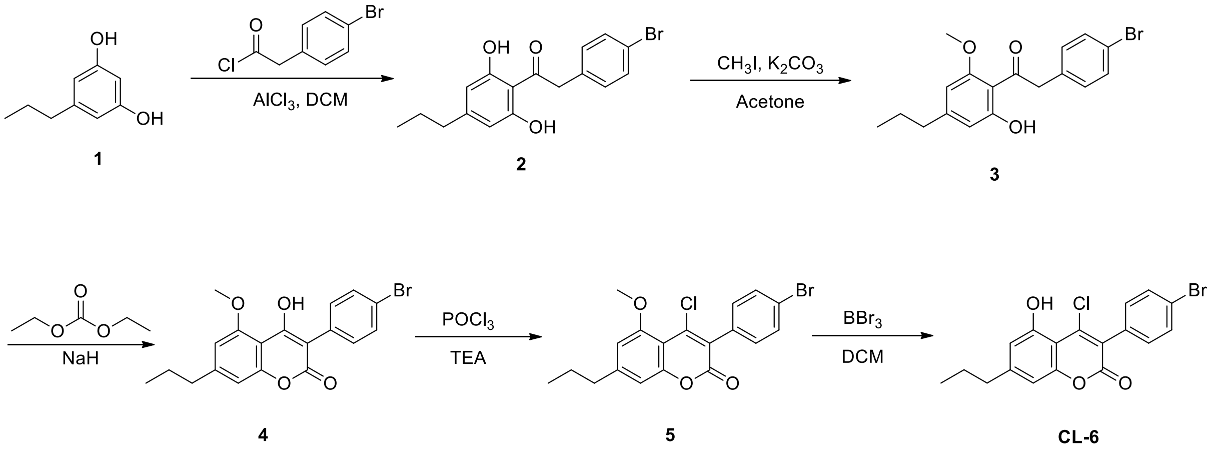


To a solution of **1** (6.75 g, 44.4 mmol), $AlCl_3$ (11.84 g, 88.8 mmol) in DCM stirred under nitrogen at 0 °C was added 2-(4-bromophenyl)acetyl chloride (10.37 g, 44.4 mmol) dropwise. The mixture was warmed to room temperature and stirred for 16 hours. The reaction mixture was quenched with 50 mL crushed ice and diluted with EtOAc. Then the mixture was washed with 1M HCl and saturated brine. The combined organic layer was dried over anhydrous $Na_2SO_4$ and concentrated in vacuum. The residue was purified by flash chromatography on silica gel (PE/EA = 8/1) to afford compound **2** (11.79 g, 68% yield) as a yellow solid. $^1$H NMR (300 MHz, DMSO-$d_6$): δ 11.78(s, 2H), 7.52 (d, $J$ = 8.1 Hz, 2H), 6.26 (s, 2H), 7.22 (d, $J$ = 8.4 Hz, 2H), 4.41 (s, 2H), 2.45 (t, $J$ = 7.2 Hz, 2H), 1.60-1.53 (m, 2H), 0.91 (t, $J$ = 7.2 Hz, 3H). LC-MS: m/z 349.0 $[M+H]^+$.

2_HNMR

To a solution of **2** (200 mg, 0.57 mmol) and $K_2CO_3$ (79.15 mg, 0.57 mmol) in acetone stirred under nitrogen at 25 °C was added $CH_3I$ (81.29 mg, 0.57 mmol) dropwise. The mixture was sealed in a tube and stirred at 58 °C for 8 hours. The reaction mixture was cooled to room temperature and filtered. The filtrate was concentrated and purified by flash chromatography on silica gel (PE/EA = 5:1) to give the product compound **3** (34 mg, 16.3% yield) as a yellow oil. $^1H$ NMR (300 MHz, $CDCl_3$) $\delta$: 13.09 (s, 1H), 7.45 (d, $J$ = 8.4 Hz, 2H), 7.08 (d, $J$ = 8.4 Hz, 2H), 6.42 (s, 1H), 6.21 (s, 1H), 4.31 (s, 2H), 3.89 (s, 3H), 2.53 (t, $J$ = 7.8 Hz, 2H), 1.69-1.60 (m, 2H), 0.95 (t, $J$ = 7.2 Hz, 3H). LC-MS: m/z 363.0 $[M+H]^+$.

3_HNMR.pdf

A solution of compound **3** (1.42 g, 4 mmol) in diethyl carbonate (10 mL) was added to a suspension of sodium hydride (60% dispersion in mineral oil, 0.94 g, 39 mmol) in diethyl carbonate (10 mL) at 0 °C. The mixture was heated to 100 °C and stirred for 3 hours. The reaction mixture was cooled to 0 °C and then quenched by dropwise addition of water until effervescence stopped. The pH of reaction mixture was adjusted to 2 with 2N HCl and extracted with EA (50 mL x 3). The combined organic phase was dried over $Na_2SO_4$, filtered and concentrated. The residue was triturated with EA to give compound **4** (1.1 g, 59% yield) as a white solid. LC-MS: m/z 388.6 $[M+H]^+$.

4_LCMS

To a solution of compound **4** (110 mg, 0.28 mmol)) in $POCl_3$ (2 mL) was added Triethylamine (43.83 mg, 0.34 mmol) at 0 °C. The mixture was warmed to 50 °C and stirred for 5 hours. The reaction mixture was quenched with aqueous $NaHCO_3$ and extracted with EA (10 mL x 3). The combined organic phase was dried over $Na_2SO_4$, filtered and concentrated. The residue was purified by flash chromatography on silica gel (PE/EA = 50/1) to give compound **5** (98 mg, 85%) as a white solid. LC-MS: m/z 306.6 $[M+H]^+$.

5_LCMS

To a solution of compound **5** (98 mg, 0.24 mmol) in DCM was added a solution of tribromoborane (1.2 mmol, 2 mol/L in DCM) dropwise under nitrogen at -70 $^oC$. The reaction mixture was stirred at -70 $^oC$ and slowly warm to room temperature, and it was continued to stir for 2 hours. Then the reaction mixture was quenched with water and extracted with EA (10 mL x 3). The combined organic phase was dried over $Na_2SO_4$, filtrated and concentrated. The residue was purified by prep-HPLC (0.1% $NH_4HCO_3$ in $H_2O$/MeCN) to give **CL-6** (30 mg, 30%) as a white solid. $^1$H NMR (400 MHz, $CD_3OD$) $\delta$: 7.63-7.60 (m, 2H), 7.32 - 7.26 (m, 2H), 6.89 (s, 1H), 6.72 (s, 1H), 2.68 (t, *J* = 7.6 Hz, 2H), 1.72-1.66 (m, 2H), 0.98 (t, *J* = 7.2 Hz, 3H). LC-MS: m/z 392.8 $[M+H]^+$.

CL6_LCMS CL6_HNMR

**Synthesis of compounds W7-W10**

Lifan Zeng, Ph. D
Chemical Genomics Core Facility,
Department of Biochemistry and Molecular Biology
School of Medicine, Indiana University
Room 1007, 635 Barnhill Dr.,
Indianapolis, IN, 46202

The LCMS analysis of the compounds was performed on Agilent 1290 LC - 6545 QTOF. Agilent ZORBAX Eclipse Plus C18 column (HD 2.1*50mm 1.8-Micron) was used for separation. 0.1% formic acid in water (v:v) as mobile phase A and 0.1% formic acid acetonitrile as mobile phase B (v:v) were used. The flow rate is 0.6 mL/min. The gradient starts as 5% B and holds for 0.5 minute, increased from 5% B to 95 %B in 4.5 minutes and holds for 0.5-minute, 95%B to 5%B in 0.5 minute and holds for 0.5 minute. NMR spectrum was collected on Bruker NMR. The chemical shifts are reported as ppm (δ) relative to the residual solvent peak.

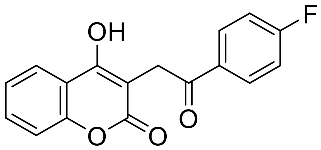


**3-(2-(4-fluorophenyl)-2-oxoethyl)-4-hydroxy-2H-chromen-2-one (W7)**

A mixture of 1-(4-fluorophenyl)-2,2-dihydroxyethan-1-one (0.1 mmol), 4-hydroxycoumarin (0.1 mmol) and 4-methoxythiophenol (0.1 mmol) in 1 mL of AcOH was heated to 130 degrees Celsius in microwave reactor for 30 min. The reaction mixture was cooled down to room temperature then poured into 15 mL of cold water. Precipitate was collected by filtration and purified by flash column chromatography using EtOAc and hexane as eluent to provide title compound as white solid. 1H NMR (300 MHz, DMSO-d6) δ 11.65 (bs, 1H), 8.19-8.16 (m, 2H), 7.96 (m, 1H), 7.64 (t, J = 10.1 Hz, 1H), 7.43-7.35 (m, 4H), 4.30 (s, 2H); Purity (98.5%, 254 nm); ESI positive mode, 299.1106 $[M+H]^+$.

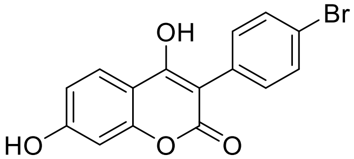


**3-(4-bromophenyl)-4,7-dihydroxy-2H-chromen-2-one (W8)**

**W8** is an in-house compound from Chemical Genomics Core. 1H NMR (300 MHz, DMSO-d6) δ 10.59 (s, 1H), 7.85 (m, 2H), 7.58 (m, 1H), 7.34 (m, 2H), 6.80 (m, 1H), 6.71 (s, 1H); Purity (98.7%, 334 nm); ESI positive mode, 334.9752 $[M+H]^+$.

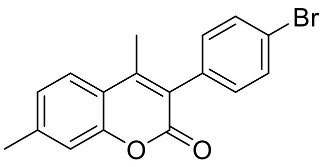

**3-(4-bromophenyl)-4,7-dimethyl-2H-chromen-2-one (W9)**

**W8** is an in-house compound from Chemical Genomics Core. 1H NMR (300 MHz, DMSO-d6) δ 7.76 (m, 1H), 7.64 (m, 2H), 7.31-7.23 (m, 4H), 2.44 (s, 3H), 2.25 (s, 3H); Purity (100%, 274 nm); ESI positive mode, 329.0156 $[M+H]^{+}$.

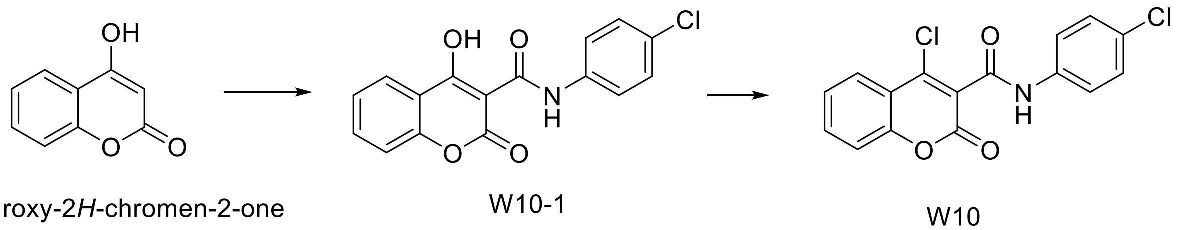


**4-chloro-N-(4-chlorophenyl)-2-oxo-2H-chromene-3-carboxamide (W10)**

20 mmol of 4-hydroxy-2H-chromen-2-one was dissolved in 30ml of dry DMSO, to which 20 mmol of pyridine and 20 mmol of 4-chlorophenyl isocyanate were added in one portion. The mixture was stirred under N2 overnight at room temperature. After adding 100 ml of 1N HCl, the mixture was stirred for 10min and the precipitate was filtered, and washed with water, EtOH, and ether. The solid was dried in vacuo overnight to give 1.14g of W10-1. 500mg of W10-1 was suspended in 20 ml of dry DCM with 3 drops of DMF. 15 mmol of oxalyl chloride was added dropwise at room temperature. The mixture was stirred overnight, and the solution became clear the next morning. The organic layer dried and evaporated to give white solid, which was purified by chromatography using DCM/Hexane to title compound as white solid. 1H NMR (300 MHz, DMSO-d6) δ 10.38 (s, 1H), 8.01 (m, 1H), 7.85 (m, 1H), 7.60 (m, 2H), 7.58 (m, 2H), 7.43 (m, 2H); Purity (100%, 254 nm); ESI positive mode, 334.0007 $[M+H]^{+}$.
